# MICRA-Net: MICRoscopy Analysis Neural Network to solve detection, classification, and segmentation from a single simple auxiliary task

**DOI:** 10.1101/2021.06.29.448970

**Authors:** Anthony Bilodeau, Constantin V.L. Delmas, Martin Parent, Paul De Koninck, Audrey Durand, Flavie Lavoie-Cardinal

## Abstract

High throughput quantitative analysis of microscopy images presents a challenge due to the complexity of the image content and the difficulty to retrieve precisely annotated datasets. In this paper we introduce a weakly-supervised MICRoscopy Analysis neural network (MICRA-Net) that can be trained on a simple main classification task using image-level annotations to solve multiple the more complex auxiliary semantic segmentation task and other associated tasks such as detection or enumeration. MICRA-Net relies on the latent information embedded within a trained model to achieve performances similar to state-of-the-art architectures when no precisely annotated dataset is available. This learnt information is extracted from the network using gradient class activation maps, which are combined to generate detailed feature maps of the biological structures of interest. We demonstrate how MICRA-Net significantly alleviates the Expert annotation process on various microscopy datasets and can be used for high-throughput quantitative analysis of microscopy images.

## 1 Introduction

The development of powerful microscopy techniques that allow to characterize biological structures with subcellular resolution and on large field of views tremendously increased the complexity of quantitative image analysis tasks [1]. The resulting images exhibit a wide range of structures that need to be identified, counted, precisely located, and segmented. Expert knowledge is commonly required to achieve successful identification and segmentation of the multiple structures of interest in microscopy images [2, 3]. These tasks can be tedious and time consuming especially for large databanks or for the comparison of multiple biological conditions. It was recently demonstrated that deep convolutional neural networks (CNN) are excellent feature extractors [4]. They were successfully applied to segmentation (e.g. whole cells, nuclei, dendritic spines), enumeration (e.g. cell counting), and classification (e.g. state of cell) of structures in microscopy images [5–12]. The most common deep learning (DL) approaches applied to microscopy and biomedical images are fully-supervised and require precisely annotated datasets [9, 11, 12]. Hence, it is often a limiting step in the application of DL for quantitative analysis of biomedical imaging [3, 13, 14]. To alleviate the annotation process, weakly-supervised DL methods were introduced [14–17]. Bounding box annotations are commonly used for weakly-supervised segmentation tasks as they are simple, allow the task to be spatially constrainted [2, 16, 18–20], and were shown to decrease the annotation phase by 15-fold compared to precise identification of structure boundaries [21]. Methods for training with binary, image-level targets, reducing even further the complexity and duration of the annotation task, have been proposed when multiple instances are displayed on a single image [22]. Unfortunately, when applied to microscopy and biomedical image analysis, such weakly-supervised approaches using whole image annotations, resulted in lower segmentation precision compared to approaches using precisely identified structures [23–25].

In this paper we propose MICRA-Net (MICRoscopy Analysis Neural Network), a new approach relying only on image-level classification annotations for training a deep neural network to perform different type of microscopy image analysis tasks such as semantic segmentation, cell counting, and detection of sparse features. MICRA-Net builds on *latent learning* [26], which refers to a model retaining information (*i.e.* latent space) that is not required for the task at hand in order to learn new auxiliary complementary tasks [26]. In this work, we leverage the information embedded within a trained classification network to solve multiple complementary, yet very different, tasks relevant to microscopy image analysis. The network uses binary classification targets as input to build a general representation of the specific dataset and generates detailed feature maps from which specific tasks, such as instance segmentation, semantic segmentation, detection, and classification, can be addressed. Even further this showcases the potential of MICRA-Net for addressing various high-throughput microscopy analysis challenges, relying solely on weak image-level annotations for training.

## 2 Results

The generation of precisely annotated large datasets to train deep neural networks in a fully-supervised manner remains a challenge in the field of microscopy and biomedical imaging. MICRA-Net, a CNN-based method, addresses this challenge by using solely whole-image binary targets for training. This approach outperforms state-of-the-art DL baselines trained in a weakly-supervised manner for the semantic segmentation of diverse biological structures. It is therefore of great interest for the automated quantitative analysis of microscopy datasets for which no fully-supervised training dataset is available. In the following we first investigate the impacts of the annotation burden, before characterizing the performance of MICRA-Net on synthetic and real data for various tasks. We then evaluate how MICRA-Net can be fine-tuned in order to leverage information from a previously acquired, but different, dataset. Finally, we show how the proposed approach could be used to support Experts in the annotation of sparse and small structures in large images.

### 2.1 Annotation task reduction analysis

MICRA-Net is trained on a simple multi-class classification task and therefore only requires the Expert to identify class-specific positive and negative images with respect to the structures of interest. In contrast to the identification of the structure boundaries using precise or bounding box contours, image-level annotations do not require to specify the positions of the object in the field of view of the microscopy images (Figure 1a).

**Figure 1:**
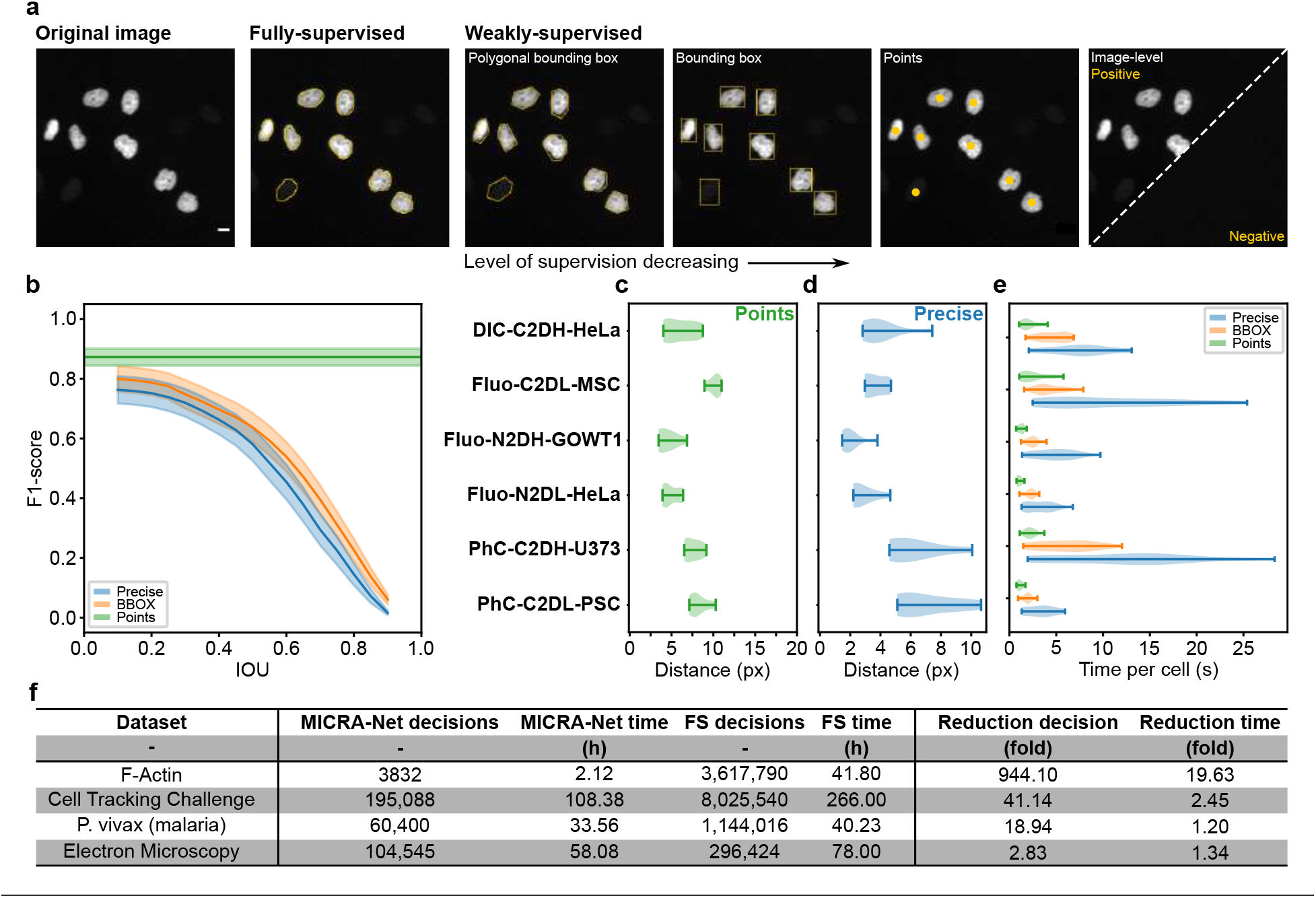
Various supervision levels can be employed for training a DL model to segment structures of interest in microscopy images. a) Representative image from the Cell Tracking Challenge dataset [8] overlayed with the corresponding fully- and weakly-supervised annotations. Annotated images are presented in decreasing spatial level of supervision and required annotation time (*from left to right*). b) We report the averaged inter-participant variability from the User-Study from 6 selected cell lines of the Cell Tracking Challenge using three levels of supervision (precise, bounding boxes (BBOX), and points). Representative examples from the participants may be found in Supplementary Fig. 1-3 as well as the specific curves per cell line in Supplementary Fig. 4. The inter-participant agreement was calculated using the F1-score as a function of IOU for precise (blue) and BBOX (orange) annotations in a all versus one manner [27]. The F1-score for points annotation (green) was calculated with a maximal distance of association of 30 pixels. Plotted are the bootstrapped mean (line) and 95% confidence interval (shade, 10 000 repetitions). c-e) Shown are the distribution of median scores from the inter-participant comparison calculated in a all versus one manner. c) Distance between associated point markers. d) Average distance between the precise contours of participants annotations was calculated for precise annotations. e) Average required time per objects on different cell lines for each supervision level. f) Evaluation of the annotation task required to generate the training set for all microscopy datasets used throughout the paper for fully-supervised (FS) and MICRA-Net approaches. Reported above is the effective number of decisions (number of extracted crops for MICRA-Net and number of edge pixels for fully-supervised learning) and the required time in hours. For MICRA-Net the number of decisions corresponds to the number of extracted crops and the annotation time per crop (assignation of a positive or negative annotation) was on average 2 seconds for all datasets. For fully-supervised learning, the decision and annotation time was evaluated for each dataset separately on a precisely annotated subset of images (see Methods).

We quantified the required time to generate annotations with different levels of precision (precise, bounding boxes, and points) by conducting a User-Study in which we asked participants to annotate the testing images from the Cell Tracking Challenge on 6 different cell lines [8] (see Methods). We analysed the inter participant variability by comparing the annotations of the participants in a one-versus-all manner. The metric used to assess this variability combines both the level of association between objects (F1-score) and the precision on the contour of annotated objects [27] (IOU, Figure 1b and Supplementary Fig. 1-3). Since it is not possible to report the IOU between points annotations, we show the average F1-score as a constant line on Figure 1b. As a general tendency, simpler annotation tasks reduced the inter-participant variability (higher F1-score at given IOU). For each selected cell lines, we report the median distances between associated point markers (centroid of objects, Figure 1c) and the average distance between the contours of associated objects (Figure 1d) as a mean to probe the variability of annotations. We measured a median error on the cell boundaries ranging from 2 to 7 pixels depending on the cell line (Figure 1d). Several factors can reduce the precision of the annotations, such as the contrast (Fluo-N2DL-HeLa - high contrast vs PhC-C2DL-PSC - low contrast) and the shape (Fluo-N2DH-GOWT1 - round vs PhC-C2DH-U373 - irregular) (Figure 1c,d and Supplementary Fig. 3). The required time to annotate a single cell is increased by approximately 2 folds when going from points annotations to bounding boxes, and from bounding boxes to precise annotations (Figure 1e). Finally, we evaluated the difference between weak-supervision using MICRA-Net’s training scheme and fully-supervised training both in terms of interactions and annotation time (Figure 1f and Methods). Compared to the precise annotations required to train fully-supervised DL approaches, the generation of whole image binary annotations reduces on average by 6 folds the required annotation duration.

### 2.2 MICRA-Net architecture and baselines

Figure 2a shows the architecture of MICRA-Net, which was designed around a CNN architecture, more specifically using a U-Net-like encoder, composed of 8 convolutional layers (*L*^1^ to *L*^8^) followed by a fully connected layer. The rationale is that U-Net is an established method able to solve multiple biomedical tasks. The gradient class activated maps (Grad-CAM, see Methods) were extracted for each predicted class and at every layer of the network (Figure 2a-c & Supplementary Fig. 5a,b). Thereafter, Rectified Linear Unit (ReLU) activation and thresholding on the Grad-CAM of the last convolutional layer (*L*^8^) were applied to generate a coarse class-specific feature map [28]. To increase the information contained in the extracted feature map, local maps from layers *L*^1–7^ were concatenated, resulting in a class-specific 7-dimensions feature space (Figure 2b,c). We retrieved the first principal component of every pixel using principal component analysis (PCA) decomposition on the feature space to generate a single feature map that was used to solve different sets of specific auxiliary tasks (Figure 2b,c & Methods).

**Figure 2:**
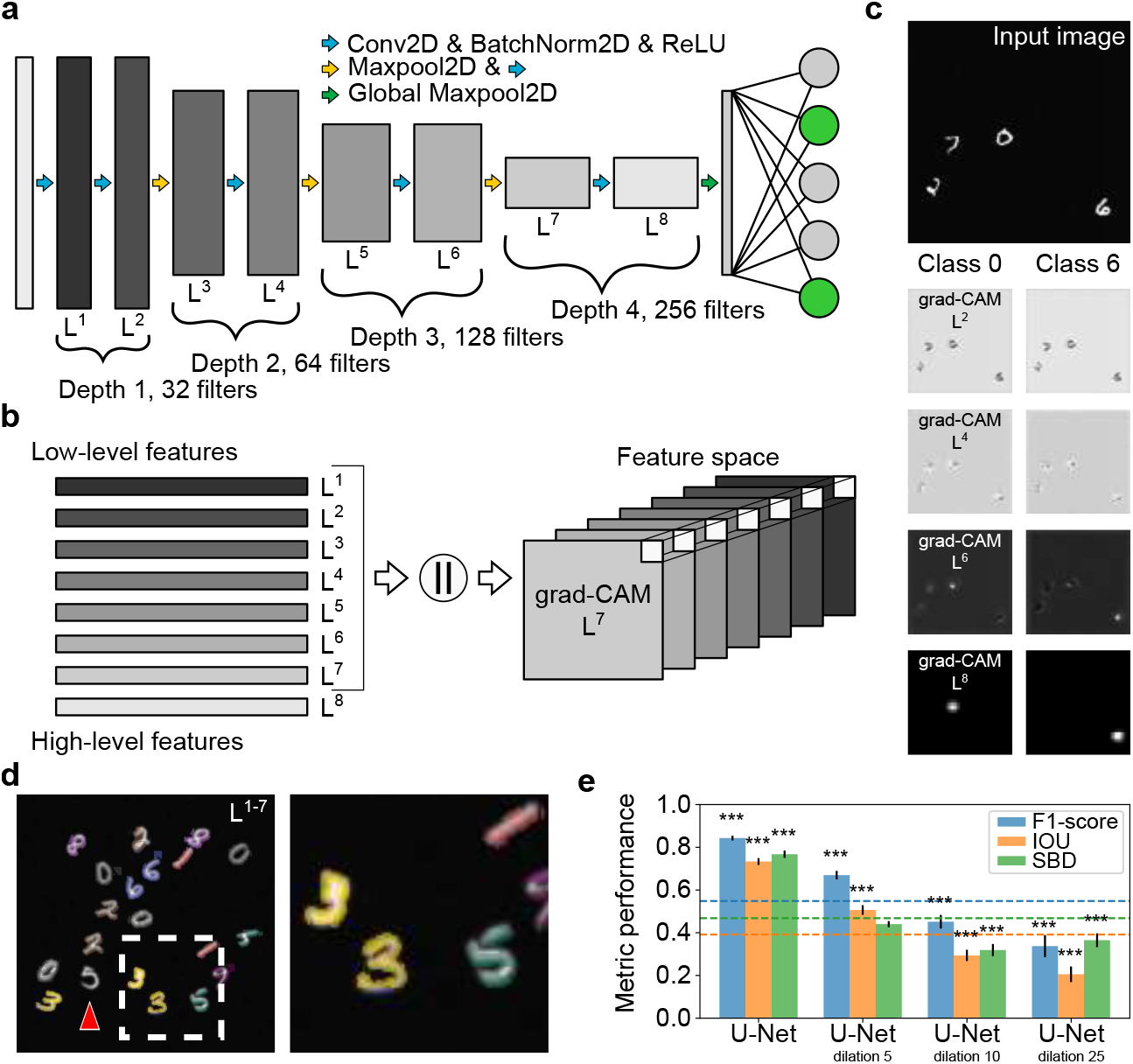
MICRA-Net architecture and experimental results on the modified MNIST dataset. a) MICRA-Net architecture (detailed in the Methods section). Each depth is composed of two sequential convolutional layers (Conv2D), batch normalization (BatchNorm2D), and Rectified Linear Unit activation (ReLU). A 2 × 2 max pooling (MaxPool2D) was employed to increase the richness of the representation from the model. A linear layer is used to project the globally pooled *L*^8^ layer (256 filters, Global Maxpool2D) to the specified number of classes. b) Concatenation of low- and high-level feature maps obtained from the Grad-CAMs of every layer is performed to generate the multi-dimensional feature space for every predicted class. c) Feature maps generated from the calculated Grad-CAMs for class 0 and 6 on the modified MNIST dataset. Each activated class is backpropagated through the network and a local map for each layer of the network (*L*^1–8^) is computed. See Supplementary Fig. 5 for layer specific grad-CAMs. d) Detailed segmentation maps of the digits of a representative image (256 × 256 pixel) and insets (right, dashed white box) from the modified MNIST dataset using MICRA-Net. The color code corresponds to the digit class and the red arrow indicates a missed digit in the field of view. e) Mean performance over the 10 classes obtained with the U-Net trained with and without dilation of the ground truth contours. The segmentation maps are presented in Supplementary Fig. 7a. MICRA-Net segmentation performance (color-coded dashed lines, see Supplementary Fig. 5 for distributions) surpasses the U-Net trained with 10 pixels dilation and is not statistically different from the U-Net trained with 5 pixels dilation on all measured metrics. Only fully supervised training outperforms MICRA-Net segmentation on all measured metrics. *p*-values are calculated using resampling (see Methods) and are reported in Supplementary Tab. 1. Bar graphs show the mean values and standard deviation.

To characterize the performance of MICRA-Net we compared the results obtained on different datasets with three established baselines: i) pretrained U-Net (in the following sections referred to as U-Net) [9], ii) Mask R-CNN [10], and iii) Ilastik [29]. These baselines were chosen as they are widely used in the literature and they allow semantic segmentation with none or simple modifications (see Supplementary Note 2 & 3 for dataset specific implementation details). This rendered a similar task between the baselines and MICRA-Net.

### 2.3 Multi-class segmentation of synthetic images

To validate the classification and segmentation performance of MICRA-Net, we created a synthetic dataset containing *N* randomly sampled cluttered handwritten digits from the MNIST dataset [30] (Modified MNIST dataset, Figure 2c & Methods). Each image may contain several instances of digits (from 0 to 9), as well as variable levels of noise and signal to mimic slight variations akin to those that may be observed in microscopy images (see Methods). The first step was to classify the digits appearing on each image to validate the representation capability of the network, which is confirmed by the obtained class-wise mean classification testing accuracy of (98.9 ± 0.5)% (mean ± std).

In addition to the classification task, MICRA-Net generates class-specific segmentation maps of the digits in the modified MNIST dataset. Using the information embedded in the Grad-CAMs of the hidden layers (*L*^1–7^) to precisely locate each digit in the image significantly increased the segmentation performance of the network when compared to the maps obtained from the Grad-CAMs of the last layer only (*L*^8^) (Figure 2d, Supplementary Fig. 5c,d & Supplementary Fig. 6). A U-Net [31] trained on the same dataset using a fully- and weakly-supervised training scheme was used as a baseline to better evaluate the performance of MICRA-Net. Fully-supervised learning consisted in training with the binary digits contours from MNIST, while weak contours were generated by a dilation of the digits with a square of size {5, 10, 25} pixel as a structuring element (see Supplementary Note 1). Figure 2e shows that MICRA-Net achieves similar or superior segmentation performance compared to all weakly-supervised training instances of the U-Net and is only outperformed on all measured metrics (F1-score, intersection over union (IOU), and symmetric boundary dice (SBD)) by fully-supervised training (Supplementary Fig. 7 & Supplementary Tab. 1).

### 2.4 Class-specific segmentation of super-resolution microscopy images

The next question that needed to be addressed was the applicability of our approach for super-resolution microscopy image segmentation, for which precisely annotated datasets are rarely available. The auxiliary task was the semantic segmentation of STimulated Emission Depletion (STED) microscopy images of two nanostructures of the F-actin cytoskeleton in neurons: 1) a periodical lattice structure (rings) and 2) longitudinal fibers (Figure 3a,b) [2]. The F-actin nanostructure segmentation task is challenging since the morphology of neurons is highly variable throughout the dataset, and there are many distractors around the structures of interest [2]. Figure 1f shows that image-level annotation reduced by more than 19 folds the time required by an Expert to generate the training dataset compared to precise identification of the structure boundaries that would be required for fully-supervised DL approaches. This also corresponds to a reduction of the annotation time of more than 3 folds compared to the tracing of polygonal bounding boxes, which were recently used for weakly-supervised training of the U-Net architecture on this dataset [2].

**Figure 3:**
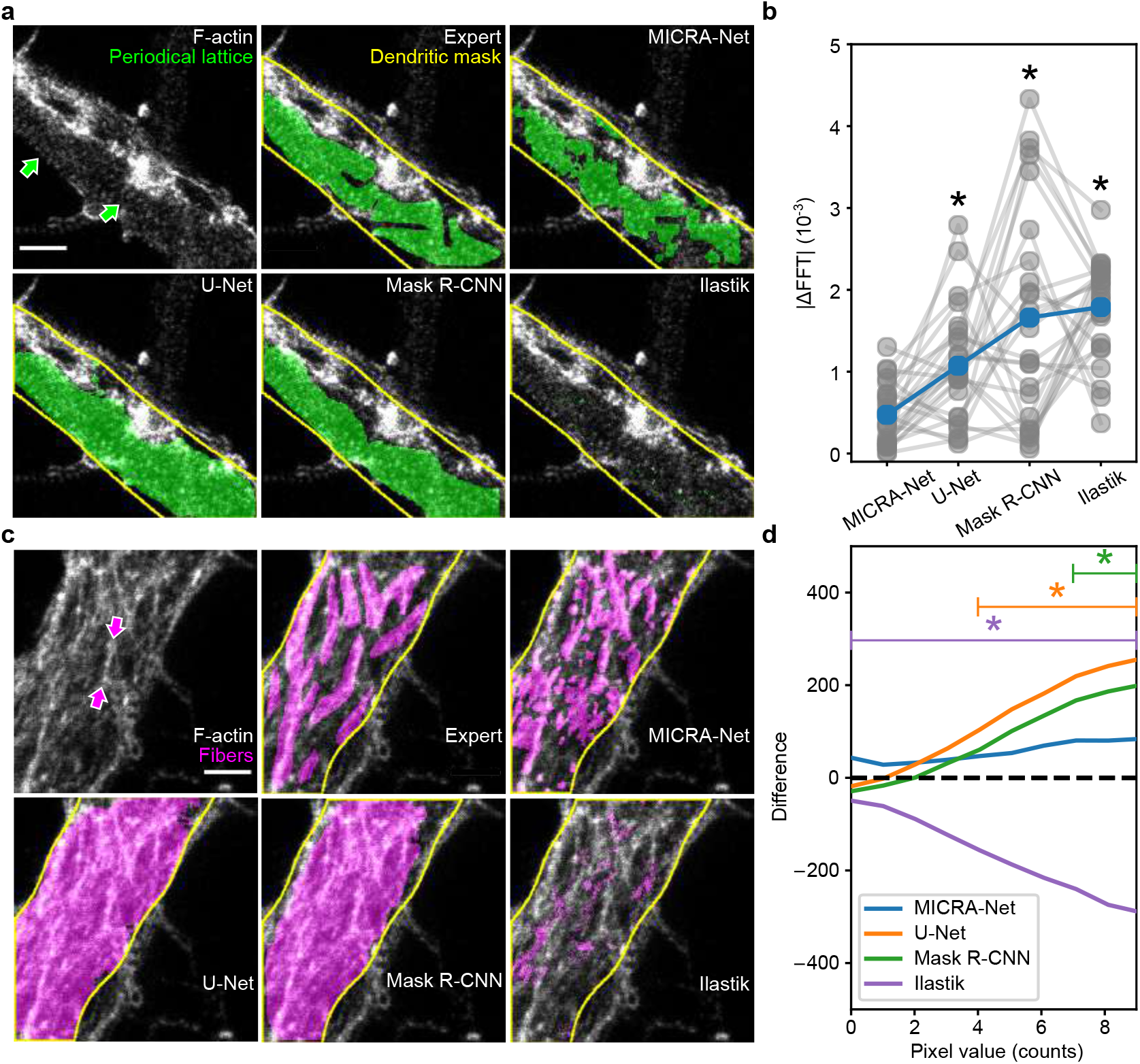
Semantic segmentation of F-actin nanostructures observed on super-resolution microscopy images. a,c) Representative raw images from a dataset of STimulated Emission Depletion (STED) microscopy images of two F-actin nanostructures in fixed cultured hippocampal neurons: periodical lattice (a) and longitudinal fibers (c). Arrows point towards the periodical lattice (green) and longitudinal fibers (magenta). Segmentation masks obtained from an Expert, MICRA-Net, weakly-supervised U-Net, weakly-supervised Mask R-CNN, and weakly-supervised Ilastik are also reported for both structures as comparison. b) Performance evaluation of MICRA-Net and weakly-supervised baselines segmentation on the precisely annotated testing dataset using custom metrics for periodical lattice (rings). The FFT metrics compares the frequency content of the provided masks. The segmentation resulting from MICRA-Net is not significantly different from the Expert annotations, while the other baselines are (U-Net, Mask R-CNN, and Ilastik). d) The intensity distribution metric evaluates the difference between the pixels found within the precise Expert annotations and the DL-based segmentation approaches for the F-actin fibers nanostructures (see Methods). The raw number of low intensity pixel segmented by MICRA-Net is not significantly different for any low value of intensity pixel from the Expert. This is not the case for all baselines (U-Net, Mask R-CNN, and Ilastik) which annotated a significantly different number of low intensity pixels. The complete range of pixel values is shown in Supplementary Fig 12. *p*-values are calculated using resampling (see Methods) and are reported in Supplementary Tab. 4, 5. Performance evaluation was performed within the dendritic mask (a,c: yellow line). a,c) Scale bars: 1 *μm*.

On the main classification task, MICRA-Net achieves an accuracy of 75.2% and 83.7% on the testing dataset for the F-actin periodical lattice and longitudinal fibers, respectively. This is inline with a mean inter-participant classification accuracy of (80 ± 5)% and (75 ± 7)% for periodical lattice and longitudinal fibers respectively (calculated from 6 participants using a leave-one-out scheme from 50 images), confirming the model capability to handle data of this nature (Supplementary Fig. 8). As described in the previous section, an informative feature map was generated from the PCA decomposition of the combined *L*^1–7^ extracted features. Thresholding of this feature map resulted in detailed binary masks that were used to solve the segmentation task. We relied on a *precisely annotated dataset* consisting of 25 images of each structure (Supplementary Fig. 9) to evaluate the performance of all trained models: i) MICRA-Net, ii) multi-participants polygonal bounding box annotations (6 participants on 25 images of each structure: *User-Study*), iii) U-Net trained with polygonal bounding boxes [2], iv) Mask R-CNN trained with polygonal bounding boxes, and v) Ilastik trained using scribbles (see Methods & Supplementary Note 2 for specific details). MICRA-Net achieved equivalent or superior segmentation performance on the *precisely annotated dataset* in comparison to both the *User-Study* and all baselines when comparing the common segmentation metrics (Supplementary Fig. 9-11 & Supplementary Tab. 2, 3). Thus, even if trained with weak image-level annotations, MICRA-Net can extract the necessary structural information to generate detailed segmentation maps for both nanostructures.

A qualitative visual inspection of the segmentation masks suggested that MICRA-Net segmentation produced a finer detailed mask compared to the weakly-supervised baseline segmentation [2], especially for fibers, for which it provides detailed segmented contours of single fiber strains (Figure 3c, Fibers). Custom performance metrics that were adapted to the F-actin nanostructures were required to better characterize this observation. For the F-actin periodical lattice, we measured the Fourier Transform (FFT) of the segmented areas for frequencies corresponding to the periodicity of the lattice (180-190 nm [32]) (Figure 3b & Methods). The FFT-metric calculated on the areas segmented with MICRA-Net is not significantly different from the one obtained from the *precisely annotated dataset* (Figure 3b). For all other baselines, evaluation of the FFT-metric on the segmented areas shows a significant difference with the *precisely annotated dataset*. This suggests a better segmentation of the periodic structure for our approach over weakly-supervised baselines (Supplementary Tab. 3, 4). Similarly, a custom metric based on the pixel intensity distribution of the segmented areas was developed to evaluate the approaches on the fiber segmentation task (see Methods). While no difference was observed for the regions identified with MICRA-Net compared to the regions from the *precisely annotated dataset*, a significant increase in the proportion of low-intensity pixels (regions between single fibers) was observed for all weakly-supervised baselines (Figure 3d & Supplementary Tab. 5). This supports a higher accuracy to precisely identify the contours of individual fibers or periodical lattice regions of MICRA-Net over weakly-supervised U-Net segmentation.

### 2.5 Single cell semantic segmentation

Cell counting and segmentation is a common challenge in high-throughput analysis of optical microscopy images [8, 9, 12, 33, 34]. Both fully- and weakly-supervised DL approaches were shown to be very powerful to assess these tasks on multiple cell lines [7, 25]. We first highlight some prerequisite of the dataset to train MICRA-Net (and baselines) at solving an instance segmentation task using 6 selected cell lines from the Cell Tracking Challenge (CTC) [8]. For weakly-supervised learning from image-level targets, a sufficient amount of negative samples (images not containing the object of interest) is required to extract informative context from an image, *i.e.* to distinguish the cells in the field of view. We trained MICRA-Net on 256 × 256 pixel crops from the resampled images of the CTC (with an effective pixel size of 0.5 μm, Supplementary Tab. 6) and obtained a classification accuracy of (95.8 ± 0.4) % (calculated from 5 network instances). Despite having a high classification accuracy, MICRA-Net detection and segmentation performances were strongly reduced when no negative samples were provided (Supplementary Fig. 13, DIC-C2DH-HeLa and Fluo-N2DH-GOWT1). It is therefore necessary to adapt the size of the training images that are provided to the network to the size of the structures of interest, ensuring that enough images contain only background (Supplementary Tab. 6 for selected factors). Another requirement when training a deep learning architecture is that the object of interest can fit entirely within the field of view. Otherwise the model has no information on how different parts of an object should be tied together. To reflect this statement, we trained both U-Net and Mask R-CNN on a resized version of the CTC dataset containing positive and negative samples on all cell lines (Supplementary Fig. 14 and Supplementary Tab. 6 for scale factors). We observe that the performance of all models is significantly lower on the DIC-C2DH-HeLa cell line at this scale. Since both training conditions cannot be met on this cell line, we removed it from training. Hence, we report the performance of all trained models on 5 selected cell lines from the CTC.

As a proof of concept, using the CTC, for which precise annotations are available, we compared the semantic instance segmentation of MICRA-Net with fully- and weakly-supervised baselines: U-Net [9], Mask R-CNN [10], and Ilastik [29] (see Supplementary Note 3 for specific implementation details). The weak supervision consisted in dilating/eroding each object of the fully-supervised dataset by a value sampled from a normal distribution with 0 mean and standard deviation in {2, 5, 10} (Altered-χ or ALT-χ), or by taking the bounding boxes of each objects (see Methods and Figure 4a). Since no precisely annotated testing dataset was provided for the CTC, we precisely annotated 4 images for each cell line to evaluate the segmentation performance of both approaches (*precisely annotated dataset*). We compared the achievable annotation precision from participants to that of altered versions of the *precisely annotated dataset* (Figure 4a,b). Figure 4b shows the distribution of IOU between associated objects (Object-IOU) of the User-Study (8 participants) and the altered versions of the dataset (8 repetitions) when comparing to the original *precisely annotated dataset* testing set for each selected cell lines of the CTC. From Figure 4b we can conclude that the distribution of the User-Study is similar to the distribution of ALT-5. Hence, training a DL architecture with a training set obtained from multiple participants (*e.g.* crowd-sourced) should result in baseline performance similar to the one trained with ALT-5.

**Figure 4:**
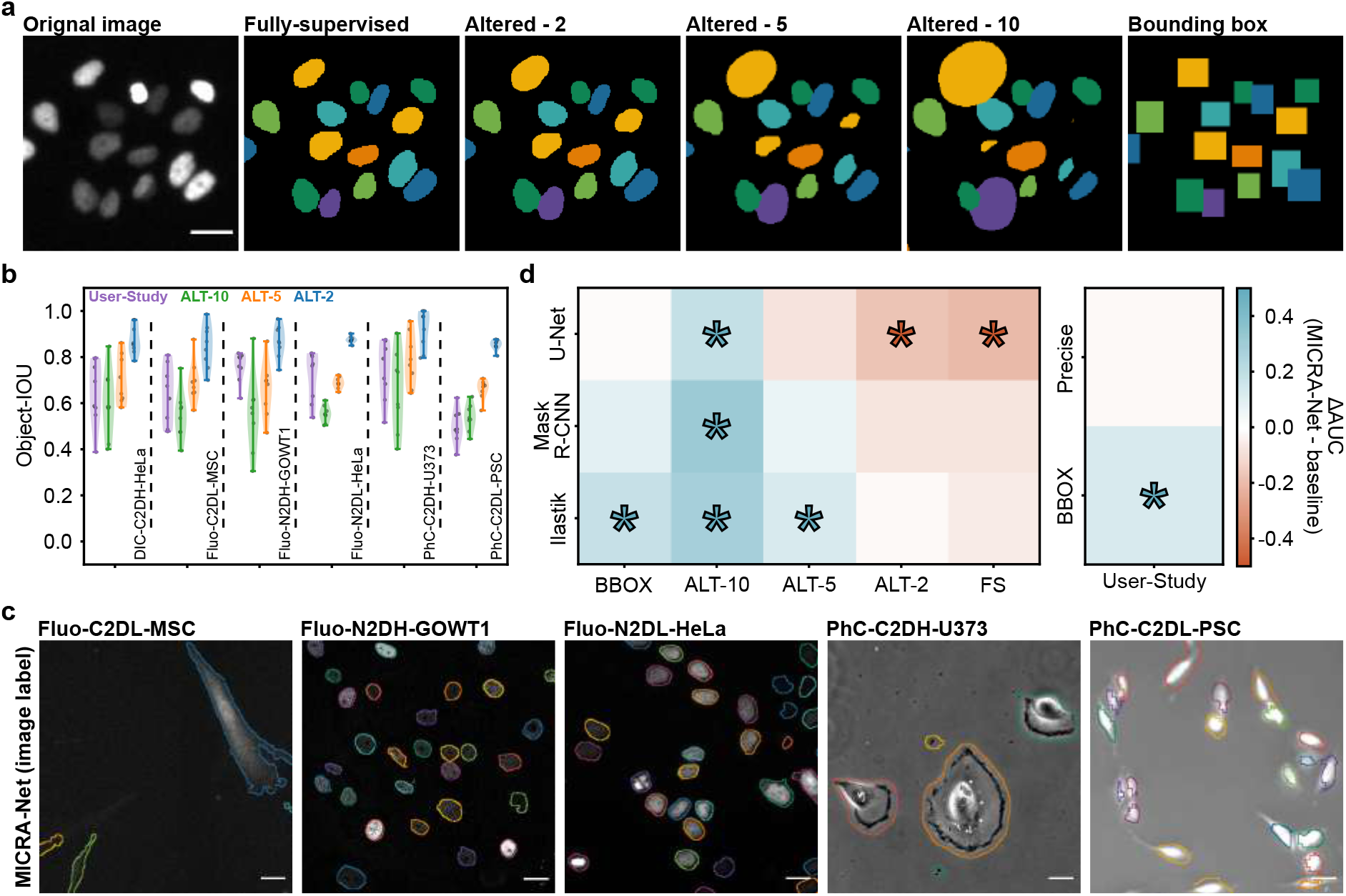
Cell counting and segmentation on 5 selected cell lines of the *Cell Tracking Challenge dataset* (CTC). a) Representative examples of various level of supervision used to train the selected baselines. The altered-χ are obtained from a binary object dilation/erosion where the transformation is sampled from a normal distribution with 0-mean and χ-standard-deviation. b) Quantification of the IOU between associated objects for the User-Study and altered versions (ALT-χ) of the testing set with the ground truth objects for each cell line of the CTC. The precision of the participants is equivalent to an ALT-5 version of the testing set. c) Representative examples of MICRA-Net semantic instance segmentation. Each outline color depicts a different segmented object. See Supplementary Fig. 16-18 for baseline examples. d, *left*) We compared the difference of the pooled area under the curve (AUC, F1-score vs. IOU) of all cell lines for MICRA-Net over the baselines on the precisely annotated dataset. The raw curves are available in Supplementary Fig. 19-23, and the non-pooled data in Supplementary Fig. 24. Higher and lower performance of MICRA-Net are reported in blue and red respectively. MICRA-Net is only outperformed by U-Net trained using ALT-2 or fully-supervised training. d, *right*) We compared the pooled AUC for all cell lines for the conducted User-Study using precise annotations and bounding boxes. The precision of the segmentation masks generated with MICRA-Net is similar to the precise annotations and better than the bounding boxes obtained in the User-Study. Stars are used to highlight a significant change (Supplementary Tab. 9, 10). All scale bars are 25 μm.

To solve the semantic instance segmentation task for MICRA-Net, we trained MICRA-Net to predict both the presence of a cell and the contact between cells, which was subtracted from the former (see Methods & Supplementary Fig. 15). A binary segmentation map was obtained by using an Otsu threshold [35] (Figure 4c). We used the F1-score detection as a function of the intersection over union (IOU) between associated objects (see Methods) to quantify the results [27]. We extracted a single score from the curves by calculating the normalized area under the curve (AUC). Figure 4d (left) reports the variation of MICRA-Net in AUC from baselines trained with various level of supervision when pooling data from all selected cell lines (see Methods and Supplementary Fig. 16-24). As shown in Figure 4d and Supplementary Fig 24, the performance of baselines which were developed for fully-supervised datasets is affected when reducing the supervision level (Supplementary Fig. 16-18). This is also depicted by the low classification accuracy of the baselines compared to MICRA-Net (Supplementary Tab. 7). Strikingly, MICRA-Net achieves cumulative similar performance to fully-supervised Mask R-CNN and Ilastik for the semantic instance segmentation on the 5 cell lines. MICRA-Net is only significantly outperformed by U-Net when training is performed on the fully-supervised or a slightly altered (ALT-2) dataset. Therefore, when no precisely annotated and proofed dataset is available, or when the annotation error may be high, the performance of baseline architectures cannot be guaranteed to achieve superior semantic instance segmentation performance on all cell lines (see Supplementary Fig. 24, and Supplementary Tab. 8 and 9). The performance of the conducted User-Study on the testing dataset were also compared to MICRA-Net (Figure 4d (right), Supplementary Fig. 2, 3). A significant increase in performance is measured for MICRA-Net for bounding boxes and no significant change is observed when comparing to precise annotations. Given the previous results, an approach like MICRA-Net will perform similarly (or better) to the presented baselines for semantic instance segmentation when no precisely annotated dataset is available. More importantly, MICRA-Net reduced by a factor of 40 the number of Expert decisions required to annotate the training dataset and by more than 150h the necessary annotation time usually needed to complete this task while achieving precise human-level precision (Figure 1f and Figure 4c).

### 2.6 Multi-device analysis

While DL approaches can be very powerful when tackling tasks on very similar images, challenges are often encountered when the imaging conditions change over time (e.g. due to a new device) [37, 38]. To increase the applicability of the proposed method to various experimental conditions, we investigated how MICRA-Net could be fine-tuned on a new dataset that contains similar structures but acquired on a new device. To address this, a brightfield microscopy dataset of Giemsa-stained [39] P. Vivax (malaria) infected human blood smears was used (Figure 5a), for which the training and testing datasets had very distinct intensity distributions (Figure 5a,b) [33, 36].

**Figure 5:**
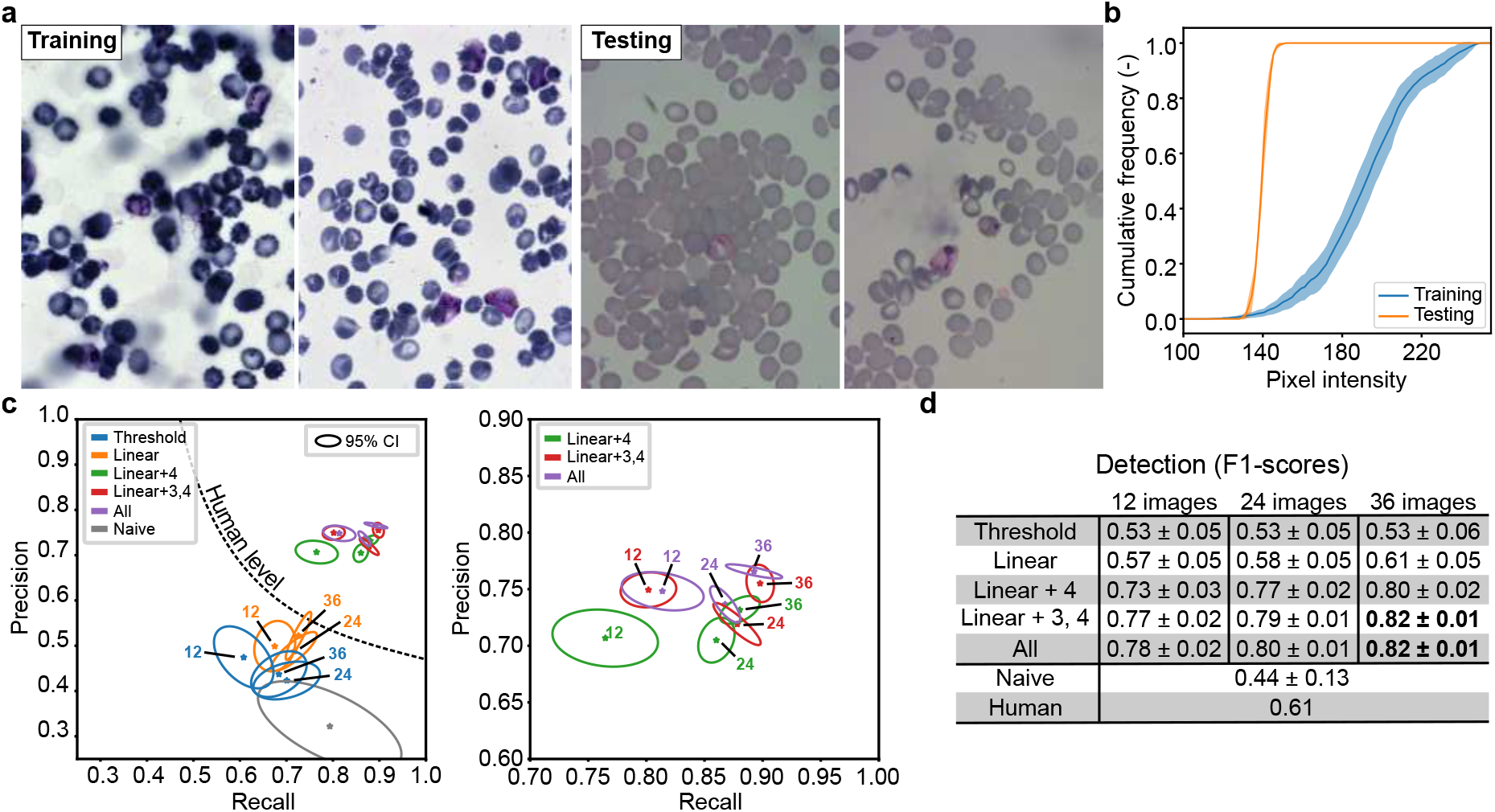
Segmentation of two different datasets of bright field microscopy images of Giemsa-stained red blood cells from [36]. a) Representative images from the training (2 left) and testing (2 right) datasets. The training dataset is composed of images taken from two different laboratories, while the testing images were acquired in a third laboratory. b) A change in the brightness and contrast is observed between the training and testing dataset. This results in a large difference in the mean pixel intensities (training: blue line, testing: orange line, with standard deviation: pale region) of the training and testing images. c, left) A precision-recall graph quantifies the detection performance of MICRA-Net on the testing dataset. Without fine-tuning, the performance on the testing dataset (Naive, grey ellipse) is characterized by a recall of 0.79, and a poor precision of 0.32. A variable number of images ({12, 24, 36}) from the testing dataset were used to adjust the detection threshold (Threshold, blue ellipse), which increased the precision but also reduced the recall by approximately 2 folds. Fine-tuning of the model on the sampled {12, 24, 36} images from the testing set with different settings: i) allowing the linear layer (orange), and ii) different depths (depth 4: green; depth 3, 4: red) to be updated (see Supplementary Fig. 25 & Supplementary Note 4) resulted in precision-recall above human level detection. c, right) Zoomed region of the precision-recall performance of MICRA-Net. When the number of trainable parameters increases, the number of images required for a model with good generalization properties also increases. d) Detection efficiency (F1-score) of the various trained fine-tuned models. As a general tendency, increasing the number of images sampled from the testing set and allowing more layers to be updated resulted in better detection of infected red blood cells. The best detection accuracy of all trained models is highlighted in bold. See Supplementary Tab. 17 for calculated *p*-values.

The first attempt to solve the classification task consisted in predicting the presence of infected smears in a 256 × 256 pixel image. A mean testing classification accuracy of (80 ± 10)% (mean ± standard deviation, calculated from 5 different instances of the network) was obtained. Since the testing images had a very different pixel intensity distribution, we investigated whether the classification results could be improved by adjusting for this. To this aim, we considered i) modifying the threshold of the linear layer and ii) fine-tuning a model by training on {12, 24,} 36 sampled images from the test set using a *k*-fold training scheme (see Supplementary Note 4 & Supplementary Fig. 25). We repeated the fine-tuning process 5 times from each of the 5 naive instantiations (as starting points) while allowing i) linear layer [*Linear*], ii) linear layer and depth 4 [*Linear + 4*], iii) linear layer and depths 3 and 4 [*Linear + 3, 4*], and iv) all [*All*] layers to be updated (Figures 2a & 5c). A testing classification accuracy over 87% was obtained when updating the threshold and over 88% for all fine-tuned models, demonstrating the capability of MICRA-Net to be fine-tuned on similar tasks performed on images acquired on different devices (Supplementary Table. 16 for detailed classification results).

In the context of parasite detection and stage determination for malaria, the most important task consists in the detection of infected cells [33]. When trained solely on the original training set, MICRA-Net performed worse on the detection task, obtaining a F1-score of 0.44 ± 0.13 (Figure 5c, d). However, with fine-tuning of at least the linear layer and the depth 4 of the architecture, the F1-score was significantly increased, beating the inter-expert accordance (0.61 [36]). Additionally, increasing the number of images sampled from the testing set can significantly increase the detection accuracy (Supplementary Tab. 17). The best detection accuracy (0.82 ± 0.01) was obtained by updating either *Linear + 3, 4* or *All* layers. This again demonstrates the capability of MICRA-Net to be fine-tuned and used across different microscopes.

We compared the segmentation results of MICRA-Net with Expert precise annotations. Due to the lack of a precisely annotated dataset in the original publication by [33], we manually segmented all infected smears from the test set (303 smears). In contrast to the results obtained for the detection accuracy, updating more layers while fine-tuning (*Linear + 3, 4* {12, 24, 36}, and *All* {12, 24}) significantly reduced the IOU compared to only updating the linear layer (Supplementary Fig. 26 & Supplementary Table 18). Hence, a trade-off should be made by the users according to their specific needs. For instance, with these P. Vivax datasets, the best trade-off to maximize both detection and segmentation efficiency requires the fine-tuning of at least the linear layer and depth 4.

### 2.7 Expert detection and segmentation assistance

The next step was to assess how MICRA-Net could be implemented as a tool to guide Experts in the annotation of sparse and small structures in large images of an electron microscopy dataset. Our approach was tested on a dataset of Scanning Electron Microscopy (SEM) images of ultrathin mouse brain sections in which axons were genetically labeled with a small engineered peroxidase APEX2 [40] (refered to as Axon DAB, see Methods). In the SEM dataset, 1-10 small axonal regions (with an averaged size of 113 × 113 pixel) needed to be identified in images of around 10 000 × 10 000 pixel (Figure 6a). Applied to this dataset, MICRA-Net was used to suggest regions containing the Axon DAB marker and generate segmentation masks of the structure in the regions that were accepted by the Expert.

**Figure 6:**
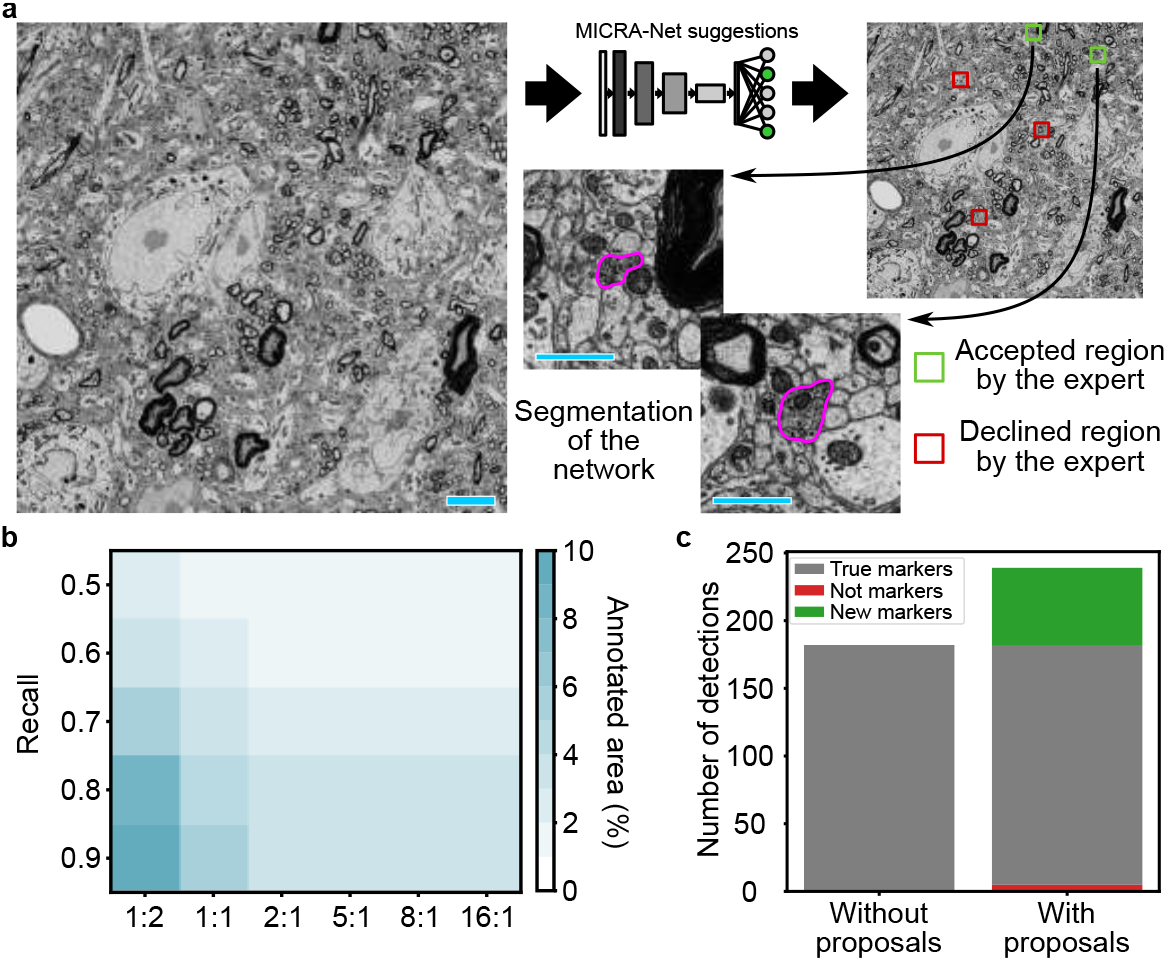
MICRA-Net is used as a tool to assist Experts in the detection of sparse Axon DAB markers in large SEM images of ultrathin mouse brain sections. a) Schematic representation of the proposed approach. MICRA-Net is first swept over the entire field of view with a 75% overlap in both directions to output the probability of presence of an axonal DAB markers. The probability of overlapping crops are then averaged to generate a probabilistic map of positions. The plausible regions are then viewed by the Expert who can accept or decline it. For each accepted region, MICRA-Net generates a segmentation map of the Axon DAB. b) The total percentage of annotated area is color-coded as a function of the positive-unlabeled (PU) ratio at the inter-expert for different recall. Using MICRA-Net trained with a PU ratio of 1:5 as an assisting tool results in the validation of approximately 3% of an image which would require an Expert less than 15 minutes to validate the complete testing set (44 images) and result in a recall of 0.9. The annotated area as a function of the recall for each PU ratio is shown in Supplementary Fig. 27. c) Total number of detections from the testing dataset with and without assistance from MICRA-Net. Using MICRA-Net the Expert could identify 57 new Axon DAB positive regions which correspond to an increase of 25% in the total number of detections. The scale bar is 5 μm for the full field of view and and is 1 μm for extracted crops.

An Expert identified Axon DAB positive regions on the training (158 images) and testing (44 images) sets using point annotations (see Methods). To train MICRA-Net, all positive regions (1024 × 1024 pixel i.e. 5.12 × 5.12 μm^2^) centered on the detected Axon DAB were extracted from the original images (image size of 10 240 × 10 240). As previously stated, MICRA-Net requires negative crops (not containing Axon DAB) for training. Therefore, all negative 1024 × 1024 pixel crops without overlap (Figure 6a, Methods & Supplementary Note 5) were also included in the dataset.

In the context of very sparse detections, positive-unlabeled (PU) learning can improve the performance of a given architecture [41]. On the main classification task, an accuracy between 83% and 90% was obtained for all PU ratios (Supplementary Tab. 19). We next investigated how PU learning could improve the detection rate of Axon DAB in the SEM images and obtained best performances for a PU ratio between 1:5 and 1:16 (Figure 6b & Supplementary Tab. 20). The usage of MICRA-Net for this sparse detection task resulted in an increase of the measured recall above the inter-expert accordance (0.791, Supplementary Fig. 27), while requiring from an Expert to proof only 3.13% of a newly acquired image. Accordingly, the area that was inspected by the Expert and consequently the annotation time were reduced by 30 folds. Additionally, MICRA-Net allowed the Expert to detect 57 new Axon DAB regions in the test set (representing 25% more detections) that had been missed by the Expert during the initial image annotation process (Figure 6c). This demonstrates the potential of MICRA-Net as a tool to assist Experts in the analysis of newly acquired images, not only reducing the manual annotation time, but also increasing the recall above the inter-expert variability. An attempt was made at comparing the detection results with Ilastik as a baseline trained on positive pixels obtained from points annotations with constant size. Ilastik achieved a classification accuracy of 8% resulting in an almost complete annotation of a new image (Supplementary Fig. 28). We also inspected how MICRA-Net performed on a second auxiliary task: the segmentation of Axon DAB regions (Supplementary Fig. 29a). For this purpose, an Expert carefully highlighted the boundaries of 170 positive Axon DAB regions sampled from the testing set. As in the detection task, MICRA-Net had the same tendency of achieving better performance with PU ratios above 1:2 and could obtain a maximal IOU score of 0.62 ± 0.03 with the 1:5 ratio (Supplementary Fig. 29 & Supplementary Tab. 21). Application of MICRA-Net to this electron microscopy annotation task was thus successful to reduce the burden of generating the training dataset, while also significantly increasing the discovery of regions of interest that were missed by the manual Expert annotation.

## 3 Discussion

While pixel-wise metrics and ground-truth annotations are well established in the field of DL and computer vision with natural images, retrieval of ground truth annotations in biomedical imaging is a laborious process, requires highly-trained Experts, and annotation imprecision often occurs [3, 42] (Figure 1). This stresses the need for weakly-supervised DL approaches that do not rely on spatially precise annotations of the structure of interest, but rather on annotations that are easier and faster to retrieve. MICRA-Net, a CNN-based method, relies on the information embedded in the latent space of a main simple task, in our case classification, to learn multiple complementary tasks without the need to generate task-specific precisely annotated training sets. We designed multiple experiments to challenge MICRA-Net at solving common microscopy tasks (segmentation, enumeration, or localization) relevant to high-throughput microscopy image analysis [3, 9]. Unlike multi-task learning [43], MICRA-Net does not combine auxiliary tasks to increase the learning performance of a main task, nor requires more annotations from the dataset for each task [44, 45]. Hence, the use of MICRA-Net should significantly reduce the burden of task-specific annotation of bioimaging datasets thereby increasing the accessibility of such deep learning based microscopy image analysis.

Our results show that MICRA-Net can be applied to various microscopy modalities and biological contexts, while significantly reducing the number of required Expert decisions to generate the training dataset (Figure 1b). While fully-supervised DL approaches (e.g. based on U-Net or Mask R-CNN architectures) have the drawback of being costly to train, they can benefit from pre-training [9, 46, 47] given the image space is similar [48], and have access to precise information about the structure boundaries. On the other hand, MICRA-Net leverages on the extraction of spatial features from the hidden layers of the network to generate detailed feature maps using solely, easy to retrieve, binary image-level annotations for training. Considering the observed reduction of the inter-expert variability when diminishing the complexity of the annotations, this will be an important aspect for future applications leveraging on crowd-sourced annotations for training. MICRA-Net provides similar or even superior performance on multiple tasks to the state-of-the art weakly- and fully-supervised learning approaches, thus making it an unprecedented alternative to address bioimaging analysis challenges for which large and precisely annotated datasets are not available.

Additionally we demonstrated that MICRA-Net could be fine-tuned when facing strong variations in the quality of the available datasets, for example when images were acquired on two different microscopes. Fine-tuning of the architecture on few images from another microscopy system was sufficient to achieve better detection efficiency than inter-expert agreement. This is of particular interest for large-scale studies, conducted on multiple sites, that require analysis framework to be easily adaptable to new experimental conditions [33, 49, 50]. Future work on fine-tuning of such approaches to new structures of interest and analysis task will be an important step to increase their accessibility to a larger network of researchers.

Lastly, MICRA-Net was used to assist an Expert to perform a complex annotation task, that is the detection of small sparse objects (sections of genetically-labeled axons) in large fields of view of brain sections imaged with Scanning Electron Microscopy. Originally, this task was prone to identification errors and fatigue, limiting the performance of the Experts, and increasing inter-expert variability. MICRA-Net was successfully applied to assist the Experts at finding possible positive regions in the images. Instead of screening the whole field of view, Experts could focus their attention on less than 5% of the image and quickly decline or accept the proposed regions. This allowed an increase in the total number of detected regions of interest (genetically-tagged axons) by 25% while reducing the required annotation time for newly acquired images by 30 folds.

Precise annotations, even if obtained from trained Experts, are associated with inter-participant variability, especially when defining the boundaries (Figure 1). This variability needs to be assessed to characterize the annotated dataset and the precision of the neural network precision [3, 51]. We observed that image-level binary annotations can help to increase the consistency among Experts by reducing the complexity of the annotation task. By alleviating the annotation burden, an approach such as MICRA-Net can help increasing the accessibility of deep learning assisted quantitative image analysis in microscopy. As a whole, it can be used in multi-class detection, segmentation, counting, and classification tasks in bioimaging, for which a precisely annotated dataset is not available or tedious to obtain.

## Acknowledgments

Laurence Emond for F-Actin sample preparation and immunocytochemistry. Francine Nault, Charleen Salesse and Laurence Emond for the neuronal cell culture. Jonathan Marek and Renaud Bernatchez for the development of a custom Python annotation application. Thibault Dhellemmes for inter-expert axon DAB annotations in electron microscopy images. Christian Gagné and Marc-André Gardner for preliminary discussion on semantic segmentation. Annette Schwerdtfeger and Ana Gabela for careful proofreading of the manuscript. Funding was provided by grants from the Natural Sciences and Engineering Research Council of Canada (P.D.K. and F.L.C.), Canadian Institutes of Health Research (P.D.K.), Neuronex Initiative (National Science Foundation and Fond de recherche du Québec - Santé) (P.D.K., F.L.C.), CERVO Brain Research Center Foundation (F.L.C.), the Canadian Foundation for Innovation (P.D.K.). F.L.C. is a Canada Research Chair Tier II, Audrey Durand is a CIFAR AI Chair, and A.B. is supported by a PhD scholarship from the Fonds de Recherche Nature et Technologie Quebec (FRQNT) and an excellence scholarship from the FRQNT strategic cluster UNIQUE.

## Author contributions

A.B. and F.L.C. designed the approach. A.B. implemented the neuronal network architectures, generated the modified MNIST dataset, created the annotation application for the user study and performed all deep learning experiments. A.B., A.D. and F.L.C analysed the results. F.L.C. acquired and annotated the F-actin dataset. C.V.L.D. and M.P. generated and provided the annotated electron microscopy dataset. F.L.C., A.D. and P.D.K. supervised the project. F.L.C, A.D. and A.B. wrote the manuscript.

## Data and code availability

The MNIST, Cell Tracking Challenge, and P. Vivax datasets are all publicly available online. The F-actin dataset is available at the following website: https://s3.valeria.science/flclab-micranet/index.html. The Electron Microscopy dataset included in this study is available from the corresponding author upon reasonable request. All relevant material related to this paper is available on the following website: https://s3.valeria.science/flclab-micranet/index.html. Open source code for the MICRA-Net approach is available online: https://github.com/FLClab/MICRA-Net

## 4 Methods

### 4.1 MICRA-Net

#### 4.1.1 Architecture

Figure 2a shows the schematic representation of the MICRA-Net architecture. MICRA-Net is based on the encoder part of a U-Net [31]. The rationale is that U-Net is an established method to solve different analysis tasks (e.g. segmentation, localization, detection) on biomedical datasets. Each depth of the network contains two blocks of convolutions (kernel size of 3) followed by batch normalization, and ReLU activation. The number of filters in the convolutional layers is doubled after maxpooling (stride and kernel size of 2) to increase the richness of the representation. The number of filters for each layer is {32, 64, 128, 256}. Global maxpooling on the output layer allows a reduction of the dimensionality and a fully connected layer (FCL) is used to provide a classification prediction. Dropout (probability of 0.5) is applied on the input features of the FCL.

At inference, MICRA-Net predicts a whole image target from a given sample. Then, from each activated class *c*, a local map *L^l^* is calculated from the weighted combination of the activation map *A^l,k^* and the mean gradient 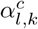 of each *l* layer [28]. The mean gradient 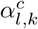 is calculated from the backpropagated class activation *y^c^*

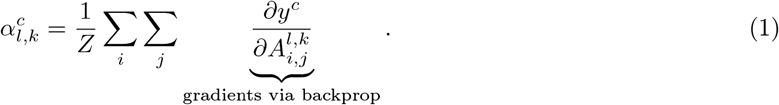

The local map *L^l^* is calculated as the linear combination of the activation map and the mean gradient of each layer of convolutions in the network

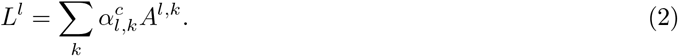

Since MICRA-Net produced spatially reduced feature maps, local maps were upsampled using nearest neighbor interpolation to match the input image size of 256 × 256 pixel. These images were then normalized in the range [0, 1] using a min-max scaling. ReLU activation is applied on the last layer (*L*^8^) of the network, as in the seminal implementation of Grad-CAM [28], to be used for the coarse segmentation. Local maps from layers *L*^1–7^ (Figure 2a-c) were concatenated into a feature space and retrieved the first principal component of every pixel using principal component analysis (PCA) [52] decomposition to retain prominent information from the feature space. The network was built and trained with the PyTorch library [53].

To facilitate the analysis of new images using MICRA-Net, a graphical user interface (GUI) is provided to qualitatively analyse the influence of each local map (Supplementary Fig. 30). While the implementation of MICRA-Net uses layers *L*^1–7^ with a PCA decomposition of the resultant feature space, the GUI allows to arbitrarily combine different local maps of the MICRA-Net architecture and threshold the resultant detailed feature map.

#### 4.1.2 Training procedure

The general training procedure of the MICRA-Net architecture are reported within this section. For specific training details for each dataset, see Supplementary Notes 1-5. MICRA-Net was trained using the Adam optimizer with a learning rate specific to each dataset and other default parameters [54]. A learning rate scheduler was used to reduce the learning rate of the optimizer with a minimal possible learning rate of 1 × 10^−5^. The number of training epochs was adapted to the specific dataset (Supplementary Tab. 22-26). Early stopping was used to reduce overfitting. Unless otherwise specified, we used binary cross entropy with logits loss. We kept the model with the best generalization properties on the validation set (calculated from the objective loss function).

Data augmentation was used to increase the performance of the network. Refer to Supplementary Tab. 22-26 for a detailed data augmentation procedure for each dataset. All operations were applied in a random order with a probability of 50%.

#### 4.1.3 Auxiliary tasks

This section presents how MICRA-Net can be used to solve the common auxiliary tasks in microscopy images.

##### Classification

The classification task is used on all presented dataset in the paper. It serves as a guideline to validate the representation capability of MICRA-Net. The classification task is solved by design using MICRA-Net since it is trained using a classification task. The prediction from MICRA-Net are mapped in the [0, 1] range using a sigmoid function.

##### Semantic segmentation

The semantic segmentation task is solved on all presented dataset in the paper. This task is solved by first extracting a detailed semantic feature map as described in Section 4.1.1. The semantic segmentation masks are obtained by thresholding the resultant semantic feature map using common thresholding algorithm (*e.g.* Otsu or percentile thresholding). The dataset specific thresholding is detailed in Supplementary Notes 1-5.

##### Detection

The detection task on the P. Vivax and EM microscopy dataset is solved by predicting the probability of presence of an object on all extracted crops. The overlap between the crops is of 75% in both directions. Overlapping crops are averaged and reassigned to an output feature map of the same shape as the image. The detection threshold is inferred from the validation set using a precision-recall curve.

##### Semantic instance segmentation

The semantic instance segmentation task is required on the Cell Tracking Challenge dataset. MICRA-Net is required to predict i) the presence of an object and ii) the contact between objects. The grad-CAMs of the activated objects are extracted from the architecture and combined using a principal component analysis (PCA) as presented in Section 4.1.1. If a contact is predicted on an image, the grad-CAM from L8 which contains the prominent information of the contact is extracted. The contact feature map is subtracted from the object feature map as in some fully-supervised techniques [27]. An Otsu threshold is used to generate the semantic segmentation masks of the instances.

### 4.2 Datasets

#### 4.2.1 Modified MNIST dataset

We generated the modified MNIST training dataset by randomly sampling *N* digits from the original MNIST training dataset and randomly distributed them on a 256 × 256 pixel field of view. To avoid overlap between digits we used a random Poisson disc sampling algorithm with a radius size of 25 pixels [55]. The number of digits *N* was uniformly sampled from {1, 2, 3, 4, 5, 10, 15, 20, `Max`}, where `Max` corresponds to the maximum number of digits that can be placed without overlap. A rotation of ±30° uniformly sampled was applied to the digits before placement on the image. We applied, in a random order, a Gaussian blur with sigma uniformly sampled in [0, 2 [and artificial normalized Poisson noise with 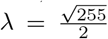. The resulting image intensities were clipped to lie in [0, 1]. Using this technique, we generated 2000 and 1000 images for training and validation respectively.

The modified MNIST testing dataset consists of 1000 images of handwritten digits sampled from the original MNIST testing dataset. As for the training dataset, we also applied, in a random order, Gaussian blur and artificial normalized Poisson noise sampled as before.

#### 4.2.2 F-actin dataset

The F-actin dataset was generated by using a sliding window of size 256 × 256 pixel with a stride of 192 pixels over 260 complete images with an approximate size of 1000 × 1000 pixel. Since the super-resolution microscopy images used are mostly composed of background, we set out to keep the crops containing at least 10% of dendritic area thereby reducing the number of crops to identify. The dendritic mask was obtained from the foreground detection on the confocal imaged of the dendritic marker MAP2 using a global Otsu thresholding on the normalized Gaussian blurred image [2, 35]. The sigma parameter of the Gaussian blur was set to 20 pixels as it provided suitable dendrite detection over a wide range of images. We next annotated each generated crop as being positive to the presence of the F-actin periodical lattice or longitudinal fibers. The resulting training dataset contained 3832 crops (256 × 256 pixel, 897 images positive to the periodical lattice and 1456 positive to the longitudinal fibers), the validation dataset contained 1287 crops (405 positive to periodical lattice and 377 positive to fibers), and the testing dataset contained 416 crops (83 positive to periodical lattice and 132 positive to fibers). The images were rescaled to lie in the [0, 1] interval. The maximum value for scaling (`max`) was obtained by sampling the maximal value of all training images from which we calculated the median in addition to 3 standard deviation. The minimum value was calculated as the median of minimas (`min`). To ensure a proper scaling of the images we also added a scaling factor of 0.8

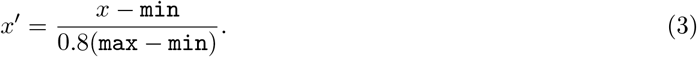

To evaluate the segmentation performance of the trained models, an Expert precisely highlighted the contours of the structures in 50 images (25 images positive to periodical lattice and 25 images positive to fibers) randomly sampled from the testing set. This small segmentation dataset only served to compare the segmentation performance from the MICRA-Net, weakly-supervised baselines (U-Net, Mask-R-CNN, Ilastik), and User-Study.

#### 4.2.3 Cell Tracking Challenge dataset

We selected 6 cell line datasets from the Cell Tracking Challenge (CTC) [8]: the DIC-C2DH-HeLa dataset which was acquired using differential interferometry contrast microscopy, three non-synthetic fluorescence microscopy datasets (Fluo-C2DL-MSC, Fluo-N2DH-GOWT1, and Fluo-N2DL-HeLa) and two phase contrast microscopy datasets (PhC-C2DH-U373, and PhC-C2DL-PSC). All original images were rescaled in the [0, 1] range using a per image min-max scale. We then resized each image and associated precise annotations according to the specific needs using bi-linear interpolation and nearest neighbors respectively with the `Scikit-Image` [56] Python library (Supplementary Table 6 for scaling factors). We used a sliding window of size 128 × 128 pixel or 256 × 256 pixel with a 25% overlap between crops in both directions. Using this sliding window technique yielded a total of 27,106 positive crops and 3,364 negative crops for the 256 × 256 pixel crops resized to have an effective pixel size of 0.5 μm. The sliding window with size 128 × 128 pixel crops and resized to have single cells in the field of view yielded a total of 66,466 positive crops (20,724 positive to contact) and 88,722 negative crops for training and 17,621 positive crops (5,606 positive to contact) and 22,279 negative crops for validation. We simulated weak annotations from the precise contours of the cells provided in the original CTC dataset by identifying an image crop as positive if the corresponding annotated crop contained at least the size of the average annotated cell, and negative otherwise. To evaluate the segmentation and detection tasks, we manually segmented 4 images randomly sampled per cell line in the testing set.

#### 4.2.4 P. Vivax dataset

We used image set BBBC041v1, available from the Broad Bioimage Benchmark Collection [33]. The complete dataset contained 1327 3-channel images and was already split into a training (1207 images) and testing (120 images) set. The dataset is composed of blood smears that were stained with Giemsa reagent [39] and acquired on three different brightfield microscopes from three different laboratories. All blood smears (infected or uninfected) were annotated using bounding boxes. The blood smears were later classified as infected (gametocytes, rings, trophozoites, and schizonts) or uninfected (red blood cells, and leukocytes) by an Expert. The task was to differentiate infected from uninfected blood smears. The dataset is highly unbalanced towards red blood cells which composes over 95% of the annotated cells.

For training and testing, we applied a whitening normalization (null mean and standard deviation of 1) to each image (and channel) to minimize the impact of a very different intensity distribution. The binary targets for training were generated using the provided bounding boxes. A crop was considered as positive if it contained at least 5% of overlap with an infected cell, otherwise as negative. The crops were 256 × 256 pixel.

We manually extracted and precisely annotated all infected cells in the testing set resulting in 303 small crops of size 256 × 256 pixel centered on the cell of interest.

#### 4.2.5 Scanning Electron Microscopy dataset

The dataset contained 92 images of 10, 240 × 10, 240 pixel for training, 66 for validation, and 44 for testing. An Expert annotated the images using positional markers to locate the Axon DAB markers. On average the large fields of view contained 3 small detections (113 × 113 pixel, between 1 and 10 detections per image). This resulted in an annotation time of approximately 30 minutes per field of view. Training and inference was performed on 512 × 512 pixel size crops. The dataset contained all positive crops (1024 × 1024 pixel, centered on the Axon DAB markers), and all negative crops (without overlap). To manually annotate the images the Expert inverted the acquired images. Hence, we provided MICRA-Net with the inverted image to mimic the Expert task. We rescaled the provided 8-bit depth images in the [0, 1] range by dividing by a scalar value of 255.

All Axon DAB markers were extracted from the testing set (170 positive markers) and an Expert carefully identified their contours.

### 4.3 Evaluation procedure

#### 4.3.1 Classification

The classification accuracy of MICRA-Net was evaluated by inferring the testing images. To quantitatively assess the performances, the classification accuracy was calculated for each trained model. We reported the mean ± standard deviation of the trained models.

#### 4.3.2 Detection

The centroid of each detected object was obtained from MICRA-Net by using the dataset specific procedures detailed in Supplementary Notes 1-5. Each detected centroid was associated with the centroid of objects in the ground truth mask using the Hungarian algorithm [57] with a maximal distance of *N* pixels, where *N* is approximately the object radius. In this context, an associated detected object is considered as a true positive, a non-associated detected object is a false positive, and a missed ground truth object is a false negative. To evaluate the detection capability of MICRA-Net, we reported the F1-score. For a quantitative comparison, we repeated the evaluation for each trained model. We then bootstrapped the average of the trained models to show the bootstrapped mean and 95% confidence interval (10 000 repetitions).

#### 4.3.3 Segmentation

The segmentation performance of the trained models was evaluated using three common evaluation metrics: F1-score, Intersection Over Union (IOU), and the Symmetric Boundary Dice (SBD) [58]. If multiple instances of a model were trained on the same task, we bootstrapped the average of the trained models to show the bootstrapped mean and 95% confidence interval (10 000 repetitions).

#### 4.3.4 Instance segmentation

Prior to evaluation, we removed small objects (<20 × 20 pixels) from the segmentation mask and filled holes for all trained models. All segmentation masks were resized to the baseline scale (Supplementary Table 6) for proper comparison. The instance segmentation performance were evaluated using the method proposed by [27] (Supplementary Figures 19-22). Briefly, this method evaluates the detection and failures of the architecture dependant on the IOU. [27] used a minimal IOU of 0.5 to avoid multiple predicted objects to be associated with a ground truth object. The goal is to maximize the F1-score vs. IOU, while the failure modes should be minimized. We on the other hand solved the association between the ground truth and predicted objects using the Hungarian algorithm [57], which allowed to report the performance and failure modes across the entire range of IOU. Using a broader range of IOU allows to report the performance in instance detection and segmentation. The normalized area under the resultant curves for each trained model is bootstrapped to obtain the mean and 95% confidence interval (10 000 repetition) and is reported in Figure 4.

#### 4.3.5 Custom performance metrics

The F-actin periodical lattice is detected as an oscillating pattern between high- and low-intensity stripes with 180-190 nm periodicity [32]. We designed a metric that would take this periodicity into account to evaluate the MICRA-Net detailed segmentation performance. We computed, as a baseline, the Fourier transform (FT) of the original image (FT*_b_*) and the FT of the segmented regions: for the Expert (FT*_e_*), and for the predicted segmentation masks (FT_pred_). The variation from the baseline was computed as the difference in the FT spectrum, for spatial frequencies in the range [170, 200[nm, between FT_e,pred_ and FT*_b_* over the sum of FT*_b_*. A smaller absolute difference between the variation of the Expert and the variation of the predicted mask implies more similar segmentation.

Since F-actin fibers are contiguous and have a high intensity on the dendrites, we designed a metric that would use the distribution of pixels under a segmented mask. The rational behind this metric is that the F-actin nanostructures on dendrites are composed of both high- and low-intensity pixels. Since F-actin fibers have high intensities, a detailed segmentation of fibers would imply few low intensity pixels annotated, while a coarse segmentation would introduce more low-intensity identified pixels. Hence, we considered a pixel within the segmentation mask as part of a fiber if its value was superior to a given threshold. We calculated this threshold by first measuring the 25^th^ percentile of pixel intensities outside of the Expert mask for all images. We then extracted the 90^th^ percentile intensity values from all images containing F-actin fibers. This resulted in a threshold between high- and low-intensity pixels within the dendritic mask of 9.

### 4.4 User-Study

We conducted two different User-Study in this paper, one for the F-actin nanostructure segmentation and one for the instance segmentation on the Cell Tracking Challenge. All participants were familiar with bio-medical images.

#### 4.4.1 F-Actin segmentation

We performed a User-Study in which six participants highlighted the contours of the F-actin periodical lattice and longitudinal fibres on a small dataset of 50 images using polygonal bounding boxes. We used polygonal bounding boxes as this annotation method reduces the time required by a participant by more than 3 folds compared to precisely identifying the boundaries of the structures (Supplementary Fig. 11). We used our own annotation application that was optimized for this type of task. Annotation of the full dataset required approximately 40 minutes for the participants. The averaged performance of the six participants was compared to MICRA-Net using F1-score, IOU, and SBD.

#### 4.4.2 Cell Tracking Challenge instance segmentation

A User-Study was conducted using the Cell Tracking Challenge to analyse the required time per cells and the achievable performance of inter-participant annotation for such task. The User-Study consisted in the annotation the 24 testing image using different level of supervision (precise, bounding boxes, and points). For each level of supervision, the participants were asked to annotate a quarter of the testing image, which was the same for all participants. The image intensity scale was set at a constant value for all participants. The participants used the Fiji software to annotate the images. The median of the participant scores on the testing set are reported, as well as the inter-participant scores. The time required by the participant to annotate each image was recorded, which allowed to calculate the time per cell for each cell-line.

### 4.5 In-house datasets acquisition

#### 4.5.1 Cell culture, Immunostaining and STED imaging for F-actin imaging

Before dissection of hippocampi, neonatal Sprague Dawley rats were sacrificed by decapitation, in accordance to the procedures approved by the animal care committee of Université Laval. Dissociated cells were plated on poly-d-lysine coated glass coverslips, fixed and immunostained as described previously [2]. F-Actin was stained with Phalloidin-STAR635 (Abberior GmbH, Germany). Dendrites Microtubule-Associated-Protein (MAP2) [2]. STED images of the F-Actin nanostructures were acquired on a 4 color Abberior Expert-Line STED microscope (Abberior Instruments GmbH, Germany), equiped with a 100x 1.4 NA oil objective and using pulsed (40 MHz) excitation (640 nm) and depletion (775 nm) lasers. Fluorescence was detected with an Avalanche Photodiode (APD) and a ET685/70 (Chroma, USA) fluorescence filter. Pixel size was set to 20 nm.

#### 4.5.2 Animals and stereotaxic injections for scanning electron microscopy dataset

This study was carried out on 3-month-old mice, weighing 25-35g. Animals were housed under a 12h light dark cycle with water and food ad libitum. All procedures were approved by the *Comité de Protection des Animaux de l’Université Laval*, in accordance with the Canadian Council on Animal Care’s Guide to the Care and Use of Experimental Animals (Ed2), and with the ARRIVE guidelines. Maximum efforts were made to minimize the number of animals used. Transgenic e-Pet Cre mice expressing Cre recombinase under the control of Fev promoter, known to be specific for serotonin (5-HT) neurons [59], were injected in the dorsal raphe nucleus (DRN) with 1 μl of AAV9-CAG-DIO-APEX2NES-WPRE. Stereotaxic injections were done using a 30° angle along the frontal plane at AP: −4.78; ML: +2.00 and DV: −3.20. In these injected transgenic mice, the small engineered peroxidase APEX2 [40] is specifically expressed in the cytosol/cytoplasm of 5-HT-infected neurons of the DRN and is used, in presence with hydrogen peroxide, to oxidize 3,3 Diaminobenzidine (DAB) chromogen that can readily be visible at the light and electron microscope levels.

#### 4.5.3 Tissue preparation for scanning electron microscopy dataset

After a period of 21 days following stereotaxic injection, mice were anesthetized with a mixture of ketamine (100 mg*/*kg) and xylazine (10 mg*/*kg) and transcardially perfused with 50 ml of phosphate-buffered-saline (PBS: 50 mM at pH 7.4) followed by 150 ml of 4% paraformaldehyde (PFA) and 1% glutaraldehyde diluted in phosphate buffer (PB; 100 mM at pH 7.4). Brains were dissected out, post-fixed for 24h in the same fixative solution and cut with a vibratome (model VT1200; Leica, Germany) into 50 *μ*m-thick frontal sections, which were serially collected in sodium phosphate buffer saline (PBS, 100 mM, pH 7.4). Frontal brain sections at the level of the subthalamic nucleus (STN) were processed to reveal the presence of APEX2 in axons arising from DRN-infected neurons using 3,3’diaminobenzidine (DAB; catalog no. D5637; Sigma-Aldrich) as the chromogen. Briefly, selected 50 *μ*m-thick sections were washed 3 times in PBS and then twice in Tris. Sections were then incubated for 1h in 0.05% DAB solution diluted in Tris, then for 1h in 0.05% DAB solution containing 0.015% hydrogen peroxide (H_2_O_2_). Sections were then rinsed twice in Tris and 3 times in PBS. Sections were temporally mounted in PBS and coversliped for light microscope examination. STN sections containing DAB-labeled axons were selected for further processing. These sections were washed 3 times in PB, then incubated during 1h in 2% osmium tetroxide diluted in 1.5% potassium ferrocyanide solution. They were then washed 3 times in ddH_2_O, incubated for 20 min in 1% thiocarbohydrazide (TCH) solution and washed again 3 times in ddH_2_O. Sections were placed 30 min in 2% osmium tetroxide and washed 3 times in ddH_2_O. Sections were then dehydrated in ethanol and propylene oxide and flat-embedded in Durcupan (Electron microscopy Science). Areas of interest were cut from embedded sections and glued to the tip of resin blocks. Blocks were cut with an ultramicrotome (Leica EM UC7) in ultrathin sections (80 nm), which were serially collected on silicon-coated 10 × 10 mm chip wafer (Ted Pella, Inc; #16006).

#### 4.5.4 Scanning electron microscopy (SEM)

Serial sections were imaged in a SEM (Zeiss Gemini 540) with the help of the ATLAS acquisition software. Images were acquired at a resolution of 5 nm/pixel, using acceleration voltage of 1.4 kV and current of 1.2 nA. Serial sections acquisitions produced a stack of 38 rectangle images of 25370 × 25633 pixel (126.850 × 128.165 microns) taken out of 38 ultrathin sections. In addition, a large single section acquisition was acquired and produced a single trapezoidal image of 31065 pixels for the small base (155.329 microns), 91393 pixel for the large base (456.967 microns) and 53161 pixels for the height (265.809 microns). All acquired images were subdivided into overlapping square tiles of 10240 *×* 10240 pixel (51.2 × 51.2 microns).

### 4.6 Statistical assessment using resampling

Resampling was used as a statistical test to verify the statistical difference between two groups [60]. Statistical analysis was performed using a randomization test with the null hypothesis being that the different conditions (A, B) belong to the same distribution. The absolute difference between mean values of A and B was calculated (*D*_gt_ = |*μ_A_ − μ_B_*|). For the randomization test, each value belonging to A and B was randomly reassigned to A’ and B’, with the sizes of A’ and B’ being *N_A_* and *N_B_*, respectively. The absolute difference between the mean values of A’ and B’ was determined (*D*_rand_ = |*μ_A′_ − μ_B′_*|) and the randomization test was repeated 10 000 times. The obtained distribution was compared with the absolute difference of the mean of A and B (*D*_gt_) to verify the null hypothesis.

When the number of groups was greater than 2, the F-statistic was sampled from each group using a resampling method. The F-statistic was calculated from all groups (A, B, C, etc.) as a ground truth (*F*_gt_). Each value was randomly re-assigned to new groups (A’, B’, C’, etc.) where group X’ has the same size as group X. The F-statistic of newly formed groups (*F*_rand_) was calculated and this process was repeated 10 000 times. We compared *F*_rand_ with *F*_gt_ to confirm the null hypothesis that the groups have the same mean distribution. When the null hypothesis was rejected, *i.e.* at least one group did not have the same mean distribution, we compared each group in a one-to-one manner using the randomization test described above. In all cases, a confidence level of 0.05 was used to reject the null hypothesis. Since the precision of the calculation of the *p*-value is limited to 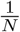, where *N* in the number of repetitions, we report a *p*-value of <1.0000 × 10^−4^ instead of 0.

### 4.7 Evaluation of required decisions and time for fully-supervised training

#### F-actin

The number of decisions for a fully-supervised training dataset was estimated as the mean number of edge pixels in the 50 precisely annotated images multiplied by the total number of positive crops. The mean annotation time per crop was calculated using the precisely annotated dataset.

#### Cell Tracking Challenge

The mean image annotation time of 900 seconds was obtained from the precise annotation of each image of the testing set.

#### P. Vivax

The annotation time for fully-supervised annotations was estimated at 2 minutes per image from the precise annotation of 10 images.

#### Electron Microscopy

The required annotation time was calculated as the average time required by the Expert per image (30 minutes per image, 156 images) to detect all axon DAB markers. We added 14 seconds (calculated from highlighting the contours of the Axon DAB regions on the testing set) for each positive detection (537 detections) to account for precise annotation.

## References

[1] Schermelleh, L. et al. Super-resolution microscopy demystified. Nature Cell Biology 21, 72 (2019).

[2] Lavoie-Cardinal, F. et al. Neuronal activity remodels the f-actin based submembrane lattice in dendrites but not axons of hippocampal neurons. Scientific Reports (Nature Publisher Group) 10 (2020).

[3] Schlegl, T., Seeböck, P., Waldstein, S. M., Langs, G. & Schmidt-Erfurth, U. f-anogan: Fast unsupervised anomaly detection with generative adversarial networks. Medical image analysis 54, 30–44 (2019).

[4] LeCun, Y., Bengio, Y. & Hinton, G. Deep learning. Nature 521, 436 (2015).

[5] Gupta, A. et al. Deep learning in image cytometry: a review. Cytometry Part A 95, 366–380 (2019).

[6] Caicedo, J. C. et al. Nucleus segmentation across imaging experiments: the 2018 Data Science Bowl. Nature Methods 16, 1247–1253 (2019).

[7] Moen, E. et al. Deep learning for cellular image analysis. Nature methods 1–14 (2019).

[8] Ulman, V. et al. An objective comparison of cell-tracking algorithms. Nature methods 14, 1141–1152 (2017).

[9] Falk, T. et al. U-net: deep learning for cell counting, detection, and morphometry. Nature Methods 16, 67 (2019).

[10] He, K., Gkioxari, G., Dollár, P. & Girshick, R. Mask R-CNN. arXiv:1703.06870 [cs] (2018). 1703.06870.

[11] Kromp, F. et al. An annotated fluorescence image dataset for training nuclear segmentation methods. Scientific Data 7, 262 (2020).

[12] Stringer, C., Wang, T., Michaelos, M. & Pachitariu, M. Cellpose: A generalist algorithm for cellular segmentation. Nature Methods 18, 100–106 (2021).

[13] Wilhelm, B. G. et al. Composition of isolated synaptic boutons reveals the amounts of vesicle trafficking proteins. Science 344, 1023–1028 (2014).

[14] Cheplygina, V., de Bruijne, M. & Pluim, J. P. W. Not-so-supervised: A survey of semi-supervised, multi-instance, and transfer learning in medical image analysis. Medical Image Analysis 54, 280–296 (2019).

[15] Papandreou, G., Chen, L.-C., Murphy, K. P. & Yuille, A. L. Weakly-and semi-supervised learning of a deep convolutional network for semantic image segmentation. In Proceedings of the IEEE international conference on computer vision, 1742–1750 (2015).

[16] Khoreva, A., Benenson, R., Hosang, J., Hein, M. & Schiele, B. Simple does it: Weakly supervised instance and semantic segmentation. In Proceedings of the IEEE conference on computer vision and pattern recognition, 876–885 (2017).

[17] Xu, J., Schwing, A. G. & Urtasun, R. Tell me what you see and i will show you where it is. In Proceedings of the IEEE conference on computer vision and pattern recognition, 3190–3197 (2014).

[18] Pesce, E. et al. Learning to detect chest radiographs containing pulmonary lesions using visual attention networks. Medical image analysis 53, 26–38 (2019).

[19] Rajchl, M. et al. Deepcut: Object segmentation from bounding box annotations using convolutional neural networks. IEEE transactions on medical imaging 36, 674–683 (2016).

[20] Yang, L. et al. Boxnet: Deep learning based biomedical image segmentation using boxes only annotation. arXiv preprint arXiv:1806.00593 (2018).

[21] Lin, T.-Y. et al. Microsoft coco: Common objects in context. In European conference on computer vision, 740–755 (Springer, 2014).

[22] Vezhnevets, A., Ferrari, V. & Buhmann, J. M. Weakly supervised structured output learning for semantic segmentation. In Proceedings of the IEEE conference on computer vision and pattern recognition, 845–852 (IEEE, 2012).

[23] Dubost, F. et al. Weakly supervised object detection with 2d and 3d regression neural networks. arXiv preprint arXiv:1906.01891 (2019).

[24] Li, J. et al. An em-based semi-supervised deep learning approach for semantic segmentation of histopathological images from radical prostatectomies. Computerized Medical Imaging and Graphics 69, 125–133 (2018).

[25] Kraus, O. Z., Ba, J. L. & Frey, B. J. Classifying and segmenting microscopy images with deep multiple instance learning. Bioinformatics 32, i52–i59 (2016).

[26] Chatterjee, B. & Poullis, C. Semantic segmentation from remote sensor data and the exploitation of latent learning for classification of auxiliary tasks. arXiv preprint arXiv:1912.09216 (2019).

[27] Caicedo, J. C. et al. Evaluation of Deep Learning Strategies for Nucleus Segmentation in Fluorescence Images. Cytometry Part A 95, 952–965 (2019).

[28] Selvaraju, R. R. et al. Grad-cam: Visual explanations from deep networks via gradient-based localization. In Proceedings of the IEEE International Conference on Computer Vision, 618–626 (2017).

[29] Berg, S. et al. ilastik: interactive machine learning for (bio)image analysis. Nature Methods (2019). URL https://doi.org/10.1038/s41592-019-0582-9.

[30] LeCun, Y., Bottou, L., Bengio, Y., Haffner, P. et al. Gradient-based learning applied to document recognition. Proceedings of the IEEE 86, 2278–2324 (1998).

[31] Ronneberger, O., Fischer, P. & Brox, T. U-net: Convolutional networks for biomedical image segmentation. In International Conference on Medical image computing and computer-assisted intervention, 234–241 (Springer, 2015).

[32] Xu, K., Zhong, G. & Zhuang, X. Actin, spectrin, and associated proteins form a periodic cytoskeletal structure in axons. Science 339, 452–456 (2013).

[33] Ljosa, V., Sokolnicki, K. L. & Carpenter, A. E. Annotated high-throughput microscopy image sets for validation. Nature methods 9, 637–637 (2012).

[34] Kromp, F. et al. Deep Learning architectures for generalized immunofluorescence based nuclear image segmentation. arXiv:1907.12975 [cs, q-bio] (2019). 1907.12975.

[35] Otsu, N. A threshold selection method from gray-level histograms. IEEE Transactions on Systems, Man, and Cybernetics 9, 62–66 (1979).

[36] Hung, J. & Carpenter, A. Applying faster r-cnn for object detection on malaria images. In Proceedings of the IEEE conference on computer vision and pattern recognition workshops, 56–61 (2017).

[37] Belthangady, C. & Royer, L. A. Applications, promises, and pitfalls of deep learning for fluorescence image reconstruction. Nature methods 1–11 (2019).

[38] Weigert, M. et al. Content-aware image restoration: pushing the limits of fluorescence microscopy. Nature methods 15, 1090–1097 (2018).

[39] Barcia, J. J. The giemsa stain: its history and applications. International journal of surgical pathology 15, 292–296 (2007).

[40] Lam, S. S. et al. Directed evolution of apex2 for electron microscopy and proximity labeling. Nature methods 12, 51–54 (2015).

[41] Bekker, J. & Davis, J. Learning from positive and unlabeled data: a survey. Mach. Learn. 109, 719–760 (2020).

[42] Christiansen, E. M. et al. In silico labeling: predicting fluorescent labels in unlabeled images. Cell 173, 792–803 (2018).

[43] Caruana, R. Multitask learning. Machine learning 28, 41–75 (1997).

[44] Girshick, R. Fast r-cnn. In Proceedings of the IEEE international conference on computer vision, 1440–1448 (2015).

[45] Ruder, S. An overview of multi-task learning in deep neural networks. arXiv preprint arXiv:1706.05098 (2017).

[46] Mathis, A. et al. Deeplabcut: markerless pose estimation of user-defined body parts with deep learning. Nature neuroscience 21, 1281–1289 (2018).

[47] He, K., Girshick, R. & Dollár, P. Rethinking imagenet pre-training. In Proceedings of the IEEE international conference on computer vision, 4918–4927 (2019).

[48] Raghu, M., Zhang, C., Kleinberg, J. & Bengio, S. Transfusion: Understanding Transfer Learning for Medical Imaging. In Wallach, H. et al. (eds.) Advances in Neural Information Processing Systems 32, 3347–3357 (Curran Associates, Inc., 2019).

[49] Eliceiri, K. W. et al. Biological imaging software tools. Nature Methods 9, 697–710 (2012).

[50] Ouyang, W. et al. Analysis of the Human Protein Atlas Image Classification competition. Nature Methods 16, 1254–1261 (2019).

[51] Mazzara, G. P., Velthuizen, R. P., Pearlman, J. L., Greenberg, H. M. & Wagner, H. Brain tumor target volume determination for radiation treatment planning through automated MRI segmentation. International Journal of Radiation Oncology, Biology, Physics 59, 300–312 (2004).

[52] Hotelling, H. Analysis of a complex of statistical variables into principal components. Journal of educational psychology 24, 417 (1933).

[53] Paszke, A. et al. Automatic differentiation in pytorch. In 31st Conference on Neural Information Processing Systems (2017).

[54] Kingma, D. P. & Ba, J. Adam: A method for stochastic optimization. arXiv preprint arXiv:1412.6980 (2014).

[55] Cook, R. L. Stochastic sampling in computer graphics. ACM Transactions on Graphics (TOG) 5, 51–72 (1986).

[56] Van der Walt, S. et al. scikit-image: image processing in python. PeerJ 2, e453 (2014).

[57] Kuhn, H. W. The hungarian method for the assignment problem. Naval research logistics quarterly 2, 83–97 (1955).

[58] Yeghiazaryan, V. & Voiculescu, I. D. Family of boundary overlap metrics for the evaluation of medical image segmentation. Journal of Medical Imaging 5, 015006 (2018).

[59] Scott, M. M. et al. A genetic approach to access serotonin neurons for in vivo and in vitro studies. Proceedings of the National Academy of Sciences 102, 16472–16477 (2005).

[60] Good, P. I. Resampling Methods (Birkhäuser Basel, 2006), 3 edn.

