## Supplementary Material for "MICRA-Net: MICRoscopy Analysis Neural Network to solve detection, classification, and segmentation from a single simple auxiliary task"

#### Supplementary Figures and Tables

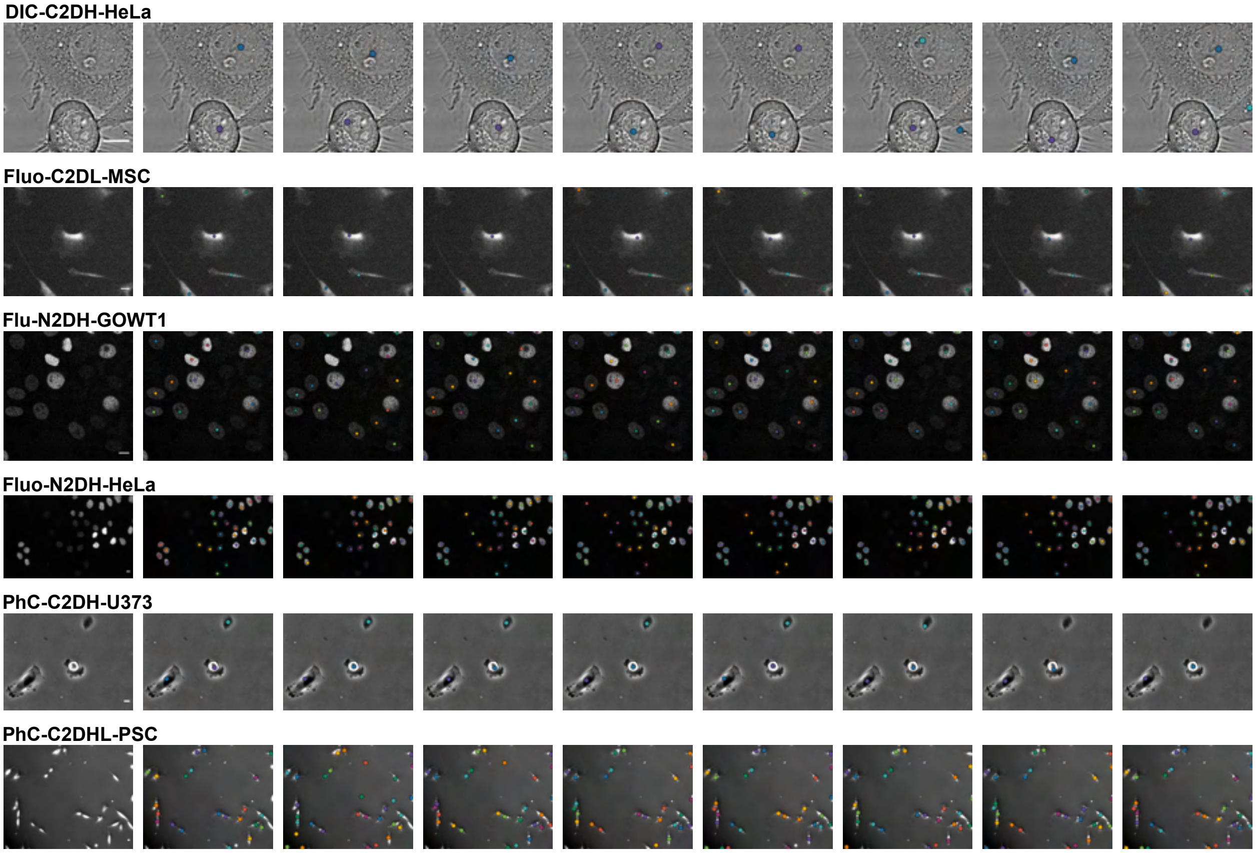


Supplementary Figure 1: Examples of points annotations sampled from the conducted User-Study on 6 different cell lines from the Cell Tracking Challenge [1]. The intensity scale is the same that was used for images shown to the participants during the task. The image on the left is the original image without annotations. Each other column correspond to the annotations of the same participant. Scale bar : 10 µm.

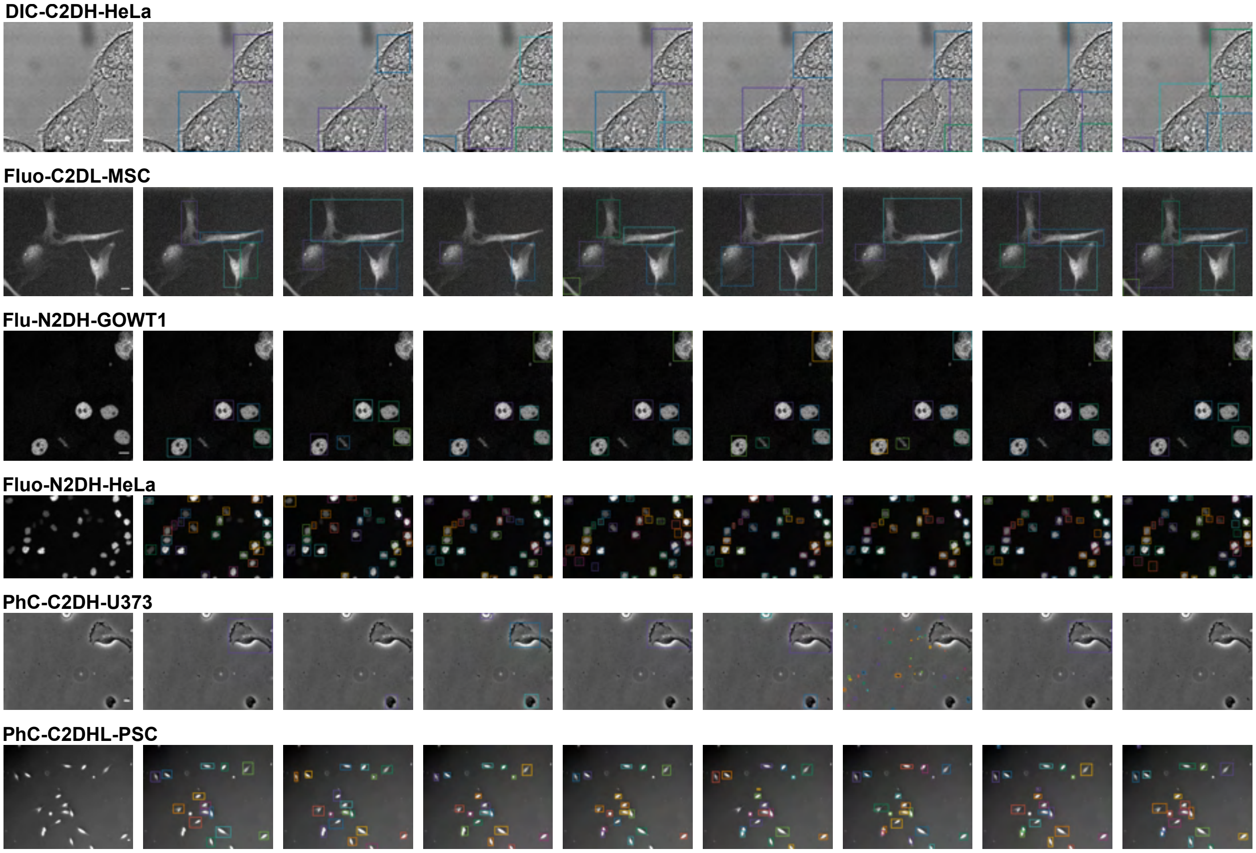


Supplementary Figure 2: Examples of bounding boxes annotations sampled from the conducted User-Study on 6 different cell lines from the Cell Tracking Challenge [1]. The intensity scale is shown as it was shown to the participants during the task. The image on the left is the original image without annotations. Each other column correspond to the annotations of the same participant. Scale bar : 10 µm.

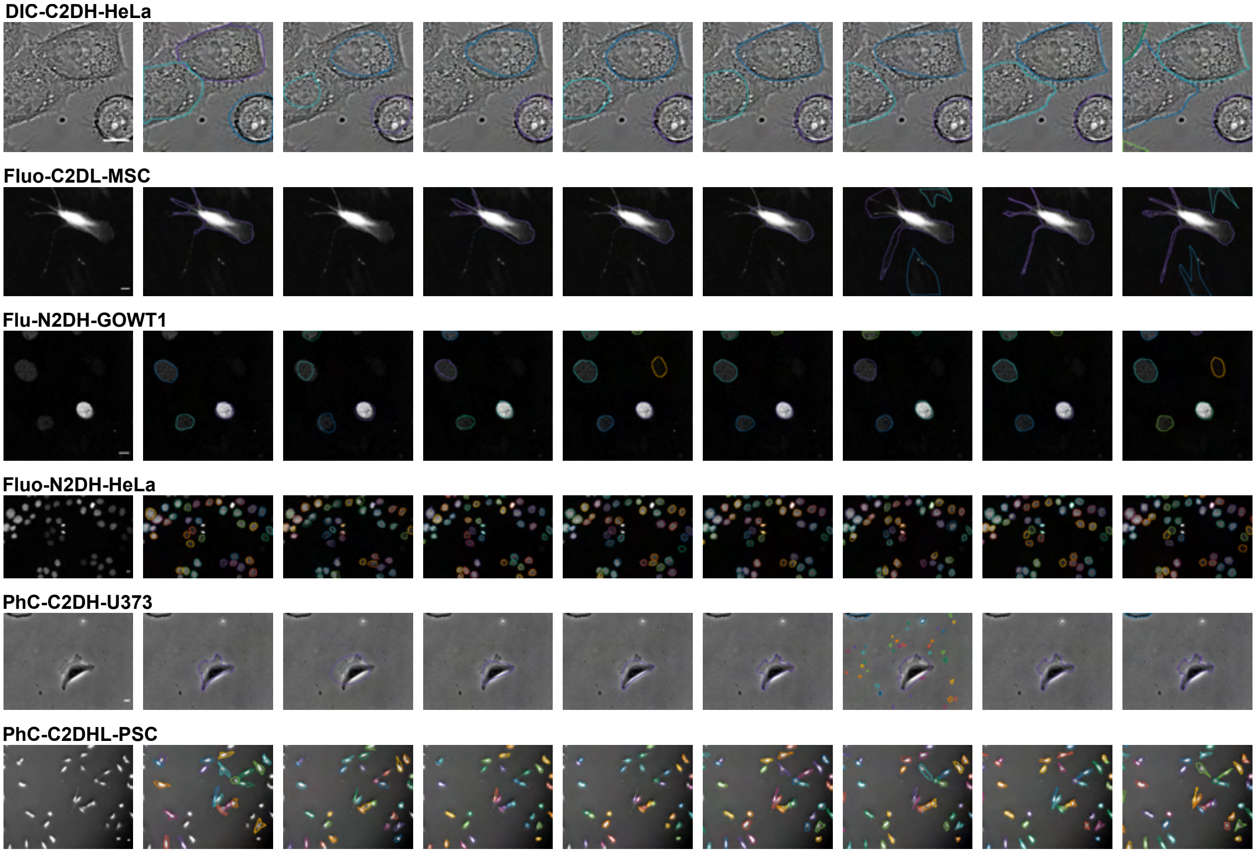


Supplementary Figure 3: Examples of precise contour annotations sampled from the conducted User-Study on 6 different cell lines from the Cell Tracking Challenge [1]. The intensity scale is shown as it was shown to the participants during the task. The image on the left is the original image without annotations. Each other column correspond to the annotations of the same participant. Scale bar : 10 µm.

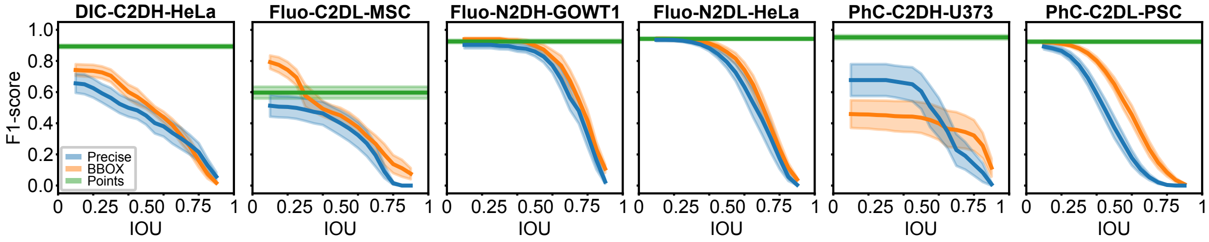


Supplementary Figure 4: Inter-participant variability from the User-Study on 6 selected cell lines of the Cell Tracking Challenge using three levels of supervision (precise, bounding boxes (BBOX), and points). Representative examples from the participants may be found in Supplementary Fig. 1-3. The inter-participant agreement was calculated using the F1-score as a function of IOU for precise (blue) and BBOX (orange) annotations in a all versus one manner [2] (see Supplementary Fig. 4). The F1-score for points annotation (green) was calculated with a maximal distance of association of 30 pixels. Plotted are the bootstrapped mean (line) and 95% confidence interval (shade, 10 000 repetitions).

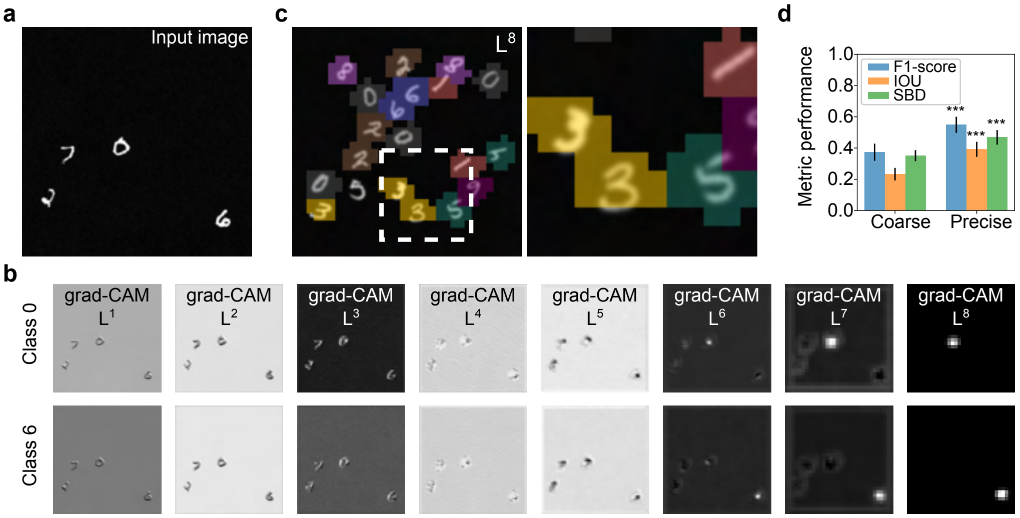


Supplementary Figure 5: Coarse segmentation implementation of MICRA-Net. a) Representative example image of the modified MNIST dataset. b) The gradient class activated maps (grad-CAM) are extracted for all activated class in the image from all layers of the network (image in a). Here are only shown grad-CAMs for activated class 0 and 6. c) Coarse segmentation maps of the digits of a representative image ($256 \times 256$ pixel) and insets (right, dashed white box) obtained by thresholding the $L^8$ layer of the network. The color code is the same as in Figure 2 and corresponds to the digit class. d) The segmentation performance between the coarse ($L^8$) and precise ($L^{1-7}$) method is compared using F1-score, Intersection Over Union (IOU) and Symmetric Boundary Dice (SBD). The bar graph shows the average and standard deviation over the 10 classes (Supplementary Fig. 6 for class-wise and density-wise performances). A significant increase is shown for the precise method over the coarse method for all calculated metrics (Supplementary Tab. 1).

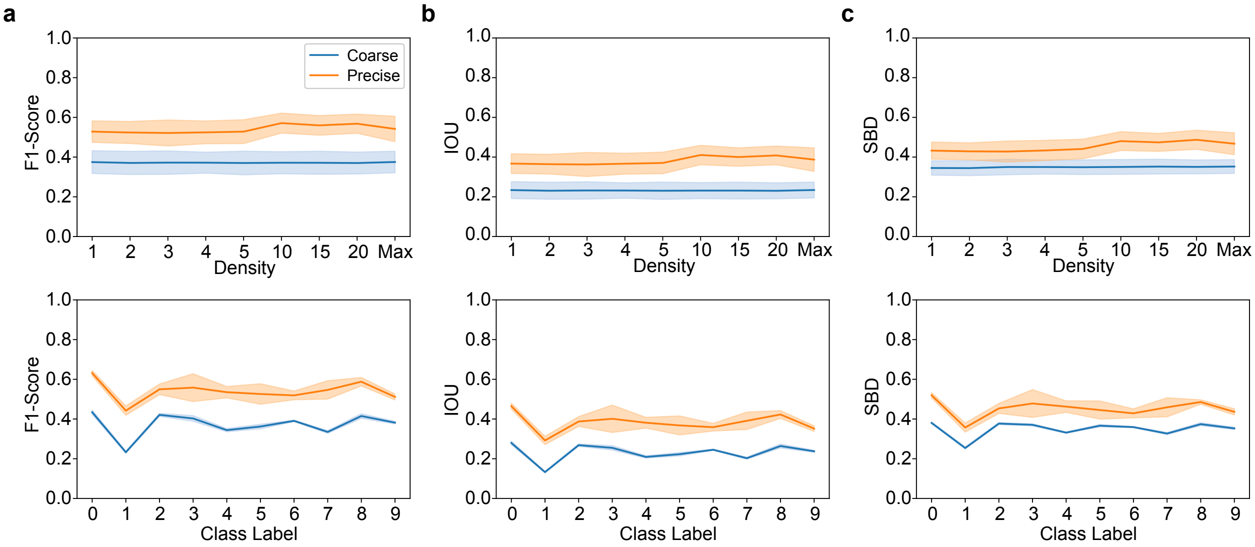


Supplementary Figure 6: Comparison of coarse and precise segmentations on the modified MNIST dataset for three different metrics: a) F1-Score, b) IOU, and c) SBD. Top row shows the performance for each metric depending on the number of digits in the field of view (Density) while bottom row shows the performance for each metric depending on the class. Solid lines and pale regions represent the mean and standard deviation respectively.

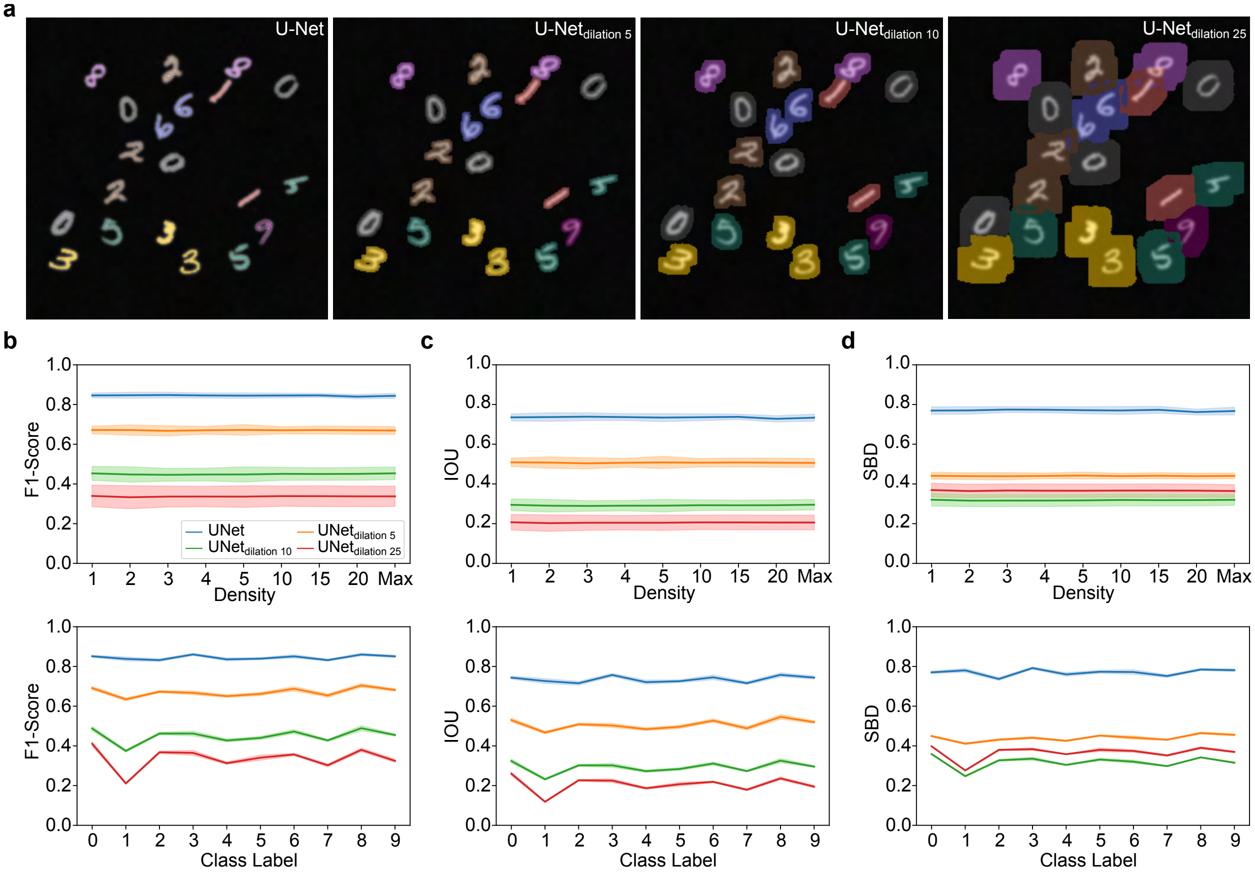


Supplementary Figure 7: U-Net performance trained in a fully- and weakly-supervised fashion on the modified MNIST dataset. a) Segmentation examples taken from the testing dataset with an increasing dilation of the MNIST binary digits to simulate weak supervision. The dilation is computed using a square structuring element of $\{5, 10, 25\}$ pixel. The different training schemes are compared using three performance metrics: b) F1-score, c) IOU, and d) SBD. Top and bottom rows display the per density and per class performance respectively. Solid lines and pale regions represent the mean and standard deviation respectively.

| **F1-score** | Coarse | Precise | U-Net | U-Net5 | U-Net10 | U-Net25 |
|---|---|---|---|---|---|---|
| Coarse | - | $1.0000 \times 10^{-4}$ | $<1.0000 \times 10^{-4}$ | $<1.0000 \times 10^{-4}$ | $<1.0000 \times 10^{-4}$ | $3.1080 \times 10^{-1}$ |
| Precise | $1.0000 \times 10^{-4}$ | - | $<1.0000 \times 10^{-4}$ | $<1.0000 \times 10^{-4}$ | $1.8000 \times 10^{-3}$ | $<1.0000 \times 10^{-4}$ |
| U-Net | $<1.0000 \times 10^{-4}$ | $<1.0000 \times 10^{-4}$ | - | $<1.0000 \times 10^{-4}$ | $<1.0000 \times 10^{-4}$ | $<1.0000 \times 10^{-4}$ |
| U-Net5 | $<1.0000 \times 10^{-4}$ | $<1.0000 \times 10^{-4}$ | $<1.0000 \times 10^{-4}$ | - | $<1.0000 \times 10^{-4}$ | $<1.0000 \times 10^{-4}$ |
| U-Net10 | $<1.0000 \times 10^{-4}$ | $1.8000 \times 10^{-3}$ | $<1.0000 \times 10^{-4}$ | $<1.0000 \times 10^{-4}$ | - | $<1.0000 \times 10^{-4}$ |
| U-Net25 | $3.1080 \times 10^{-1}$ | $<1.0000 \times 10^{-4}$ | $<1.0000 \times 10^{-4}$ | $<1.0000 \times 10^{-4}$ | $<1.0000 \times 10^{-4}$ | - |

| **IOU** | Coarse | Precise | U-Net | U-Net5 | U-Net10 | U-Net25 |
|---|---|---|---|---|---|---|
| Coarse | - | $<1.0000 \times 10^{-4}$ | $1.0000 \times 10^{-4}$ | $1.0000 \times 10^{-4}$ | $1.0000 \times 10^{-4}$ | $2.8040 \times 10^{-1}$ |
| Precise | $<1.0000 \times 10^{-4}$ | - | $<1.0000 \times 10^{-4}$ | $1.0000 \times 10^{-4}$ | $2.0000 \times 10^{-4}$ | $<1.0000 \times 10^{-4}$ |
| U-Net | $1.0000 \times 10^{-4}$ | $<1.0000 \times 10^{-4}$ | - | $<1.0000 \times 10^{-4}$ | $<1.0000 \times 10^{-4}$ | $<1.0000 \times 10^{-4}$ |
| U-Net5 | $1.0000 \times 10^{-4}$ | $1.0000 \times 10^{-4}$ | $<1.0000 \times 10^{-4}$ | - | $<1.0000 \times 10^{-4}$ | $<1.0000 \times 10^{-4}$ |
| U-Net10 | $1.0000 \times 10^{-4}$ | $2.0000 \times 10^{-4}$ | $<1.0000 \times 10^{-4}$ | $<1.0000 \times 10^{-4}$ | - | $<1.0000 \times 10^{-4}$ |
| U-Net25 | $2.8040 \times 10^{-1}$ | $<1.0000 \times 10^{-4}$ | $<1.0000 \times 10^{-4}$ | $<1.0000 \times 10^{-4}$ | $<1.0000 \times 10^{-4}$ | - |

| **SBD** | Coarse | Precise | U-Net | U-Net5 | U-Net10 | U-Net25 |
|---|---|---|---|---|---|---|
| Coarse | - | $1.7900 \times 10^{-2}$ | $<1.0000 \times 10^{-4}$ | $<1.0000 \times 10^{-4}$ | $7.0000 \times 10^{-4}$ | $4.2230 \times 10^{-1}$ |
| Precise | $1.7900 \times 10^{-2}$ | - | $2.0000 \times 10^{-4}$ | $4.5530 \times 10^{-1}$ | $1.0000 \times 10^{-4}$ | $2.5000 \times 10^{-3}$ |
| U-Net | $<1.0000 \times 10^{-4}$ | $2.0000 \times 10^{-4}$ | - | $<1.0000 \times 10^{-4}$ | $<1.0000 \times 10^{-4}$ | $<1.0000 \times 10^{-4}$ |
| U-Net5 | $<1.0000 \times 10^{-4}$ | $4.5530 \times 10^{-1}$ | $<1.0000 \times 10^{-4}$ | - | $<1.0000 \times 10^{-4}$ | $<1.0000 \times 10^{-4}$ |
| U-Net10 | $7.0000 \times 10^{-4}$ | $1.0000 \times 10^{-4}$ | $<1.0000 \times 10^{-4}$ | $<1.0000 \times 10^{-4}$ | - | $4.1000 \times 10^{-3}$ |
| U-Net25 | $4.2230 \times 10^{-1}$ | $2.5000 \times 10^{-3}$ | $<1.0000 \times 10^{-4}$ | $<1.0000 \times 10^{-4}$ | $4.1000 \times 10^{-3}$ | - |

Supplementary Table 1: Comparison of the performance metrics (F1-score, IOU, and SBD) for MICRA-Net (coarse and precise) and U-Net (weakly- and fully-supervised) segmentation on the modified MNIST dataset. The $p$-values are obtained by a resampling statistical test (10 000 repetitions) comparing each lines against each columns following a resampled ANOVA (10 000 repetitions, $p_{\text{F1-score}} =< 1.0000 \times 10^{-4}$, $p_{\text{IOU}} =< 1.0000 \times 10^{-4}$ and $p_{\text{SBD}} =< 1.0000 \times 10^{-4}$). Color code: increase (cyan), decrease (red), and no significant changes (black) of each line compared to each column of the metric scores.

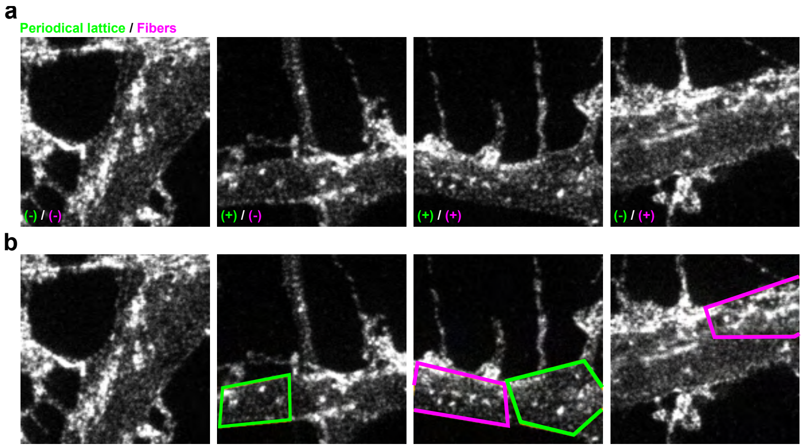


Supplementary Figure 8: Representative examples of 256 × 256 pixel crops sampled from the training set of the F-actin dataset. In a) positive (+) and negative (-) crops, and in b) the associated polygonal bounding box annotations are presented. The F-actin periodical lattice is in green, while longitudinal fibers are depicted in magenta. Each crop is 3.84 × 3.84 µm.

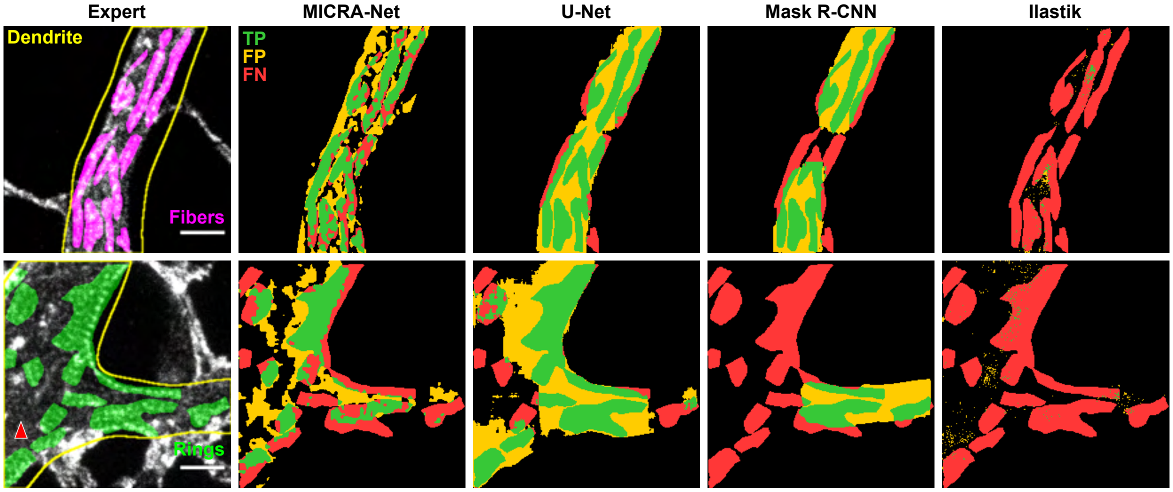


Supplementary Figure 9: Representative images of F-actin semantic segmentation on dendrites for both structures (fibers and periodical lattice [rings]). From left to right, the fine segmentation from the Expert, MICRA-Net, weakly supervised U-Net, weakly supervised Mask R-CNN and Ilastik are shown. The color code maps true positive (TP, green), false positive (FP, yellow) and false negative (FN, red) segmentation of each method compared to the fine Expert labels. A red arrow indicates a missed region in the periodical lattice image by the Expert. Scale bars $1 \mu m$.

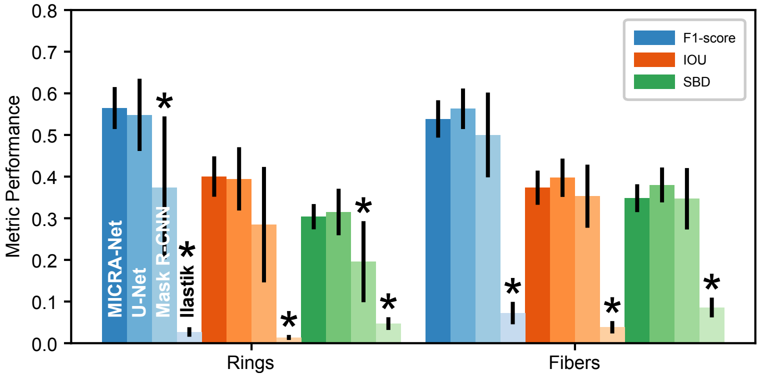


Supplementary Figure 10: F1-Score (blue), IOU (orange), and SBD (green) calculated for MICRA-Net (dark), weakly-supervised U-Net (medium-dark), weakly-supervised Mask R-CNN (medium-light) and Ilastik (light) on the F-actin dataset. Bar graphs and error bars show the bootstrapped mean and 95% confidence interval (10 000 repetitions). The $p$-values are available in Supplementary Tab. 2.

| **F1-score** | rings - MICRA-Net | rings - U-Net | rings - Mask R-CNN | rings - ilastik | fibers - MICRA-Net | fibers - U-Net | fibers - Mask R-CNN | fibers - ilastik |
|---|---|---|---|---|---|---|---|---|
| rings - MICRA-Net | - | $6.9120 \times 10^{-1}$ | $1.0400 \times 10^{-2}$ | $<1.0000 \times 10^{-4}$ | $3.2890 \times 10^{-1}$ | $9.4110 \times 10^{-1}$ | $1.7770 \times 10^{-1}$ | $<1.0000 \times 10^{-4}$ |
| rings - U-Net | $6.9120 \times 10^{-1}$ | - | $2.7300 \times 10^{-2}$ | $<1.0000 \times 10^{-4}$ | $8.1230 \times 10^{-1}$ | $7.1990 \times 10^{-1}$ | $3.8030 \times 10^{-1}$ | $<1.0000 \times 10^{-4}$ |
| rings - Mask R-CNN | $1.0400 \times 10^{-2}$ | $2.7300 \times 10^{-2}$ | - | $<1.0000 \times 10^{-4}$ | $2.2800 \times 10^{-2}$ | $1.0900 \times 10^{-2}$ | $1.1970 \times 10^{-1}$ | $<1.0000 \times 10^{-4}$ |
| rings - ilastik | $<1.0000 \times 10^{-4}$ | $<1.0000 \times 10^{-4}$ | $<1.0000 \times 10^{-4}$ | - | $<1.0000 \times 10^{-4}$ | $<1.0000 \times 10^{-4}$ | $<1.0000 \times 10^{-4}$ | $1.0000 \times 10^{-4}$ |
| fibers - MICRA-Net | $3.2890 \times 10^{-1}$ | $8.1230 \times 10^{-1}$ | $2.2800 \times 10^{-2}$ | $<1.0000 \times 10^{-4}$ | - | $3.6560 \times 10^{-1}$ | $4.1000 \times 10^{-1}$ | $<1.0000 \times 10^{-4}$ |
| fibers - U-Net | $9.4110 \times 10^{-1}$ | $7.1990 \times 10^{-1}$ | $1.0900 \times 10^{-2}$ | $<1.0000 \times 10^{-4}$ | $3.6560 \times 10^{-1}$ | - | $1.7220 \times 10^{-1}$ | $<1.0000 \times 10^{-4}$ |
| fibers - Mask R-CNN | $1.7770 \times 10^{-1}$ | $3.8030 \times 10^{-1}$ | $1.1970 \times 10^{-1}$ | $<1.0000 \times 10^{-4}$ | $4.1000 \times 10^{-1}$ | $1.7220 \times 10^{-1}$ | - | $<1.0000 \times 10^{-4}$ |
| fibers - ilastik | $<1.0000 \times 10^{-4}$ | $<1.0000 \times 10^{-4}$ | $<1.0000 \times 10^{-4}$ | $1.0000 \times 10^{-4}$ | $<1.0000 \times 10^{-4}$ | $<1.0000 \times 10^{-4}$ | $<1.0000 \times 10^{-4}$ | - |

| **IOU** | rings - MICRA-Net | rings - U-Net | rings - Mask R-CNN | rings - ilastik | fibers - MICRA-Net | fibers - U-Net | fibers - Mask R-CNN | fibers - ilastik |
|---|---|---|---|---|---|---|---|---|
| rings - MICRA-Net | - | $8.8450 \times 10^{-1}$ | $5.2700 \times 10^{-2}$ | $<1.0000 \times 10^{-4}$ | $3.0140 \times 10^{-1}$ | $9.1990 \times 10^{-1}$ | $2.0100 \times 10^{-1}$ | $<1.0000 \times 10^{-4}$ |
| rings - U-Net | $8.8450 \times 10^{-1}$ | - | $8.8500 \times 10^{-2}$ | $<1.0000 \times 10^{-4}$ | $5.4080 \times 10^{-1}$ | $9.3880 \times 10^{-1}$ | $3.4030 \times 10^{-1}$ | $<1.0000 \times 10^{-4}$ |
| rings - Mask R-CNN | $5.2700 \times 10^{-2}$ | $8.8500 \times 10^{-2}$ | - | $<1.0000 \times 10^{-4}$ | $1.3220 \times 10^{-1}$ | $5.9700 \times 10^{-2}$ | $2.7520 \times 10^{-1}$ | $<1.0000 \times 10^{-4}$ |
| rings - ilastik | $<1.0000 \times 10^{-4}$ | $<1.0000 \times 10^{-4}$ | $<1.0000 \times 10^{-4}$ | - | $<1.0000 \times 10^{-4}$ | $<1.0000 \times 10^{-4}$ | $<1.0000 \times 10^{-4}$ | $1.0000 \times 10^{-4}$ |
| fibers - MICRA-Net | $3.0140 \times 10^{-1}$ | $5.4080 \times 10^{-1}$ | $1.3220 \times 10^{-1}$ | $<1.0000 \times 10^{-4}$ | - | $3.2490 \times 10^{-1}$ | $5.6470 \times 10^{-1}$ | $<1.0000 \times 10^{-4}$ |
| fibers - U-Net | $9.1990 \times 10^{-1}$ | $9.3880 \times 10^{-1}$ | $5.9700 \times 10^{-2}$ | $<1.0000 \times 10^{-4}$ | $3.2490 \times 10^{-1}$ | - | $2.1990 \times 10^{-1}$ | $<1.0000 \times 10^{-4}$ |
| fibers - Mask R-CNN | $2.0100 \times 10^{-1}$ | $3.4030 \times 10^{-1}$ | $2.7520 \times 10^{-1}$ | $<1.0000 \times 10^{-4}$ | $5.6470 \times 10^{-1}$ | $2.1990 \times 10^{-1}$ | - | $<1.0000 \times 10^{-4}$ |
| fibers - ilastik | $<1.0000 \times 10^{-4}$ | $<1.0000 \times 10^{-4}$ | $<1.0000 \times 10^{-4}$ | $1.0000 \times 10^{-4}$ | $<1.0000 \times 10^{-4}$ | $<1.0000 \times 10^{-4}$ | $<1.0000 \times 10^{-4}$ | - |

| **SBD** | rings - MICRA-Net | rings - U-Net | rings - Mask R-CNN | rings - ilastik | fibers - MICRA-Net | fibers - U-Net | fibers - Mask R-CNN | fibers - ilastik |
|---|---|---|---|---|---|---|---|---|
| rings - MICRA-Net | - | $6.4580 \times 10^{-1}$ | $9.0000 \times 10^{-3}$ | $<1.0000 \times 10^{-4}$ | $1.7100 \times 10^{-2}$ | $1.0000 \times 10^{-4}$ | $1.7760 \times 10^{-1}$ | $<1.0000 \times 10^{-4}$ |
| rings - U-Net | $6.4580 \times 10^{-1}$ | - | $9.1000 \times 10^{-3}$ | $<1.0000 \times 10^{-4}$ | $2.1140 \times 10^{-1}$ | $2.2700 \times 10^{-2}$ | $3.9660 \times 10^{-1}$ | $<1.0000 \times 10^{-4}$ |
| rings - Mask R-CNN | $9.0000 \times 10^{-3}$ | $9.1000 \times 10^{-3}$ | - | $3.0000 \times 10^{-4}$ | $8.0000 \times 10^{-4}$ | $1.0000 \times 10^{-4}$ | $2.6000 \times 10^{-3}$ | $6.3000 \times 10^{-3}$ |
| rings - ilastik | $<1.0000 \times 10^{-4}$ | $<1.0000 \times 10^{-4}$ | $3.0000 \times 10^{-4}$ | - | $<1.0000 \times 10^{-4}$ | $<1.0000 \times 10^{-4}$ | $<1.0000 \times 10^{-4}$ | $1.2000 \times 10^{-3}$ |
| fibers - MICRA-Net | $1.7100 \times 10^{-2}$ | $2.1140 \times 10^{-1}$ | $8.0000 \times 10^{-4}$ | $<1.0000 \times 10^{-4}$ | - | $1.3590 \times 10^{-1}$ | $9.8260 \times 10^{-1}$ | $<1.0000 \times 10^{-4}$ |
| fibers - U-Net | $1.0000 \times 10^{-4}$ | $2.2700 \times 10^{-2}$ | $1.0000 \times 10^{-4}$ | $<1.0000 \times 10^{-4}$ | $1.3590 \times 10^{-1}$ | - | $3.4490 \times 10^{-1}$ | $<1.0000 \times 10^{-4}$ |
| fibers - Mask R-CNN | $1.7760 \times 10^{-1}$ | $3.9660 \times 10^{-1}$ | $2.6000 \times 10^{-3}$ | $<1.0000 \times 10^{-4}$ | $9.8260 \times 10^{-1}$ | $3.4490 \times 10^{-1}$ | - | $<1.0000 \times 10^{-4}$ |
| fibers - ilastik | $<1.0000 \times 10^{-4}$ | $<1.0000 \times 10^{-4}$ | $6.3000 \times 10^{-3}$ | $1.2000 \times 10^{-3}$ | $<1.0000 \times 10^{-4}$ | $<1.0000 \times 10^{-4}$ | $<1.0000 \times 10^{-4}$ | - |

Supplementary Table 2: Comparison of the performance metrics (F1-score, IOU, and SBD) for MICRA-Net, weakly-supervised U-Net, weakly-supervised Mask R-CNN and Ilastik segmentation on the F-actin dataset. The $p$-values are obtained by a permutation test with 10 000 repetitions comparing each line against each columns following a resampled ANOVA with 10 000 repetitions ($p_{\mathrm{F1-score}} =< 1.0000 \times 10^{-4}$, $p_{\mathrm{IOU}} =< 1.0000 \times 10^{-4}$ and $p_{\mathrm{SBD}} =< 1.0000 \times 10^{-4}$). Color code: increase (cyan), decrease (red), and no significant changes (black) of each line compared to each column of the metric scores.

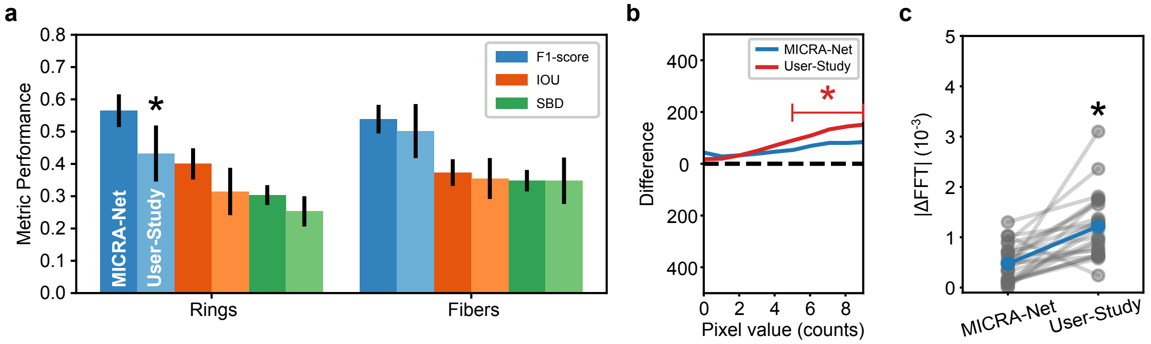


Supplementary Figure 11: Comparison of MICRA-Net with the User-Study on the segmentation of F-actin nanostructures. a) F1-Score (blue), IOU (orange), and SBD (green) are calculated using the segmentation masks from a precisely annotated testing set of 50 images as a ground truth. Bar graphs and error bars show the bootstrapped mean and 95% confidence interval (10 000 repetitions). The F1-score is significantly lower for the participants in the study compared with MICRA-Net when segmenting F-actin rings. All $p$-values are presented in Supplementary Tab. 3. b) The pixel intensity distribution metric for the User-Study was calculated as the average histogram of annotated pixels for the 6 participants. A statistical analysis revealed a significant difference in the number of low-intensity ($[5, 9]$) foreground pixels values for the User-Study when compared to the Expert annotations, while no difference is observed for MICRA-Net (Supplementary Tab. 5 for $p$-values). c) The FTT metric is significantly lower for MICRA-Net, implying less difference with precise Expert annotations, and therefore a more accurate segmentation (resampling test, 10 000 repetitions, $p =< 1.0000 \times 10^{-4}$).

| **F1-score** | MICRA-Net - rings | User-Study - rings | MICRA-Net - fibers | User-Study - fibers |
|---|---|---|---|---|
| MICRA-Net - rings | - | $6.2000 \times 10^{-3}$ | $3.3500 \times 10^{-1}$ | $1.7290 \times 10^{-1}$ |
| User-Study - rings | $6.2000 \times 10^{-3}$ | - | $1.2900 \times 10^{-2}$ | $1.9240 \times 10^{-1}$ |
| MICRA-Net - fibers | $3.3500 \times 10^{-1}$ | $1.2900 \times 10^{-2}$ | - | $3.7880 \times 10^{-1}$ |
| User-Study - fibers | $1.7290 \times 10^{-1}$ | $1.9240 \times 10^{-1}$ | $3.7880 \times 10^{-1}$ | - |

| **SBD** | MICRA-Net - rings | User-Study - rings | MICRA-Net - fibers | User-Study - fibers |
|---|---|---|---|---|
| MICRA-Net - rings | - | $6.8300 \times 10^{-2}$ | $1.9000 \times 10^{-2}$ | $1.3990 \times 10^{-1}$ |
| User-Study - rings | $6.8300 \times 10^{-2}$ | - | $2.4000 \times 10^{-3}$ | $3.3900 \times 10^{-2}$ |
| MICRA-Net - fibers | $1.9000 \times 10^{-2}$ | $2.4000 \times 10^{-3}$ | - | $9.9520 \times 10^{-1}$ |
| User-Study - fibers | $1.3990 \times 10^{-1}$ | $3.3900 \times 10^{-2}$ | $9.9520 \times 10^{-1}$ | - |

Supplementary Table 3: Comparison of the performance metrics (F1-score and SBD) for MICRA-Net and the User-Study. No significant changes are measured for the IOU. The $p$-values are obtained by a permutation test with 10 000 repetitions comparing each line against each column following a resampled ANOVA test with 10 000 repetitions ($p_{\text{F1-score}} = 0.0214$, $p_{\text{IOU}} = 0.1697$ and $p_{SBD} = 0.0380$). Color code : increase (cyan), decrease (red), and no significant changes (black) of each line compared to each column of the metric scores.

| **FFT** | Expert | MICRA-Net | U-Net | Mask R-CNN | Ilastik |
|---|---|---|---|---|---|
| Expert | - | $6.4470 \times 10^{-1}$ | $2.4000 \times 10^{-3}$ | $4.8000 \times 10^{-3}$ | $<1.0000 \times 10^{-4}$ |
| MICRA-Net | $6.4470 \times 10^{-1}$ | - | $1.0500 \times 10^{-2}$ | $4.3000 \times 10^{-3}$ | $<1.0000 \times 10^{-4}$ |
| U-Net | $2.4000 \times 10^{-3}$ | $1.0500 \times 10^{-2}$ | - | $<1.0000 \times 10^{-4}$ | $<1.0000 \times 10^{-4}$ |
| Mask R-CNN | $4.8000 \times 10^{-3}$ | $4.3000 \times 10^{-3}$ | $<1.0000 \times 10^{-4}$ | - | $1.6000 \times 10^{-1}$ |
| Ilastik | $<1.0000 \times 10^{-4}$ | $<1.0000 \times 10^{-4}$ | $<1.0000 \times 10^{-4}$ | $1.6000 \times 10^{-1}$ | - |

Supplementary Table 4: Comparison of the FFT metrics for MICRA-Net, weakly-supervised U-Net, weakly-supervised Mask R-CNN and Ilastik segmentation on the F-actin dataset for the periodical lattice. The $p$-values are obtained by a permutation test with 10 000 repetitions comparing each lines against each columns following a resampled ANOVA with 10 000 repetitions ($p =< 1.0000 \times 10^{-4}$). Color code : increase (cyan), decrease (red), and no significant changes (black) of each line compared to each column of the metric scores.

| Pixel value | 0 | 1 | 2 | 3 | 4 |
|---|---|---|---|---|---|
| MICRA-Net | $1.2810 \times 10^{-1}$ | $1.8690 \times 10^{-1}$ | $1.6790 \times 10^{-1}$ | $1.7430 \times 10^{-1}$ | $1.6920 \times 10^{-1}$ |
| U-Net | $2.8220 \times 10^{-1}$ | $9.3080 \times 10^{-1}$ | $2.9640 \times 10^{-1}$ | $1.0050 \times 10^{-1}$ | $\mathbf{3.6300 \times 10^{-2}}$ |
| Mask R-CNN | $5.3400 \times 10^{-2}$ | $3.6240 \times 10^{-1}$ | $9.8270 \times 10^{-1}$ | $3.9320 \times 10^{-1}$ | $2.0380 \times 10^{-1}$ |
| Ilastik | $\mathbf{<1.0000 \times 10^{-4}}$ | $\mathbf{<1.0000 \times 10^{-4}}$ | $\mathbf{<1.0000 \times 10^{-4}}$ | $\mathbf{<1.0000 \times 10^{-4}}$ | $\mathbf{<1.0000 \times 10^{-4}}$ |
| User-Study | $3.5620 \times 10^{-1}$ | $3.0490 \times 10^{-1}$ | $1.7940 \times 10^{-1}$ | $1.0930 \times 10^{-1}$ | $6.7400 \times 10^{-2}$ |

| Pixel value | 5 | 6 | 7 | 8 | 9 |
|---|---|---|---|---|---|
| MICRA-Net | $1.6060 \times 10^{-1}$ | $9.8200 \times 10^{-2}$ | $6.4300 \times 10^{-2}$ | $9.2400 \times 10^{-2}$ | $6.6900 \times 10^{-2}$ |
| U-Net | $\mathbf{1.6600 \times 10^{-2}}$ | $\mathbf{9.2000 \times 10^{-3}}$ | $\mathbf{3.9000 \times 10^{-3}}$ | $\mathbf{4.2000 \times 10^{-3}}$ | $\mathbf{2.8000 \times 10^{-3}}$ |
| Mask R-CNN | $9.7700 \times 10^{-2}$ | $5.1200 \times 10^{-2}$ | $\mathbf{2.4900 \times 10^{-2}}$ | $\mathbf{2.4800 \times 10^{-2}}$ | $\mathbf{1.4700 \times 10^{-2}}$ |
| Ilastik | $\mathbf{<1.0000 \times 10^{-4}}$ | $\mathbf{<1.0000 \times 10^{-4}}$ | $\mathbf{<1.0000 \times 10^{-4}}$ | $\mathbf{<1.0000 \times 10^{-4}}$ | $\mathbf{<1.0000 \times 10^{-4}}$ |
| User-Study | $\mathbf{4.5200 \times 10^{-2}}$ | $\mathbf{2.3000 \times 10^{-2}}$ | $\mathbf{1.4800 \times 10^{-2}}$ | $\mathbf{2.1100 \times 10^{-2}}$ | $\mathbf{1.6500 \times 10^{-2}}$ |

Supplementary Table 5: Comparison of the Expert annotations distribution of intensity and the predicted segmentation masks for the fiber metric. Only low intensity pixels (0-9) are shown (see Methods). The $p$-values are obtained from a permutation test of 10 000 permutations comparing the distribution of pixel counts in a given intensity bin. A significantly different value is highlighted in bold.

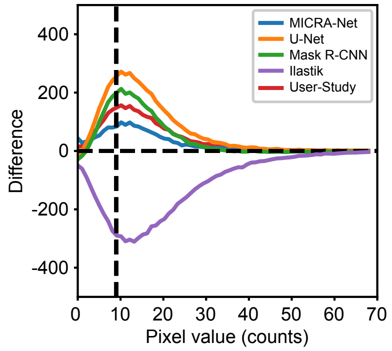


Supplementary Figure 12: The average raw difference from the Expert annotations in the number of annotated pixels is shown over the complete range of pixel values for all trained models on all 25 testing images (fibers only). Closer to zero is better. The vertical black dashed line represent the threshold of low intensity pixels which are shown in Fig 3d and Supplementary Fig. 11b.

| Cell line | Baselines | MICRA-Net |
|---|---|---|
| DIC-C2DH-HeLa | 0.38 | 1.00 |
| Fluo-C2DL-MSC | 0.79 | 1.28 |
| Fluo-N2DH-GOWT1 | 0.48 | 1.00 |
| Fluo-N2DL-HeLa | 1.29 | 2.15 |
| PhC-C2DH-U373 | 1.30 | 1.30 |
| PhC-C2DH-PSC | 3.20 | 3.20 |

Supplementary Table 6: Scale factors used to resize the Cell Tracking Challenge cell line images for baselines and MICRA-Net training.

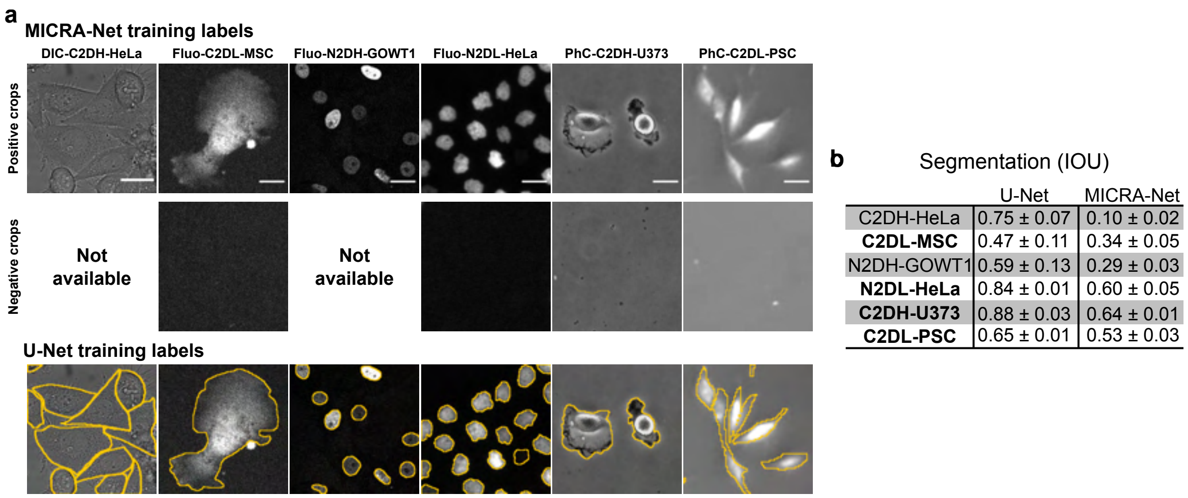


| | U-Net | MICRA-Net |
|---|---|---|
| C2DH-HeLa | 0.75 ± 0.07 | 0.10 ± 0.02 |
| **C2DL-MSC** | 0.47 ± 0.11 | 0.34 ± 0.05 |
| N2DH-GOWT1 | 0.59 ± 0.13 | 0.29 ± 0.03 |
| **N2DL-HeLa** | 0.84 ± 0.01 | 0.60 ± 0.05 |
| **C2DH-U373** | 0.88 ± 0.03 | 0.64 ± 0.01 |
| **C2DL-PSC** | 0.65 ± 0.01 | 0.53 ± 0.03 |

Supplementary Figure 13: Segmentation comparison of MICRA-Net and U-Net using the same resampling method of images as Falk et al. [3] on the Cell Tracking Challenge. While MICRA-Net obtained a high classification accuracy of $(95.8 \pm 0.4)\,\%$ on the testing set, it minimally requires examples of negative crops to be able to extract enough context to differentiate between the cells and background. This is demonstrated by the poor segmentation performance on DIC-C2DH-HeLa and Fluo-N2DH-GOWT1 cell lines when no negative crops are extracted.

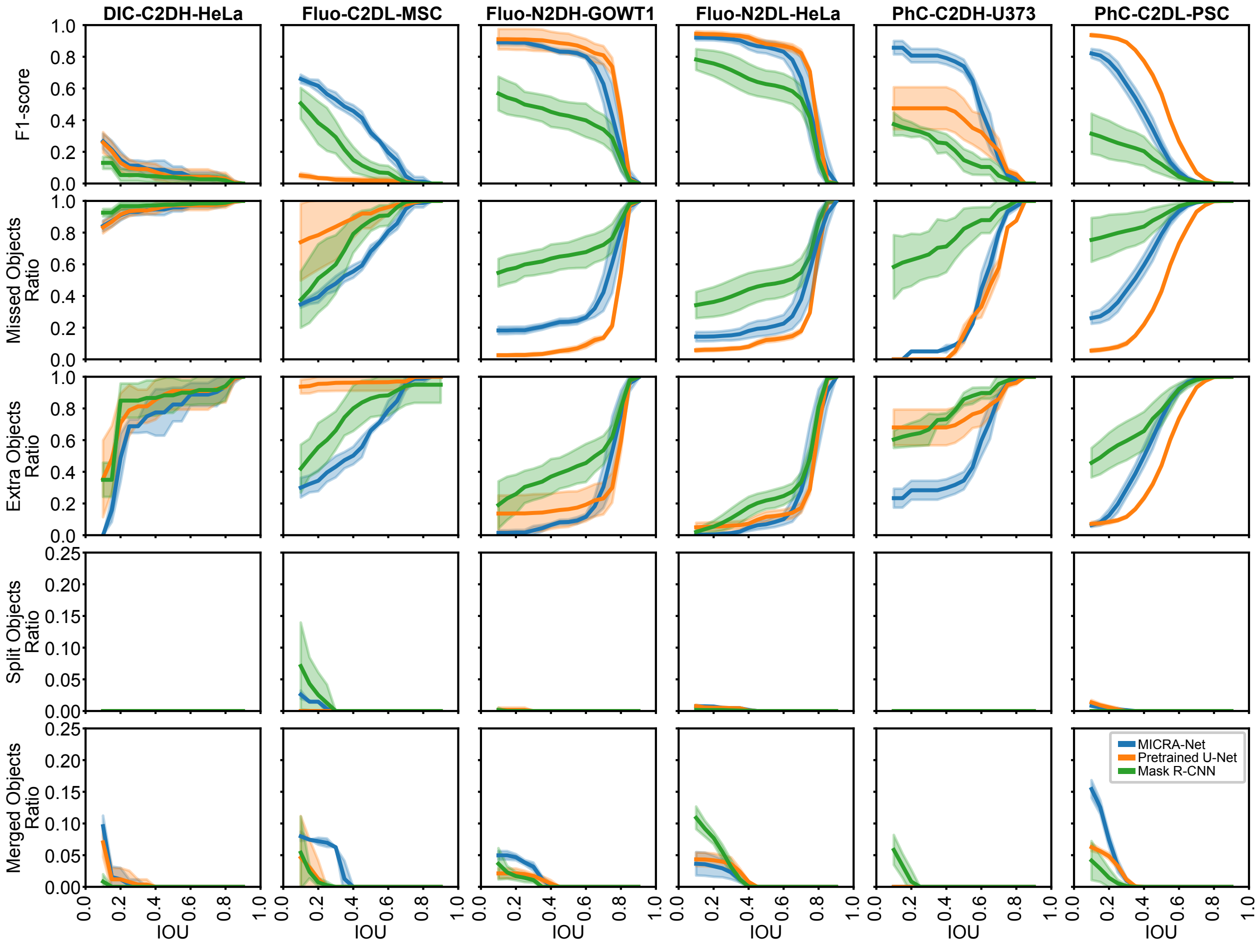


Supplementary Figure 14: Comparison of MICRA-Net with baselines (U-Net and Mask R-CNN) on the Cell Tracking Challenge using different metrics (F1-score, missed objects ratio, extra objects ratio, split objects ratio, merged objects ratio) as a function of the IOU. Baselines are trained using *bounding boxes* at the same scale as MICRA-Net. The performance on the DIC-C2DH-HeLa cell line is very low for all methods since at this scale no complete cells can be resolved within an extracted crop.

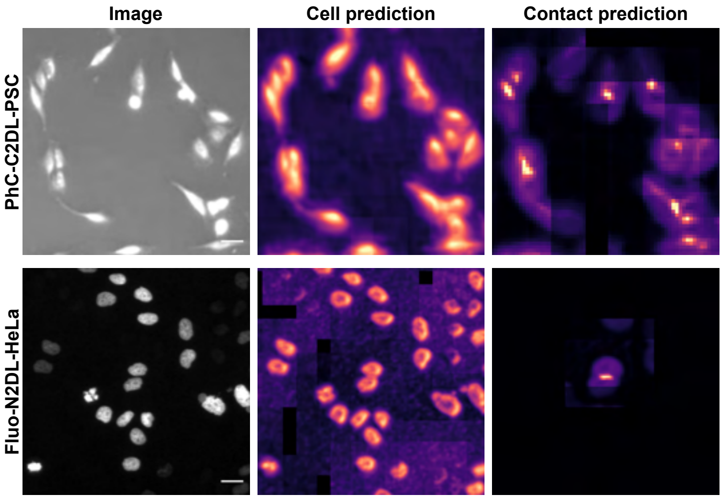


Supplementary Figure 15: Representative examples of the instance segmentation procedure using MICRA-Net for two cell lines of the Cell Tracking Challenge (top: PhC-C2DL-PSC, bottom: Fluo-N2DL-HeLa). Shown are the input image (*left*), the PCA decomposition of the raw feature maps extracted from layers $L^{1-7}$ of MICRA-Net for the cell prediction (*middle*, and the grad-CAM of layer $L^8$ for semantic contact (*right*). Scale bars: 25 µm.

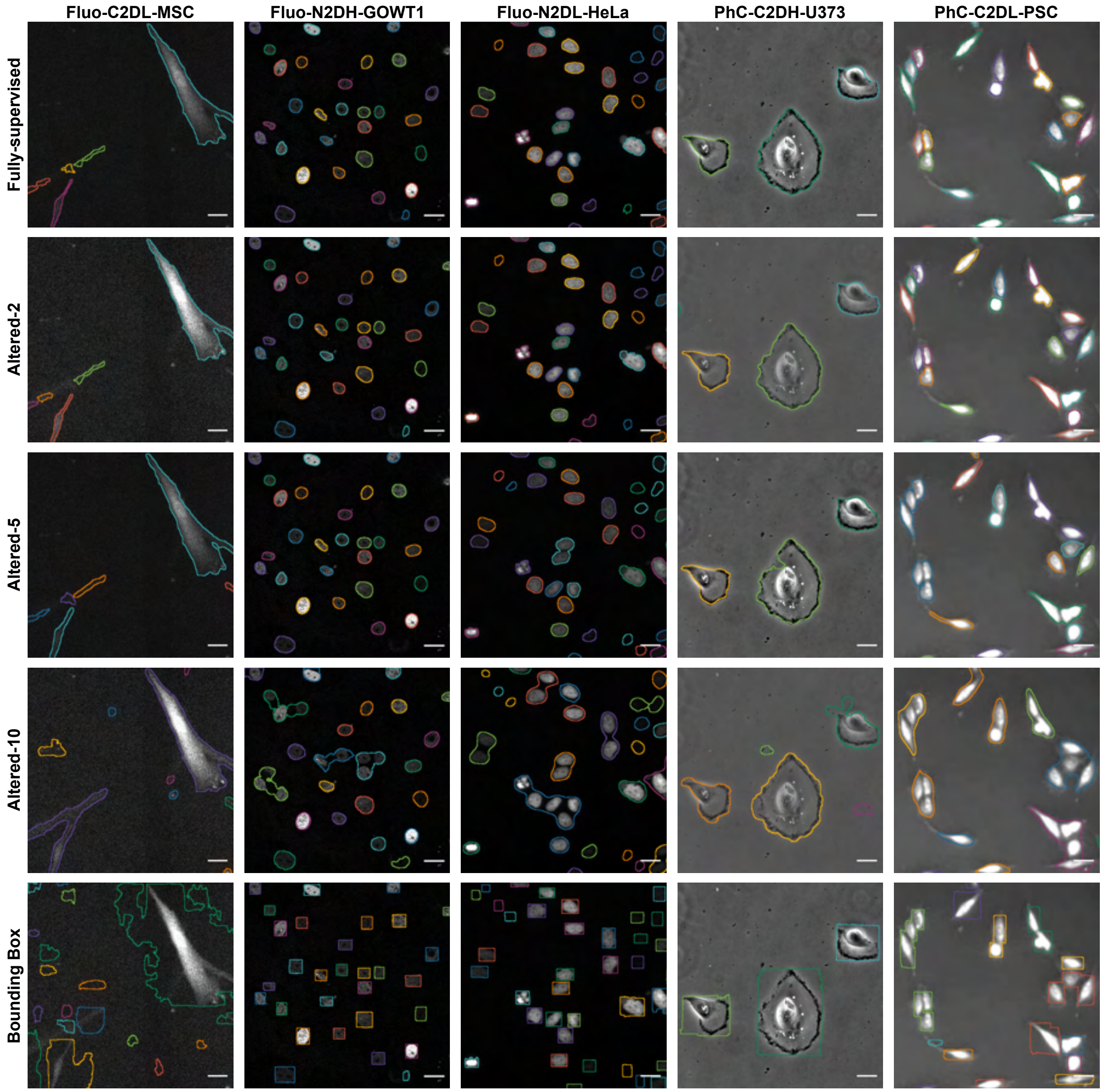


Supplementary Figure 16: Representative examples of the U-Net segmentation for each evaluated cell line of the Cell Tracking Challenge. Each row represents a different level of supervision (Fully-supervised, altered-2, altered-5, altered-10 and bounding boxes). Distinct cells are labeled using different colors. Scale bars: 25 µm.

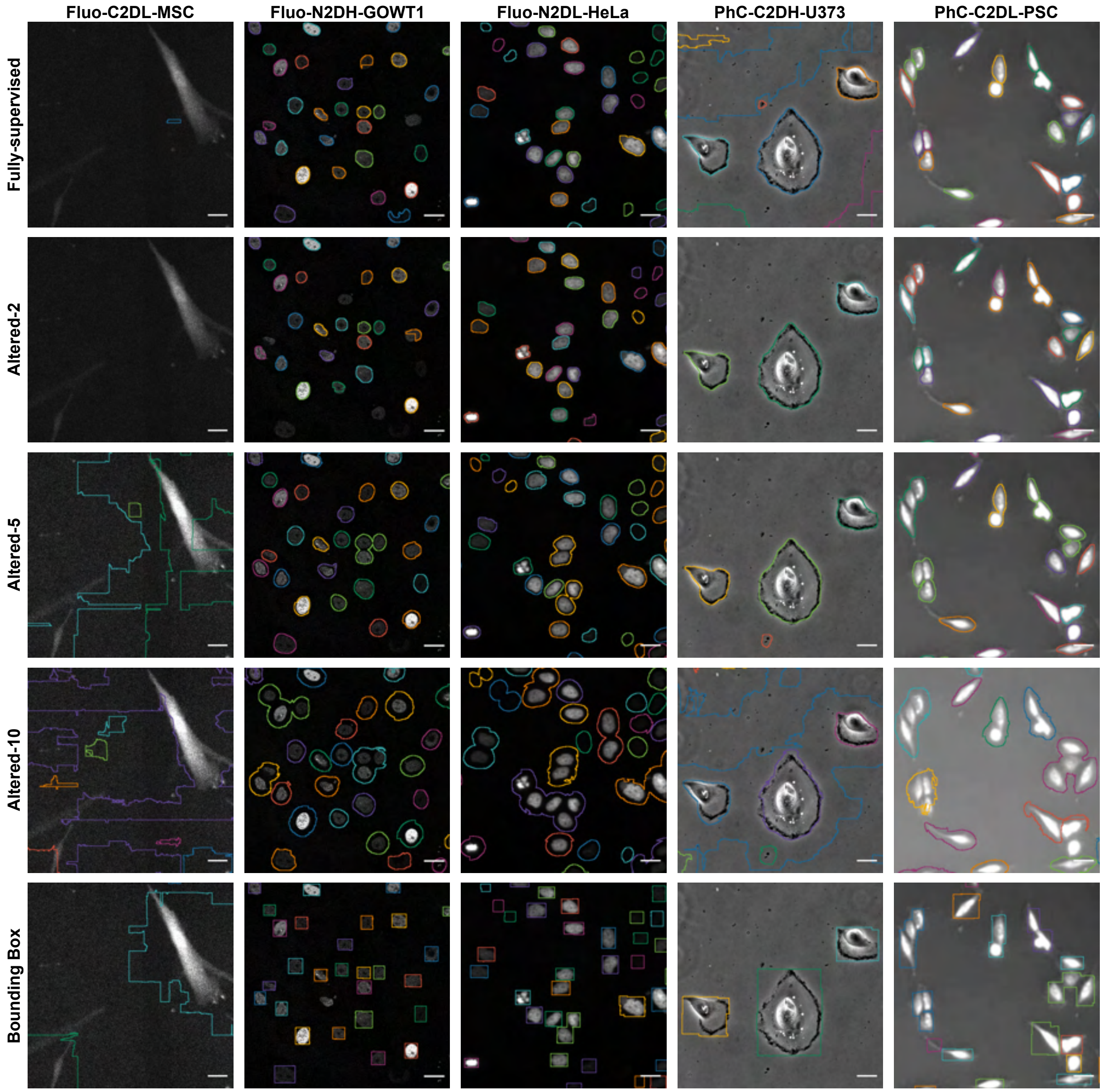


Supplementary Figure 17: Representative examples of the Mask R-CNN segmentation for each evaluated cell line of the Cell Tracking Challenge. Each row represents a different level of supervision (Fully-supervised, altered-2, altered-5, altered-10 and bounding boxes). Distinct cells are labeled using different colors. Scale bars: 25 µm.

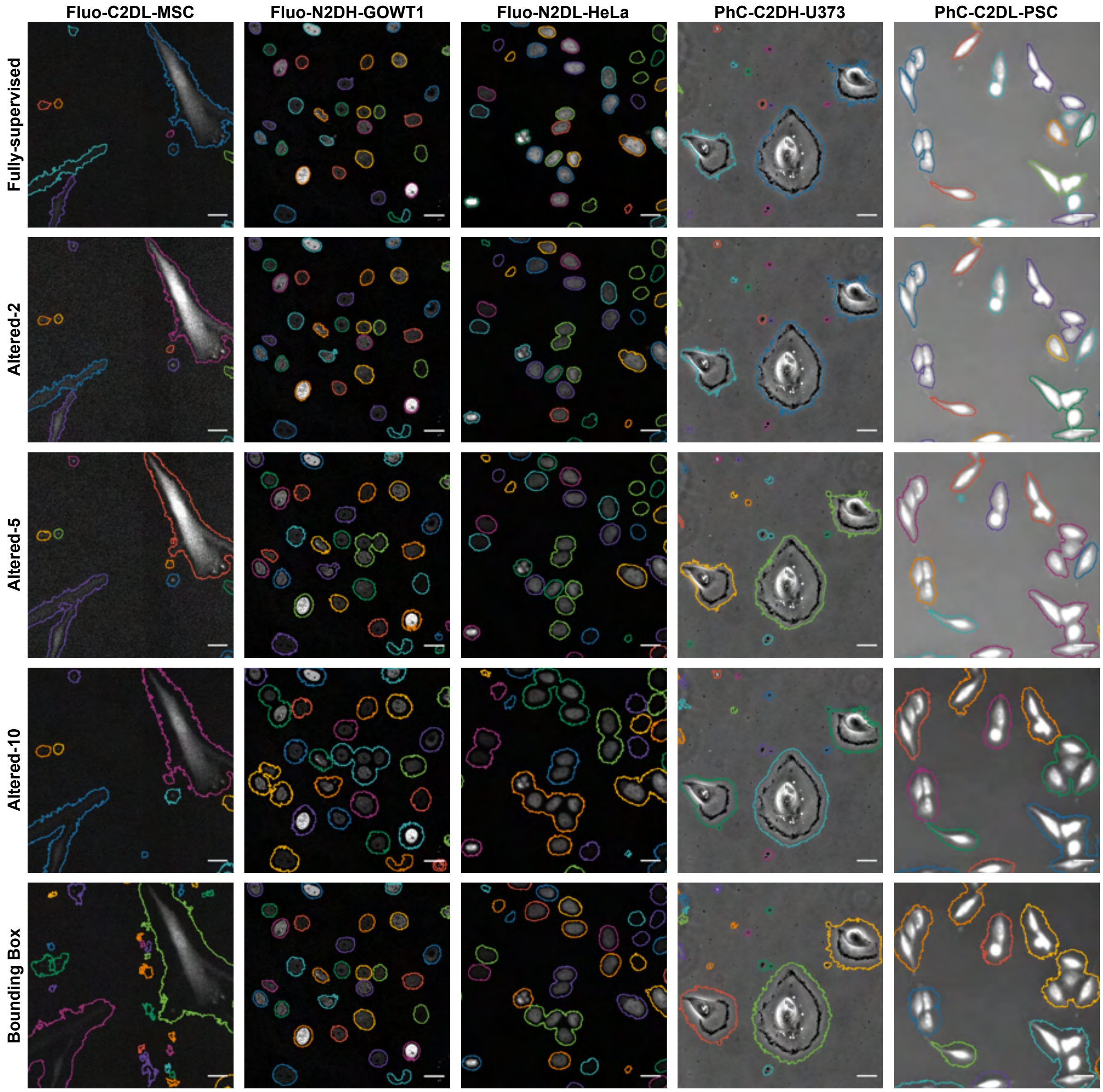


Supplementary Figure 18: Representative examples of the Ilastik segmentation for each evaluated cell line of the Cell Tracking Challenge. Each row represents a different level of supervision (Fully-supervised, altered-2, altered-5, altered-10 and bounding boxes). Distinct cells are labeled using different colors. Scale bars: 25 µm.

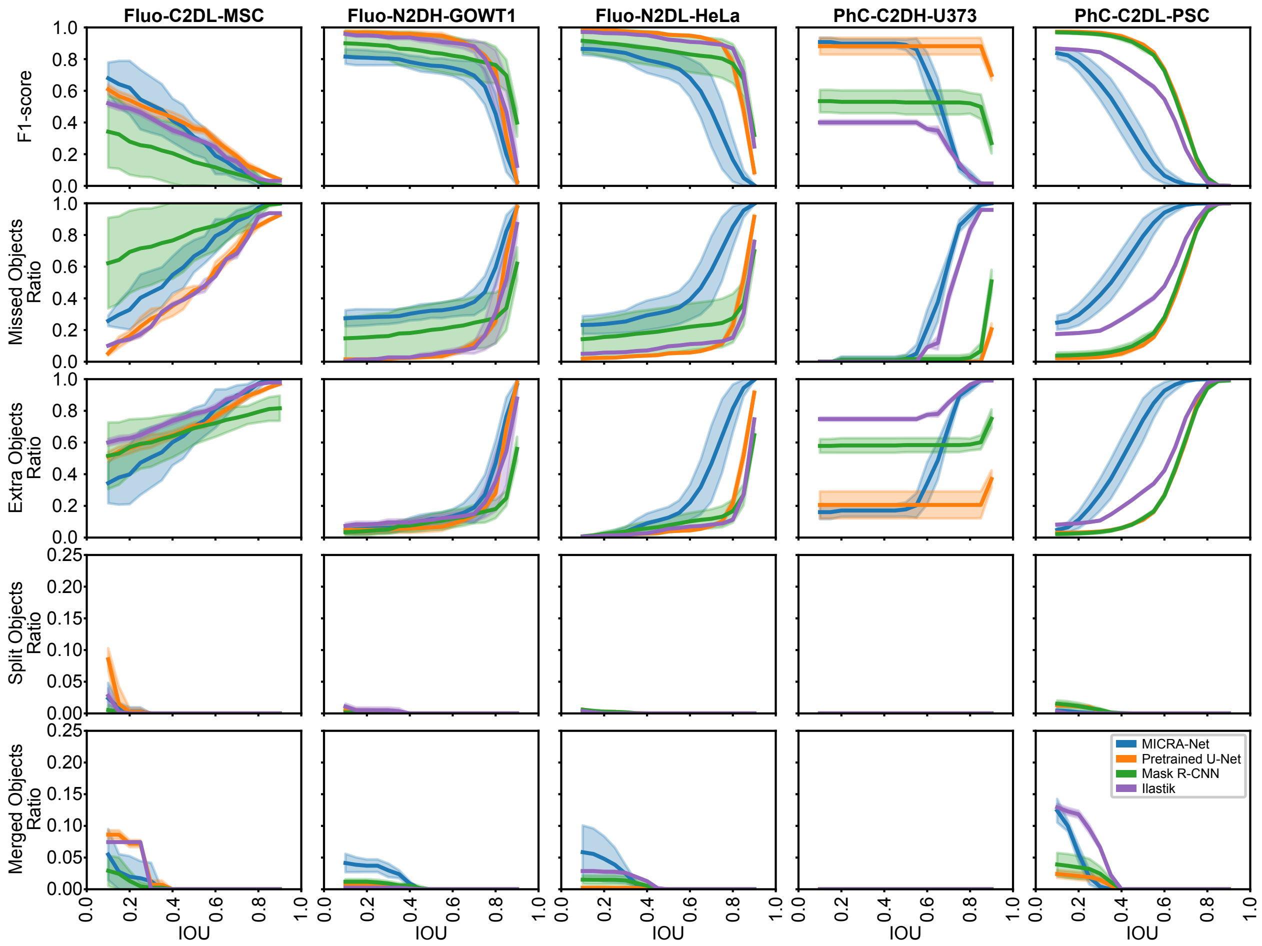


Supplementary Figure 19: Comparison of MICRA-Net with baselines (U-Net, Mask R-CNN, and Ilastik) on the Cell Tracking Challenge using different metrics (F1-score, missed objects ratio, extra objects ratio, split objects ratio, merged objects ratio) as a function of the IOU. Baselines are trained using the *fully-supervised* dataset.

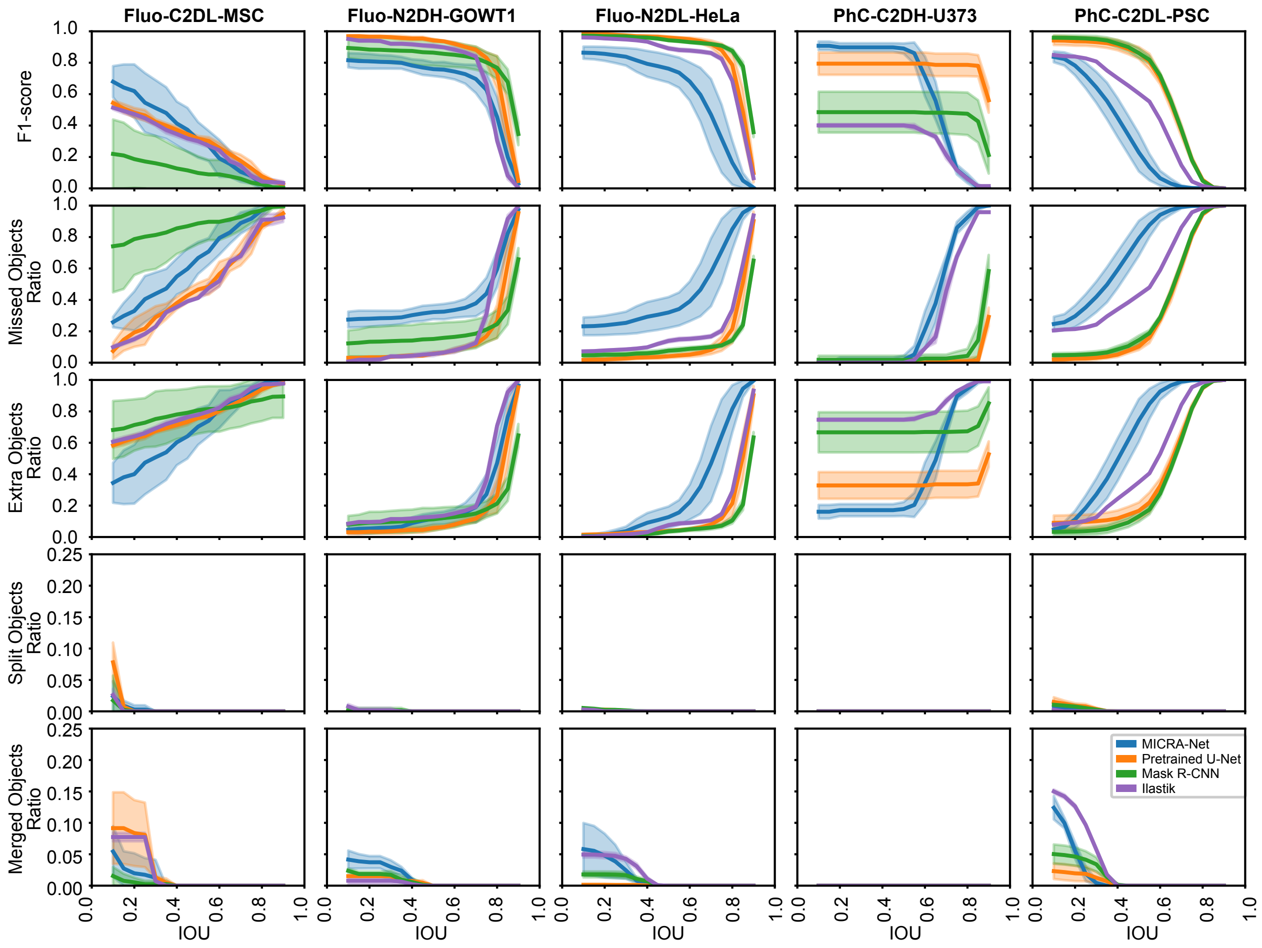


Supplementary Figure 20: Comparison of MICRA-Net with baselines (U-Net, Mask R-CNN, and Ilastik) on the Cell Tracking Challenge for different metrics (F1-score, missed objects ratio, extra objects ratio, split objects ratio, merged objects ratio) as a function of the IOU. Baselines are trained using an *altered* version of the dataset. The datasets were altered (dilation or erosion) using a disk structuring element of random radius size to mimic non-constant annotation by an Expert. The radius size was sampled from normal distribution with a mean of 0 and a standard deviation of 2 (ALT-2). A negative radius value implied an erosion, while a positive value resulted in a dilation.

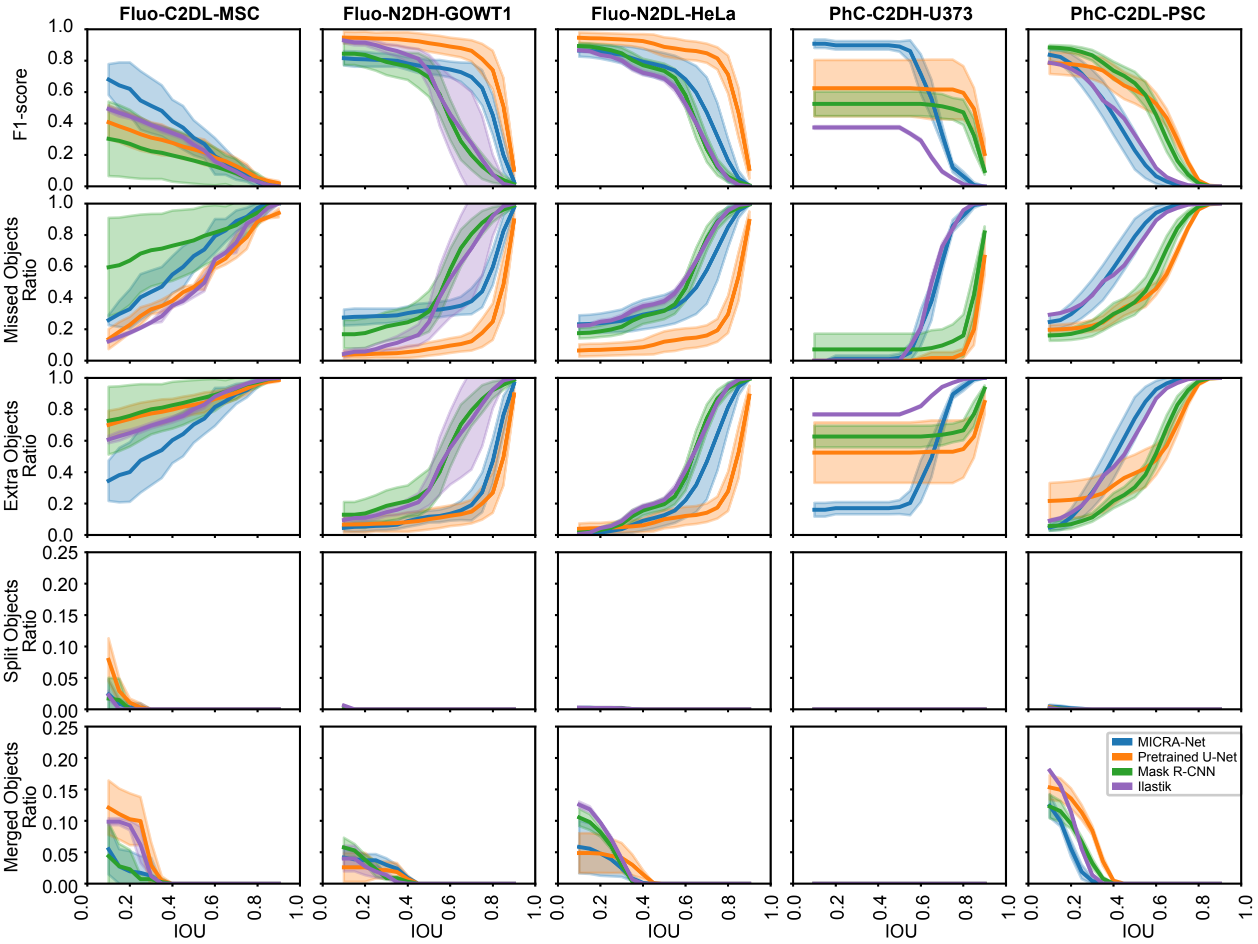


Supplementary Figure 21: Comparison of MICRA-Net with baselines (U-Net, Mask R-CNN, and Ilastik) on the Cell Tracking Challenge for different metrics (F1-score, missed objects ratio, extra objects ratio, split objects ratio, merged objects ratio) as a function of the IOU. Baselines are trained using an *altered* version of the dataset. The datasets were altered (dilation or erosion) using a disk structuring element of random radius size to mimic non-constant annotation by an Expert. The radius size was sampled from normal distribution with a mean of 0 and a standard deviation of 5 (ALT-5). A negative radius value implied an erosion, while a positive value resulted in a dilation.

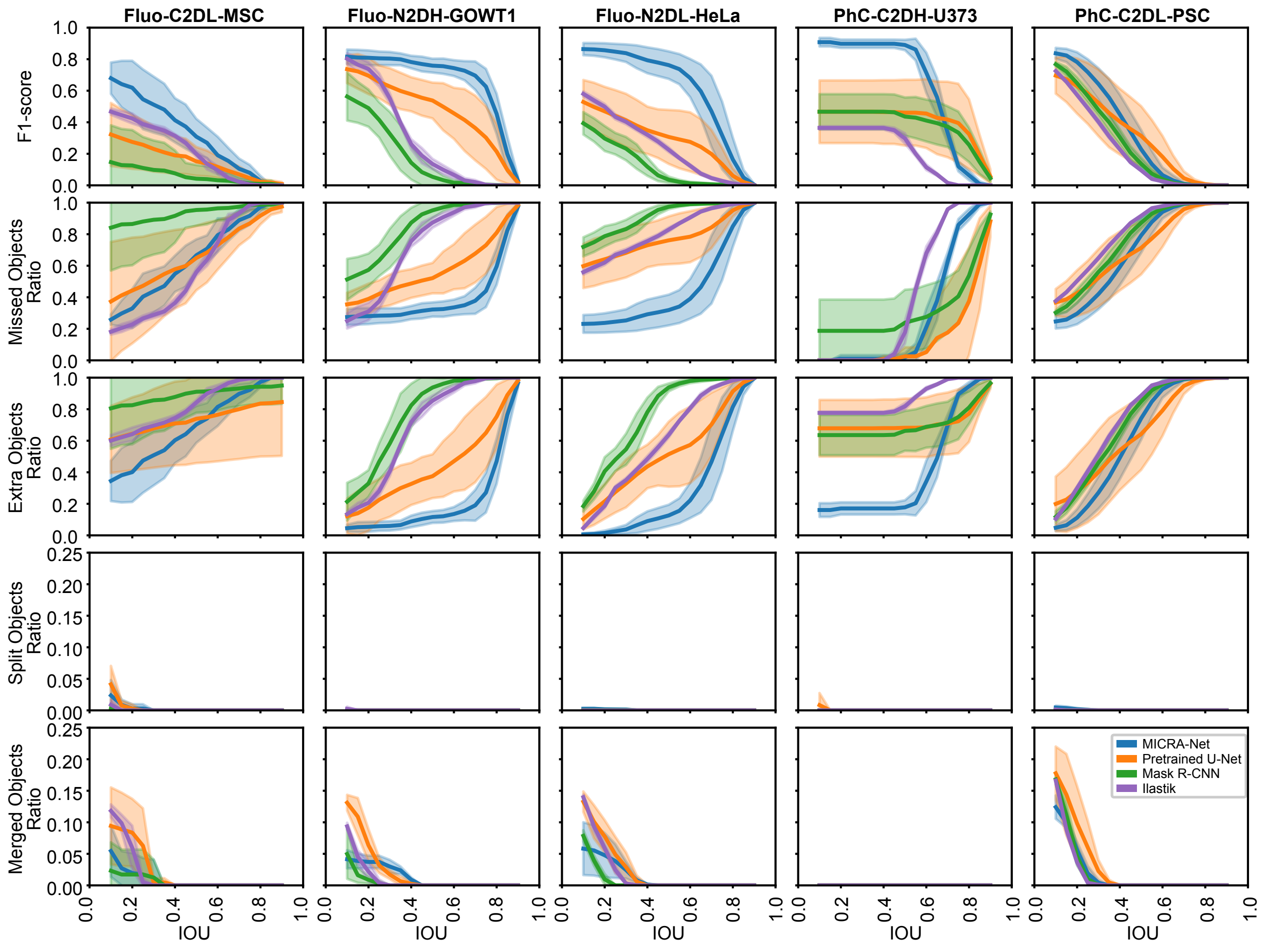


Supplementary Figure 22: Comparison of MICRA-Net with baselines (U-Net, Mask R-CNN, and Ilastik) on the Cell Tracking Challenge for different metrics (F1-score, missed objects ratio, extra objects ratio, split objects ratio, merged objects ratio) as a function of the IOU. Baselines are trained using an *altered* version of the dataset. The datasets were altered (dilation or erosion) using a disk structuring element of random radius size to mimic non-constant annotation by an Expert. The radius size was sampled from normal distribution with a mean of 0 and a standard deviation of 10 (ALT-10). A negative radius value implied an erosion, while a positive value resulted in a dilation.

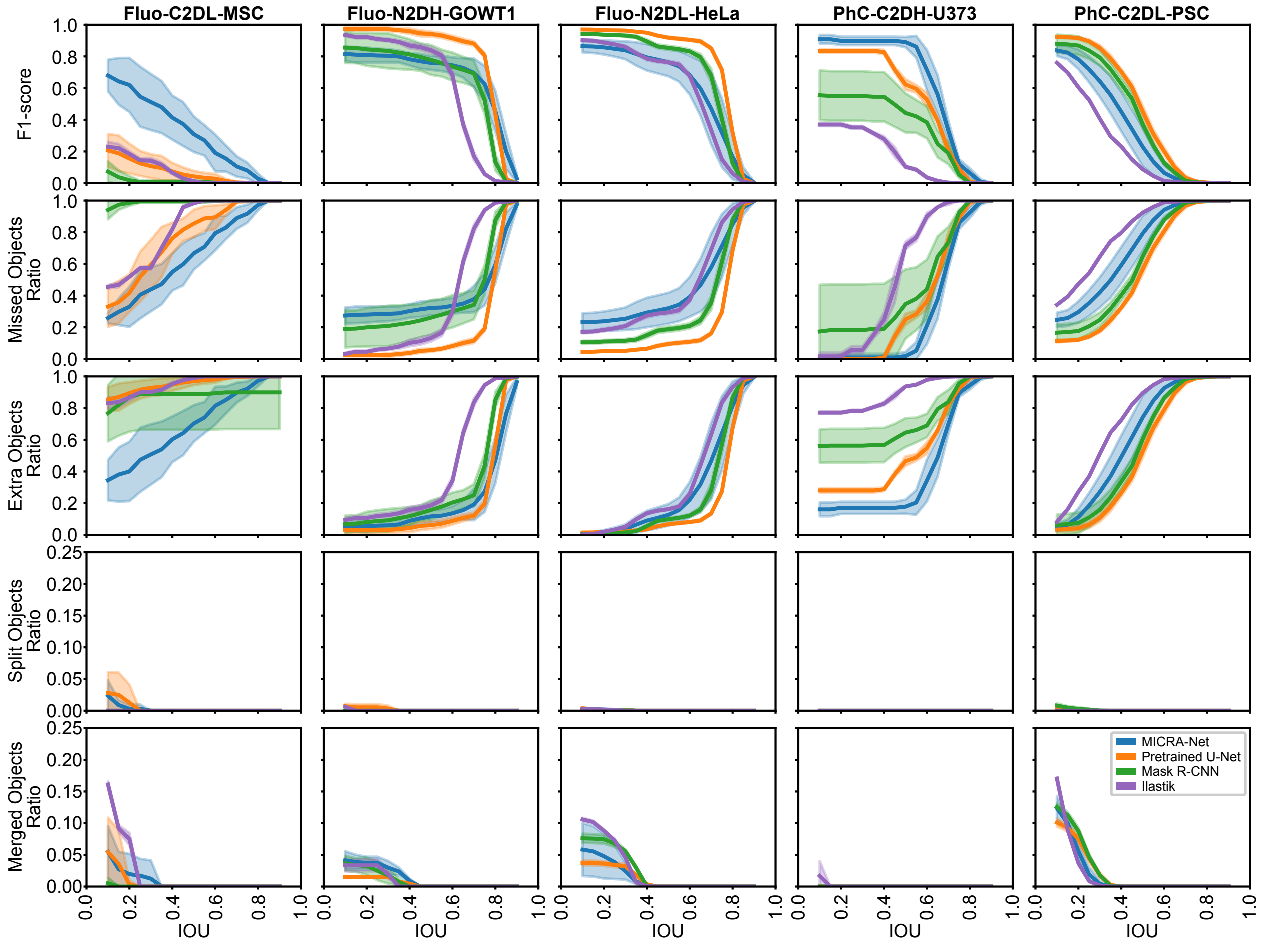


Supplementary Figure 23: Comparison of MICRA-Net with baselines (U-Net, Mask R-CNN, and Ilastik) on the Cell Tracking Challenge for different metrics (F1-score, missed objects ratio, extra objects ratio, split objects ratio, merged objects ratio) as a function of the IOU. The performance of MICRA-Net and the baselines (U-Net, Mask R-CNN, and Ilastik) trained using *bounding boxes* as targets are reported.

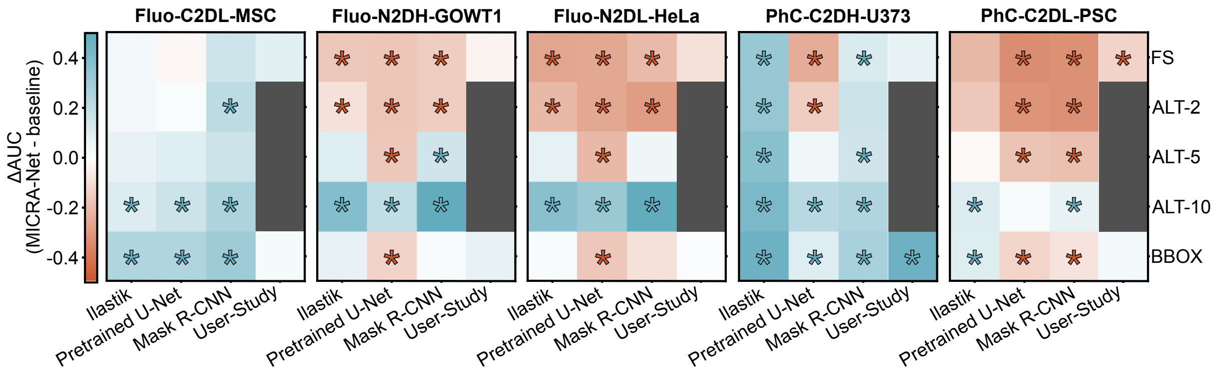


Supplementary Figure 24: Comparison of the area under the curve (AUC) of each cell line on the Cell Tracking Challenge for various level of supervisions (fully-supervised (FS), altered-2 (ALT-2), altered-5 (ALT-5), altered-10 (ALT-10), and bounding boxes (BBOX)). The difference measured for MICRA-Net with the trained baselines and the conducted User-Study is reported. The increase/decrease in performance of MICRA-Net compared with baselines varies for the different cell lines and depends on the level of supervision. An increase in performance is shown in blue and a decrease in red. A significant change is highlighted with a star (Supplementary Tab. 10,12,14, and 8).

| | FS | ALT-2 | ALT-5 | ALT-10 | BBOX |
|---|---|---|---|---|---|
| U-Net | $83.1 \pm 1.0$ | $77.1 \pm 3.1$ | $77.2 \pm 3.2$ | $78.4 \pm 1.9$ | $80.8 \pm 2.6$ |
| Mask R-CNN | $74.2 \pm 2.4$ | $66.1 \pm 4.2$ | $63.8 \pm 6.8$ | $62.9 \pm 12.4$ | $65.0 \pm 10.0$ |
| Ilastik | $49.1 \pm 0.3$ | $49.5 \pm 0.5$ | $49.9 \pm 0.7$ | $50.3 \pm 0.6$ | $51.0 \pm 1.0$ |
| MICRA-Net | $90.2 \pm 1.1$ | | | | |

Supplementary Table 7: Classification accuracy of the baselines on the Cell Tracking Challenge depending on the level of supervision (fully-supervised (FS), altered-2 (ALT-2), altered-5 (ALT-5), altered-10 (ALT-10), and bounding box (BBOX)). The classification accuracy is calculated from all extracted crops of the testing dataset. A crop is considered positive to a specific class if any pixels belong to this class. The classification accuracy of the cell line detection of MICRA-Net is reported for comparison. Shown is the bootstrapped mean and 95% confidence interval obtained from bootstrapping (10 000 repetitions).

| **PhC-C2DH-U373** | MICRA-Net | Precise | BBOX |
|---|---|---|---|
| MICRA-Net | - | $5.8780 \times 10^{-1}$ | $4.0000 \times 10^{-4}$ |
| Precise | $5.8780 \times 10^{-1}$ | - | $2.6000 \times 10^{-3}$ |
| BBOX | $4.0000 \times 10^{-4}$ | $2.6000 \times 10^{-3}$ | - |
| **PhC-C2DL-PSC** | MICRA-Net | Precise | BBOX |
| MICRA-Net | - | $2.5200 \times 10^{-2}$ | $2.4880 \times 10^{-1}$ |
| Precise | $2.5200 \times 10^{-2}$ | - | $1.7000 \times 10^{-3}$ |
| BBOX | $2.4880 \times 10^{-1}$ | $1.7000 \times 10^{-3}$ | - |

Supplementary Table 8: Statistical analysis comparing the normalized area under the curve of MICRA-Net, fully- and weakly-supervised annotations from the User-Study for each cell line of the Cell Tracking Challenge. The $p$-values are obtained by a permutation test (10 000 repetitions) comparing each line against each column following a resampled ANOVA test (10 000 repetitions, $p_{\mathrm{Fluo-C2DL-MSC}}=1.8070 \times 10^{-1}$, $p_{\mathrm{Fluo-N2DH-GOWT1}}=5.8200 \times 10^{-2}$, $p_{\mathrm{Fluo-N2DL-HeLa}}=1.1350 \times 10^{-1}$, $p_{\mathrm{PhC-C2DH-U373}}=4.0000 \times 10^{-4}$, $p_{\mathrm{PhC-C2DL-PSC}}=6.0000 \times 10^{-4}$). Color code : increase (cyan), decrease (red), and no significant changes (black) of each line compared to each column of the metric scores.

| | MICRA-Net | Precise | BBOX |
|---|---|---|---|
| MICRA-Net | - | $7.4600 \times 10^{-1}$ | $1.1500 \times 10^{-2}$ |
| Precise | $7.4600 \times 10^{-1}$ | - | $3.4000 \times 10^{-3}$ |
| BBOX | $1.1500 \times 10^{-2}$ | $3.4000 \times 10^{-3}$ | - |

Supplementary Table 9: Statistical analysis comparing the normalized area under the curve of MICRA-Net, precise User-Study, and BBOX User-Study when *averaging all selected cell line* of the Cell Tracking Challenge. The $p$-values are obtained by a permutation test (10 000 repetitions) comparing each line against each column following a resampled ANOVA test (10 000 repetitions, $p$=<$6.4000 \times 10^{-3}$). Color code : increase (cyan), decrease (red), and no significant changes (black) of each line compared to each column of the metric scores.

| **Fluo-C2DL-MSC** | MICRA-Net | BBOX | ALT-10 | ALT-5 | ALT-2 | FS |
|---|---|---|---|---|---|---|
| MICRA-Net | - | $7.5000 \times 10^{-3}$ | $3.6400 \times 10^{-2}$ | $8.3100 \times 10^{-2}$ | $7.5770 \times 10^{-1}$ | $9.1240 \times 10^{-1}$ |
| BBOX | $7.5000 \times 10^{-3}$ | - | $8.9500 \times 10^{-2}$ | $2.3000 \times 10^{-3}$ | $6.9000 \times 10^{-3}$ | $7.4000 \times 10^{-3}$ |
| ALT-10 | $3.6400 \times 10^{-2}$ | $8.9500 \times 10^{-2}$ | - | $2.3600 \times 10^{-1}$ | $8.1000 \times 10^{-3}$ | $7.9000 \times 10^{-3}$ |
| ALT-5 | $8.3100 \times 10^{-2}$ | $2.3000 \times 10^{-3}$ | $2.3600 \times 10^{-1}$ | - | $2.2700 \times 10^{-2}$ | $8.5000 \times 10^{-3}$ |
| ALT-2 | $7.5770 \times 10^{-1}$ | $6.9000 \times 10^{-3}$ | $8.1000 \times 10^{-3}$ | $2.2700 \times 10^{-2}$ | - | $1.7700 \times 10^{-2}$ |
| FS | $9.1240 \times 10^{-1}$ | $7.4000 \times 10^{-3}$ | $7.9000 \times 10^{-3}$ | $8.5000 \times 10^{-3}$ | $1.7700 \times 10^{-2}$ | - |

| **Fluo-N2DH-GOWT1** | MICRA-Net | BBOX | ALT-10 | ALT-5 | ALT-2 | FS |
|---|---|---|---|---|---|---|
| MICRA-Net | - | $6.1000 \times 10^{-3}$ | $7.2000 \times 10^{-3}$ | $5.7000 \times 10^{-3}$ | $6.0000 \times 10^{-3}$ | $7.4000 \times 10^{-3}$ |
| BBOX | $6.1000 \times 10^{-3}$ | - | $9.2000 \times 10^{-3}$ | $2.4460 \times 10^{-1}$ | $1.3700 \times 10^{-2}$ | $6.4000 \times 10^{-3}$ |
| ALT-10 | $7.2000 \times 10^{-3}$ | $9.2000 \times 10^{-3}$ | - | $7.3000 \times 10^{-3}$ | $6.9000 \times 10^{-3}$ | $7.3000 \times 10^{-3}$ |
| ALT-5 | $5.7000 \times 10^{-3}$ | $2.4460 \times 10^{-1}$ | $7.3000 \times 10^{-3}$ | - | $8.7220 \times 10^{-1}$ | $9.7560 \times 10^{-1}$ |
| ALT-2 | $6.0000 \times 10^{-3}$ | $1.3700 \times 10^{-2}$ | $6.9000 \times 10^{-3}$ | $8.7220 \times 10^{-1}$ | - | $7.3090 \times 10^{-1}$ |
| FS | $7.4000 \times 10^{-3}$ | $6.4000 \times 10^{-3}$ | $7.3000 \times 10^{-3}$ | $9.7560 \times 10^{-1}$ | $7.3090 \times 10^{-1}$ | - |

| **Fluo-N2DL-HeLa** | MICRA-Net | BBOX | ALT-10 | ALT-5 | ALT-2 | FS |
|---|---|---|---|---|---|---|
| MICRA-Net | - | $8.4000 \times 10^{-3}$ | $7.2000 \times 10^{-3}$ | $9.7000 \times 10^{-3}$ | $6.9000 \times 10^{-3}$ | $9.5000 \times 10^{-3}$ |
| BBOX | $8.4000 \times 10^{-3}$ | - | $8.4000 \times 10^{-3}$ | $4.0000 \times 10^{-2}$ | $4.8000 \times 10^{-3}$ | $6.2000 \times 10^{-3}$ |
| ALT-10 | $7.2000 \times 10^{-3}$ | $8.4000 \times 10^{-3}$ | - | $2.5000 \times 10^{-3}$ | $7.9000 \times 10^{-3}$ | $6.9000 \times 10^{-3}$ |
| ALT-5 | $9.7000 \times 10^{-3}$ | $4.0000 \times 10^{-2}$ | $2.5000 \times 10^{-3}$ | - | $2.2700 \times 10^{-2}$ | $8.3000 \times 10^{-3}$ |
| ALT-2 | $6.9000 \times 10^{-3}$ | $4.8000 \times 10^{-3}$ | $7.9000 \times 10^{-3}$ | $2.2700 \times 10^{-2}$ | - | $9.3010 \times 10^{-1}$ |
| FS | $9.5000 \times 10^{-3}$ | $6.2000 \times 10^{-3}$ | $6.9000 \times 10^{-3}$ | $8.3000 \times 10^{-3}$ | $9.3010 \times 10^{-1}$ | - |

| **PhC-C2DH-U373** | MICRA-Net | BBOX | ALT-10 | ALT-5 | ALT-2 | FS |
|---|---|---|---|---|---|---|
| MICRA-Net | - | $6.8000 \times 10^{-3}$ | $3.3200 \times 10^{-2}$ | $5.7440 \times 10^{-1}$ | $1.1700 \times 10^{-2}$ | $5.5000 \times 10^{-3}$ |
| BBOX | $6.8000 \times 10^{-3}$ | - | $2.0270 \times 10^{-1}$ | $3.4400 \times 10^{-1}$ | $3.1000 \times 10^{-3}$ | $7.8000 \times 10^{-3}$ |
| ALT-10 | $3.3200 \times 10^{-2}$ | $2.0270 \times 10^{-1}$ | - | $1.4910 \times 10^{-1}$ | $8.0000 \times 10^{-3}$ | $1.0000 \times 10^{-4}$ |
| ALT-5 | $5.7440 \times 10^{-1}$ | $3.4400 \times 10^{-1}$ | $1.4910 \times 10^{-1}$ | - | $2.3700 \times 10^{-2}$ | $7.3000 \times 10^{-3}$ |
| ALT-2 | $1.1700 \times 10^{-2}$ | $3.1000 \times 10^{-3}$ | $8.0000 \times 10^{-3}$ | $2.3700 \times 10^{-2}$ | - | $1.5600 \times 10^{-2}$ |
| FS | $5.5000 \times 10^{-3}$ | $7.8000 \times 10^{-3}$ | $1.0000 \times 10^{-4}$ | $7.3000 \times 10^{-3}$ | $1.5600 \times 10^{-2}$ | - |

| **PhC-C2DL-PSC** | MICRA-Net | BBOX | ALT-10 | ALT-5 | ALT-2 | FS |
|---|---|---|---|---|---|---|
| MICRA-Net | - | $9.0000 \times 10^{-3}$ | $7.9370 \times 10^{-1}$ | $6.6000 \times 10^{-3}$ | $7.5000 \times 10^{-3}$ | $6.0000 \times 10^{-4}$ |
| BBOX | $9.0000 \times 10^{-3}$ | - | $6.7000 \times 10^{-3}$ | $6.8400 \times 10^{-2}$ | $7.3000 \times 10^{-3}$ | $1.0000 \times 10^{-3}$ |
| ALT-10 | $7.9370 \times 10^{-1}$ | $6.7000 \times 10^{-3}$ | - | $8.1000 \times 10^{-3}$ | $6.0000 \times 10^{-3}$ | $1.2000 \times 10^{-3}$ |
| ALT-5 | $6.6000 \times 10^{-3}$ | $6.8400 \times 10^{-2}$ | $8.1000 \times 10^{-3}$ | - | $6.2000 \times 10^{-3}$ | $7.0000 \times 10^{-4}$ |
| ALT-2 | $7.5000 \times 10^{-3}$ | $7.3000 \times 10^{-3}$ | $6.0000 \times 10^{-3}$ | $6.2000 \times 10^{-3}$ | - | $4.8900 \times 10^{-2}$ |
| FS | $6.0000 \times 10^{-4}$ | $1.0000 \times 10^{-3}$ | $1.2000 \times 10^{-3}$ | $7.0000 \times 10^{-4}$ | $4.8900 \times 10^{-2}$ | - |

Supplementary Table 10: Statistical analysis comparing the normalized area under the curve of MICRA-Net, fully- and weakly-supervised U-Net for each cell line of the Cell Tracking Challenge. The $p$-values are obtained by a permutation test (10 000 repetitions) comparing each line against each column following a resampled ANOVA test (10 000 repetitions, $p_{\text{Fluo-C2DL-MSC}}$=<$1.0000 \times 10^{-4}$, $p_{\text{Fluo-N2DH-GOWT1}}$=<$1.0000 \times 10^{-4}$, $p_{\text{Fluo-N2DL-HeLa}}$=<$1.0000 \times 10^{-4}$, $p_{\text{PhC-C2DH-U373}}$=<$1.0000 \times 10^{-4}$, $p_{\text{PhC-C2DL-PSC}}$=<$1.0000 \times 10^{-4}$). Color code : increase (cyan), decrease (red), and no significant changes (black) of each line compared to each column of the metric scores.

| | MICRA-Net | BBOX | ALT-10 | ALT-5 | ALT-2 | FS |
|---|---|---|---|---|---|---|
| MICRA-Net | - | $8.5330 \times 10^{-1}$ | $1.0000 \times 10^{-4}$ | $1.8910 \times 10^{-1}$ | $3.0000 \times 10^{-3}$ | $4.0000 \times 10^{-4}$ |
| BBOX | $8.5330 \times 10^{-1}$ | - | $3.9000 \times 10^{-3}$ | $3.6180 \times 10^{-1}$ | $2.5300 \times 10^{-2}$ | $7.3000 \times 10^{-3}$ |
| ALT-10 | $1.0000 \times 10^{-4}$ | $3.9000 \times 10^{-3}$ | - | $2.0000 \times 10^{-4}$ | $<1.0000 \times 10^{-4}$ | $<1.0000 \times 10^{-4}$ |
| ALT-5 | $1.8910 \times 10^{-1}$ | $3.6180 \times 10^{-1}$ | $2.0000 \times 10^{-4}$ | - | $1.5590 \times 10^{-1}$ | $6.4000 \times 10^{-2}$ |
| ALT-2 | $3.0000 \times 10^{-3}$ | $2.5300 \times 10^{-2}$ | $<1.0000 \times 10^{-4}$ | $1.5590 \times 10^{-1}$ | - | $6.2230 \times 10^{-1}$ |
| FS | $4.0000 \times 10^{-4}$ | $7.3000 \times 10^{-3}$ | $<1.0000 \times 10^{-4}$ | $6.4000 \times 10^{-2}$ | $6.2230 \times 10^{-1}$ | - |

Supplementary Table 11: Statistical analysis comparing the normalized area under the curve of MICRA-Net, fully- and weakly-supervised U-Net when pooling data from each cell line of the Cell Tracking Challenge. The $p$-values are obtained by a permutation test (10 000 repetitions) comparing each line against each column following a resampled ANOVA test (10 000 repetitions, $p$=$<1.0000 \times 10^{-4}$). Color code : increase (cyan), decrease (red), and no significant changes (black) of each line compared to each column of the metric scores.

| **Fluo-C2DL-MSC** | MICRA-Net | BBOX | ALT-10 | ALT-5 | ALT-2 | FS |
|---|---|---|---|---|---|---|
| MICRA-Net | - | $4.1000 \times 10^{-3}$ | $1.0200 \times 10^{-2}$ | $5.2700 \times 10^{-2}$ | $1.2900 \times 10^{-2}$ | $5.0400 \times 10^{-2}$ |
| BBOX | $4.1000 \times 10^{-3}$ | - | $4.4990 \times 10^{-1}$ | $4.7900 \times 10^{-2}$ | $8.3900 \times 10^{-2}$ | $4.5800 \times 10^{-2}$ |
| ALT-10 | $1.0200 \times 10^{-2}$ | $4.4990 \times 10^{-1}$ | - | $2.2450 \times 10^{-1}$ | $4.9140 \times 10^{-1}$ | $1.9990 \times 10^{-1}$ |
| ALT-5 | $5.2700 \times 10^{-2}$ | $4.7900 \times 10^{-2}$ | $2.2450 \times 10^{-1}$ | - | $5.3740 \times 10^{-1}$ | $9.6300 \times 10^{-1}$ |
| ALT-2 | $1.2900 \times 10^{-2}$ | $8.3900 \times 10^{-2}$ | $4.9140 \times 10^{-1}$ | $5.3740 \times 10^{-1}$ | - | $5.2580 \times 10^{-1}$ |
| FS | $5.0400 \times 10^{-2}$ | $4.5800 \times 10^{-2}$ | $1.9990 \times 10^{-1}$ | $9.6300 \times 10^{-1}$ | $5.2580 \times 10^{-1}$ | - |
| **Fluo-N2DH-GOWT1** | MICRA-Net | BBOX | ALT-10 | ALT-5 | ALT-2 | FS |
| MICRA-Net | - | $5.8890 \times 10^{-1}$ | $6.6000 \times 10^{-3}$ | $2.2200 \times 10^{-2}$ | $5.9000 \times 10^{-3}$ | $3.9100 \times 10^{-2}$ |
| BBOX | $5.8890 \times 10^{-1}$ | - | $7.6000 \times 10^{-3}$ | $2.7900 \times 10^{-2}$ | $7.5000 \times 10^{-3}$ | $2.5100 \times 10^{-2}$ |
| ALT-10 | $6.6000 \times 10^{-3}$ | $7.6000 \times 10^{-3}$ | - | $8.4000 \times 10^{-3}$ | $8.3000 \times 10^{-3}$ | $1.0100 \times 10^{-2}$ |
| ALT-5 | $2.2200 \times 10^{-2}$ | $2.7900 \times 10^{-2}$ | $8.4000 \times 10^{-3}$ | - | $5.3000 \times 10^{-3}$ | $6.6000 \times 10^{-3}$ |
| ALT-2 | $5.9000 \times 10^{-3}$ | $7.5000 \times 10^{-3}$ | $8.3000 \times 10^{-3}$ | $5.3000 \times 10^{-3}$ | - | $9.1420 \times 10^{-1}$ |
| FS | $3.9100 \times 10^{-2}$ | $2.5100 \times 10^{-2}$ | $1.0100 \times 10^{-2}$ | $6.6000 \times 10^{-3}$ | $9.1420 \times 10^{-1}$ | - |
| **Fluo-N2DL-HeLa** | MICRA-Net | BBOX | ALT-10 | ALT-5 | ALT-2 | FS |
| MICRA-Net | - | $1.0090 \times 10^{-1}$ | $7.3000 \times 10^{-3}$ | $2.7630 \times 10^{-1}$ | $5.2000 \times 10^{-3}$ | $2.0600 \times 10^{-2}$ |
| BBOX | $1.0090 \times 10^{-1}$ | - | $5.1000 \times 10^{-3}$ | $3.4000 \times 10^{-3}$ | $7.3000 \times 10^{-3}$ | $5.1800 \times 10^{-2}$ |
| ALT-10 | $7.3000 \times 10^{-3}$ | $5.1000 \times 10^{-3}$ | - | $8.7000 \times 10^{-3}$ | $6.5000 \times 10^{-3}$ | $3.7000 \times 10^{-3}$ |
| ALT-5 | $2.7630 \times 10^{-1}$ | $3.4000 \times 10^{-3}$ | $8.7000 \times 10^{-3}$ | - | $4.0000 \times 10^{-3}$ | $2.8000 \times 10^{-3}$ |
| ALT-2 | $5.2000 \times 10^{-3}$ | $7.3000 \times 10^{-3}$ | $6.5000 \times 10^{-3}$ | $4.0000 \times 10^{-3}$ | - | $5.3000 \times 10^{-2}$ |
| FS | $2.0600 \times 10^{-2}$ | $5.1800 \times 10^{-2}$ | $3.7000 \times 10^{-3}$ | $2.8000 \times 10^{-3}$ | $5.3000 \times 10^{-2}$ | - |
| **PhC-C2DH-U373** | MICRA-Net | BBOX | ALT-10 | ALT-5 | ALT-2 | FS |
| MICRA-Net | - | $2.3000 \times 10^{-3}$ | $8.5000 \times 10^{-3}$ | $8.6000 \times 10^{-3}$ | $5.1200 \times 10^{-2}$ | $2.3200 \times 10^{-2}$ |
| BBOX | $2.3000 \times 10^{-3}$ | - | $6.6020 \times 10^{-1}$ | $4.6100 \times 10^{-2}$ | $1.7490 \times 10^{-1}$ | $1.5200 \times 10^{-2}$ |
| ALT-10 | $8.5000 \times 10^{-3}$ | $6.6020 \times 10^{-1}$ | - | $1.2820 \times 10^{-1}$ | $3.5740 \times 10^{-1}$ | $8.4000 \times 10^{-2}$ |
| ALT-5 | $8.6000 \times 10^{-3}$ | $4.6100 \times 10^{-2}$ | $1.2820 \times 10^{-1}$ | - | $7.8450 \times 10^{-1}$ | $5.3630 \times 10^{-1}$ |
| ALT-2 | $5.1200 \times 10^{-2}$ | $1.7490 \times 10^{-1}$ | $3.5740 \times 10^{-1}$ | $7.8450 \times 10^{-1}$ | - | $4.5590 \times 10^{-1}$ |
| FS | $2.3200 \times 10^{-2}$ | $1.5200 \times 10^{-2}$ | $8.4000 \times 10^{-2}$ | $5.3630 \times 10^{-1}$ | $4.5590 \times 10^{-1}$ | - |
| **PhC-C2DL-PSC** | MICRA-Net | BBOX | ALT-10 | ALT-5 | ALT-2 | FS |
| MICRA-Net | - | $2.2600 \times 10^{-2}$ | $2.9200 \times 10^{-2}$ | $8.5000 \times 10^{-3}$ | $3.9000 \times 10^{-3}$ | $8.0000 \times 10^{-3}$ |
| BBOX | $2.2600 \times 10^{-2}$ | - | $8.6000 \times 10^{-3}$ | $4.7000 \times 10^{-3}$ | $2.0000 \times 10^{-3}$ | $6.6000 \times 10^{-3}$ |
| ALT-10 | $2.9200 \times 10^{-2}$ | $8.6000 \times 10^{-3}$ | - | $7.3000 \times 10^{-3}$ | $3.8000 \times 10^{-3}$ | $8.8000 \times 10^{-3}$ |
| ALT-5 | $8.5000 \times 10^{-3}$ | $4.7000 \times 10^{-3}$ | $7.3000 \times 10^{-3}$ | - | $3.9000 \times 10^{-3}$ | $7.9000 \times 10^{-3}$ |
| ALT-2 | $3.9000 \times 10^{-3}$ | $2.0000 \times 10^{-3}$ | $3.8000 \times 10^{-3}$ | $3.9000 \times 10^{-3}$ | - | $4.9270 \times 10^{-1}$ |
| FS | $8.0000 \times 10^{-3}$ | $6.6000 \times 10^{-3}$ | $8.8000 \times 10^{-3}$ | $7.9000 \times 10^{-3}$ | $4.9270 \times 10^{-1}$ | - |

Supplementary Table 12: Statistical analysis comparing the normalized area under the curve of MICRA-Net, fully- and weakly-supervised Mask R-CNN for each cell line of the Cell Tracking Challenge. The $p$-values are obtained by a permutation test (10 000 repetitions) comparing each line against each column following a resampled ANOVA test (10 000 repetitions, $p_{\text{Fluo-C2DL-MSC}}$=$3.3000 \times 10^{-3}$, $p_{\text{Fluo-N2DH-GOWT1}}$=$<1.0000 \times 10^{-4}$, $p_{\text{Fluo-N2DL-HeLa}}$=$<1.0000 \times 10^{-4}$, $p_{\text{PhC-C2DH-U373}}$=$1.1000 \times 10^{-3}$, $p_{\text{PhC-C2DL-PSC}}$=$<1.0000 \times 10^{-4}$). Color code : increase (cyan), decrease (red), and no significant changes (black) of each line compared to each column of the metric scores.

| | MICRA-Net | BBOX | ALT-10 | ALT-5 | ALT-2 | FS |
|---|---|---|---|---|---|---|
| MICRA-Net | - | $1.5820 \times 10^{-1}$ | $<1.0000 \times 10^{-4}$ | $1.8180 \times 10^{-1}$ | $2.8090 \times 10^{-1}$ | $2.1650 \times 10^{-1}$ |
| BBOX | $1.5820 \times 10^{-1}$ | - | $6.0000 \times 10^{-4}$ | $7.1580 \times 10^{-1}$ | $3.8000 \times 10^{-2}$ | $2.6900 \times 10^{-2}$ |
| ALT-10 | $<1.0000 \times 10^{-4}$ | $6.0000 \times 10^{-4}$ | - | $<1.0000 \times 10^{-4}$ | $<1.0000 \times 10^{-4}$ | $<1.0000 \times 10^{-4}$ |
| ALT-5 | $1.8180 \times 10^{-1}$ | $7.1580 \times 10^{-1}$ | $<1.0000 \times 10^{-4}$ | - | $4.1700 \times 10^{-2}$ | $2.2400 \times 10^{-2}$ |
| ALT-2 | $2.8090 \times 10^{-1}$ | $3.8000 \times 10^{-2}$ | $<1.0000 \times 10^{-4}$ | $4.1700 \times 10^{-2}$ | - | $9.7130 \times 10^{-1}$ |
| FS | $2.1650 \times 10^{-1}$ | $2.6900 \times 10^{-2}$ | $<1.0000 \times 10^{-4}$ | $2.2400 \times 10^{-2}$ | $9.7130 \times 10^{-1}$ | - |

Supplementary Table 13: Statistical analysis comparing the normalized area under the curve of MICRA-Net, fully- and weakly-supervised Mask R-CNN when pooling data from each cell line of the Cell Tracking Challenge. The $p$-values are obtained by a permutation test (10 000 repetitions) comparing each line against each column following a resampled ANOVA test (10 000 repetitions, $p$=$<1.0000 \times 10^{-4}$). Color code : increase (cyan), decrease (red), and no significant changes (black) of each line compared to each column of the metric scores.

| **Fluo-C2DL-MSC** | MICRA-Net | BBOX | ALT-10 | ALT-5 | ALT-2 | FS |
|---|---|---|---|---|---|---|
| MICRA-Net | - | $3.6000 \times 10^{-3}$ | $4.8500 \times 10^{-2}$ | $1.4910 \times 10^{-1}$ | $3.9870 \times 10^{-1}$ | $4.7010 \times 10^{-1}$ |
| BBOX | $3.6000 \times 10^{-3}$ | - | $2.8000 \times 10^{-3}$ | $8.6000 \times 10^{-3}$ | $6.9000 \times 10^{-3}$ | $7.4000 \times 10^{-3}$ |
| ALT-10 | $4.8500 \times 10^{-2}$ | $2.8000 \times 10^{-3}$ | - | $6.6000 \times 10^{-3}$ | $8.1000 \times 10^{-3}$ | $7.9000 \times 10^{-3}$ |
| ALT-5 | $1.4910 \times 10^{-1}$ | $8.6000 \times 10^{-3}$ | $6.6000 \times 10^{-3}$ | - | $1.6000 \times 10^{-2}$ | $8.5000 \times 10^{-3}$ |
| ALT-2 | $3.9870 \times 10^{-1}$ | $6.9000 \times 10^{-3}$ | $8.1000 \times 10^{-3}$ | $1.6000 \times 10^{-2}$ | - | $4.8090 \times 10^{-1}$ |
| FS | $4.7010 \times 10^{-1}$ | $7.4000 \times 10^{-3}$ | $7.9000 \times 10^{-3}$ | $8.5000 \times 10^{-3}$ | $4.8090 \times 10^{-1}$ | - |

| **Fluo-N2DH-GOWT1** | MICRA-Net | BBOX | ALT-10 | ALT-5 | ALT-2 | FS |
|---|---|---|---|---|---|---|
| MICRA-Net | - | $5.0800 \times 10^{-2}$ | $7.7000 \times 10^{-3}$ | $7.8300 \times 10^{-2}$ | $5.9000 \times 10^{-3}$ | $5.3000 \times 10^{-3}$ |
| BBOX | $5.0800 \times 10^{-2}$ | - | $3.7000 \times 10^{-3}$ | $8.3620 \times 10^{-1}$ | $8.7000 \times 10^{-3}$ | $7.9000 \times 10^{-3}$ |
| ALT-10 | $7.7000 \times 10^{-3}$ | $3.7000 \times 10^{-3}$ | - | $8.1000 \times 10^{-3}$ | $8.2000 \times 10^{-3}$ | $8.6000 \times 10^{-3}$ |
| ALT-5 | $7.8300 \times 10^{-2}$ | $8.3620 \times 10^{-1}$ | $8.1000 \times 10^{-3}$ | - | $3.1600 \times 10^{-2}$ | $1.5200 \times 10^{-2}$ |
| ALT-2 | $5.9000 \times 10^{-3}$ | $8.7000 \times 10^{-3}$ | $8.2000 \times 10^{-3}$ | $3.1600 \times 10^{-2}$ | - | $3.1500 \times 10^{-2}$ |
| FS | $5.3000 \times 10^{-3}$ | $7.9000 \times 10^{-3}$ | $8.6000 \times 10^{-3}$ | $1.5200 \times 10^{-2}$ | $3.1500 \times 10^{-2}$ | - |

| **Fluo-N2DL-HeLa** | MICRA-Net | BBOX | ALT-10 | ALT-5 | ALT-2 | FS |
|---|---|---|---|---|---|---|
| MICRA-Net | - | $5.5140 \times 10^{-1}$ | $7.1000 \times 10^{-3}$ | $6.9200 \times 10^{-2}$ | $7.4000 \times 10^{-3}$ | $9.0000 \times 10^{-3}$ |
| BBOX | $5.5140 \times 10^{-1}$ | - | $7.7000 \times 10^{-3}$ | $7.6000 \times 10^{-3}$ | $5.4000 \times 10^{-3}$ | $6.7000 \times 10^{-3}$ |
| ALT-10 | $7.1000 \times 10^{-3}$ | $7.7000 \times 10^{-3}$ | - | $8.7000 \times 10^{-3}$ | $8.1000 \times 10^{-3}$ | $6.1000 \times 10^{-3}$ |
| ALT-5 | $6.9200 \times 10^{-2}$ | $7.6000 \times 10^{-3}$ | $8.7000 \times 10^{-3}$ | - | $7.3000 \times 10^{-3}$ | $8.5000 \times 10^{-3}$ |
| ALT-2 | $7.4000 \times 10^{-3}$ | $5.4000 \times 10^{-3}$ | $8.1000 \times 10^{-3}$ | $7.3000 \times 10^{-3}$ | - | $5.6000 \times 10^{-3}$ |
| FS | $9.0000 \times 10^{-3}$ | $6.7000 \times 10^{-3}$ | $6.1000 \times 10^{-3}$ | $8.5000 \times 10^{-3}$ | $5.6000 \times 10^{-3}$ | - |

| **PhC-C2DH-U373** | MICRA-Net | BBOX | ALT-10 | ALT-5 | ALT-2 | FS |
|---|---|---|---|---|---|---|
| MICRA-Net | - | $6.7000 \times 10^{-3}$ | $7.0000 \times 10^{-3}$ | $6.8000 \times 10^{-3}$ | $7.7000 \times 10^{-3}$ | $8.7000 \times 10^{-3}$ |
| BBOX | $6.7000 \times 10^{-3}$ | - | $1.5000 \times 10^{-3}$ | $1.0100 \times 10^{-2}$ | $7.3000 \times 10^{-3}$ | $5.4000 \times 10^{-3}$ |
| ALT-10 | $7.0000 \times 10^{-3}$ | $1.5000 \times 10^{-3}$ | - | $9.4000 \times 10^{-3}$ | $6.4000 \times 10^{-3}$ | $4.4000 \times 10^{-3}$ |
| ALT-5 | $6.8000 \times 10^{-3}$ | $1.0100 \times 10^{-2}$ | $9.4000 \times 10^{-3}$ | - | $8.6000 \times 10^{-3}$ | $5.4000 \times 10^{-3}$ |
| ALT-2 | $7.7000 \times 10^{-3}$ | $7.3000 \times 10^{-3}$ | $6.4000 \times 10^{-3}$ | $8.6000 \times 10^{-3}$ | - | $5.1220 \times 10^{-1}$ |
| FS | $8.7000 \times 10^{-3}$ | $5.4000 \times 10^{-3}$ | $4.4000 \times 10^{-3}$ | $5.4000 \times 10^{-3}$ | $5.1220 \times 10^{-1}$ | - |

| **Phc-C2DL-PSC** | MICRA-Net | BBOX | ALT-10 | ALT-5 | ALT-2 | FS |
|---|---|---|---|---|---|---|
| MICRA-Net | - | $5.9000 \times 10^{-3}$ | $5.8000 \times 10^{-3}$ | $5.2550 \times 10^{-1}$ | $8.0000 \times 10^{-3}$ | $5.8000 \times 10^{-3}$ |
| BBOX | $5.9000 \times 10^{-3}$ | - | $6.0000 \times 10^{-3}$ | $6.0000 \times 10^{-3}$ | $7.0000 \times 10^{-3}$ | $4.9000 \times 10^{-3}$ |
| ALT-10 | $5.8000 \times 10^{-3}$ | $6.0000 \times 10^{-3}$ | - | $7.1000 \times 10^{-3}$ | $5.1000 \times 10^{-3}$ | $5.4000 \times 10^{-3}$ |
| ALT-5 | $5.2550 \times 10^{-1}$ | $6.0000 \times 10^{-3}$ | $7.1000 \times 10^{-3}$ | - | $3.1000 \times 10^{-3}$ | $2.3000 \times 10^{-3}$ |
| ALT-2 | $8.0000 \times 10^{-3}$ | $7.0000 \times 10^{-3}$ | $5.1000 \times 10^{-3}$ | $3.1000 \times 10^{-3}$ | - | $5.2000 \times 10^{-3}$ |
| FS | $5.8000 \times 10^{-3}$ | $4.9000 \times 10^{-3}$ | $5.4000 \times 10^{-3}$ | $2.3000 \times 10^{-3}$ | $5.2000 \times 10^{-3}$ | - |

Supplementary Table 14: Statistical analysis comparing the normalized area under the curve of MICRA-Net, fully- and weakly-supervised Ilastik for each cell line of the Cell Tracking Challenge. The $p$-values are obtained by a permutation test (10 000 repetitions) comparing each line against each column following a resampled ANOVA test (10 000 repetitions, $p_{\mathrm{Fluo-C2DL-MSC}}$=$<1.0000 \times 10^{-4}$, $p_{\mathrm{Fluo-N2DH-GOWT1}}$=$<1.0000 \times 10^{-4}$, $p_{\mathrm{Fluo-N2DL-HeLa}}$=$<1.0000 \times 10^{-4}$, $p_{\mathrm{PhC-C2DH-U373}}$=$1.0000 \times 10^{-4}$, $p_{\mathrm{PhC-C2DL-PSC}}$=$<1.0000 \times 10^{-4}$). Color code : increase (cyan), decrease (red), and no significant changes (black) of each line compared to each column of the metric scores.

| | MICRA-Net | BBOX | ALT-10 | ALT-5 | ALT-2 | FS |
|---|---|---|---|---|---|---|
| MICRA-Net | - | $2.7000 \times 10^{-3}$ | $<1.0000 \times 10^{-4}$ | $9.2000 \times 10^{-3}$ | $7.7800 \times 10^{-1}$ | $3.8540 \times 10^{-1}$ |
| BBOX | $2.7000 \times 10^{-3}$ | - | $3.8700 \times 10^{-2}$ | $2.5800 \times 10^{-1}$ | $3.9000 \times 10^{-3}$ | $1.0000 \times 10^{-3}$ |
| ALT-10 | $<1.0000 \times 10^{-4}$ | $3.8700 \times 10^{-2}$ | - | $<1.0000 \times 10^{-4}$ | $<1.0000 \times 10^{-4}$ | $<1.0000 \times 10^{-4}$ |
| ALT-5 | $9.2000 \times 10^{-3}$ | $2.5800 \times 10^{-1}$ | $<1.0000 \times 10^{-4}$ | - | $1.5200 \times 10^{-2}$ | $3.4000 \times 10^{-3}$ |
| ALT-2 | $7.7800 \times 10^{-1}$ | $3.9000 \times 10^{-3}$ | $<1.0000 \times 10^{-4}$ | $1.5200 \times 10^{-2}$ | - | $5.7350 \times 10^{-1}$ |
| FS | $3.8540 \times 10^{-1}$ | $1.0000 \times 10^{-3}$ | $<1.0000 \times 10^{-4}$ | $3.4000 \times 10^{-3}$ | $5.7350 \times 10^{-1}$ | - |

Supplementary Table 15: Statistical analysis comparing the normalized area under the curve of MICRA-Net, fully- and weakly-supervised Ilastik when pooling data from each cell line of the Cell Tracking Challenge. The $p$-values are obtained by a permutation test (10 000 repetitions) comparing each line against each column following a resampled ANOVA test (10 000 repetitions, $p$=$<1.0000 \times 10^{-4}$). Color code : increase (cyan), decrease (red), and no significant changes (black) of each line compared to each column of the metric scores.

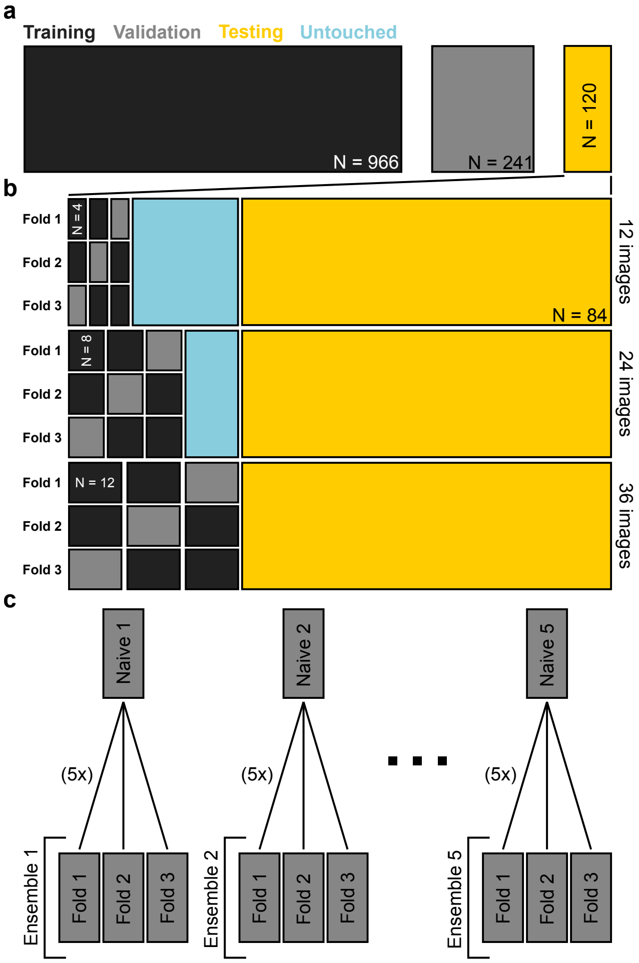


Supplementary Figure 25: Schematic of the training and fine-tuning procedure for MICRA-Net on the P. Vivax dataset. a) Data preparation: 80/20 split of the provided training set for training and validation respectively, keeping the testing set as is. b) Fine-tuning of MICRA-Net: uniform sample of $\{12, 24, 36\}$ images from the testing set. A 3-fold scheme: training on two folds and validating on a separate fold, enabling early stopping. The average prediction of the 3 fine-tuned models (termed ensemble) was used for testing. All methods were tested on the same testing set of 84 images. c) Training: 5 different models were trained on the original dataset (Naive). For fine-tuning, the 3-fold scheme was repeated 5 times, one time for each of the 5 Naive models as starting points, generating a total of 25 ensemble models.

| Condition (Number of images) | Accuracy |
|---|---|
| Naive (-) | $(0.8 \pm 0.1)\,\%$ |
| Threshold (12) | $(0.88 \pm 0.02)\,\%$ |
| Threshold (24) | $(0.87 \pm 0.02)\,\%$ |
| Threshold (36) | $(0.87 \pm 0.02)\,\%$ |
| Linear (12) | $(0.893 \pm 0.008)\,\%$ |
| Linear (24) | $(0.888 \pm 0.007)\,\%$ |
| Linear (36) | $(0.890 \pm 0.005)\,\%$ |
| Linear + 4 (12) | $(0.909 \pm 0.005)\,\%$ |
| Linear + 4 (24) | $(0.904 \pm 0.003)\,\%$ |
| Linear + 4 (36) | $(0.905 \pm 0.003)\,\%$ |
| Linear + 3, 4 (12) | $(0.911 \pm 0.003)\,\%$ |
| Linear + 3, 4 (24) | $(0.904 \pm 0.004)\,\%$ |
| Linear + 3, 4 (36) | $(0.906 \pm 0.003)\,\%$ |
| All (12) | $(0.908 \pm 0.005)\,\%$ |
| All (24) | $(0.906 \pm 0.003)\,\%$ |
| All (36) | $(0.906 \pm 0.003)\,\%$ |

Supplementary Table 16: Classification accuracy of the naive and fine-tuned models on their respective testing set on the P. Vivax dataset. The number of images used to adjust the thresholds or fine-tuning is in parentheses. The accuracy is reported as the mean ± standard deviation. For the network configuration Naive and Threshold $\{12, 24, 36\}$ (see Methods) it is calculated from 5 network instantiations. For the fine-tuned models, the scores from the ensembles that were fine-tuned from a single Naive model were averaged generating 5 classification scores that are used for calculation.

| **F1-score** | Linear (12) | Linear (24) | Linear (36) | Linear + 4 (12) | Linear + 4 (24) | Linear + 4 (36) | Linear + 3, 4 (12) | Linear + 3, 4 (24) | Linear + 3, 4 (36) | None (12) | None (24) | None (36) | Naive | Threshold (12) | Threshold (24) | Threshold (36) |
|---|---|---|---|---|---|---|---|---|---|---|---|---|---|---|---|---|
| Linear (12) | - | $8.0780 \times 10^{-1}$ | $2.3300 \times 10^{-1}$ | $3.0000 \times 10^{-3}$ | $4.9000 \times 10^{-3}$ | $5.4000 \times 10^{-3}$ | $2.7000 \times 10^{-3}$ | $9.0000 \times 10^{-4}$ | $1.7000 \times 10^{-3}$ | $2.8000 \times 10^{-3}$ | $9.0000 \times 10^{-4}$ | $4.3000 \times 10^{-3}$ | $2.4000 \times 10^{-2}$ | $1.7660 \times 10^{-1}$ | $1.5690 \times 10^{-1}$ | $2.2510 \times 10^{-1}$ |
| Linear (24) | $8.0780 \times 10^{-1}$ | - | $3.9870 \times 10^{-1}$ | $7.5000 \times 10^{-3}$ | $9.0000 \times 10^{-3}$ | $9.2000 \times 10^{-3}$ | $7.9000 \times 10^{-3}$ | $3.6000 \times 10^{-3}$ | $5.3000 \times 10^{-3}$ | $6.3000 \times 10^{-3}$ | $4.9000 \times 10^{-3}$ | $9.2000 \times 10^{-3}$ | $9.7000 \times 10^{-3}$ | $1.2000 \times 10^{-1}$ | $1.0610 \times 10^{-1}$ | $1.5400 \times 10^{-1}$ |
| Linear (36) | $2.3300 \times 10^{-1}$ | $3.9870 \times 10^{-1}$ | - | $3.9000 \times 10^{-3}$ | $5.6000 \times 10^{-3}$ | $5.4000 \times 10^{-3}$ | $3.7000 \times 10^{-3}$ | $1.7000 \times 10^{-3}$ | $2.4000 \times 10^{-3}$ | $2.1000 \times 10^{-3}$ | $1.7000 \times 10^{-3}$ | $3.8000 \times 10^{-3}$ | $7.5000 \times 10^{-3}$ | $1.6300 \times 10^{-2}$ | $2.1500 \times 10^{-2}$ | $5.3300 \times 10^{-2}$ |
| Linear + 4 (12) | $3.0000 \times 10^{-3}$ | $7.5000 \times 10^{-3}$ | $3.9000 \times 10^{-3}$ | - | $9.6000 \times 10^{-3}$ | $7.8000 \times 10^{-3}$ | $4.6000 \times 10^{-3}$ | $1.9000 \times 10^{-3}$ | $3.7000 \times 10^{-3}$ | $5.5000 \times 10^{-3}$ | $3.6000 \times 10^{-3}$ | $7.4000 \times 10^{-3}$ | $8.2000 \times 10^{-3}$ | $6.4000 \times 10^{-3}$ | $1.9000 \times 10^{-3}$ | $3.6000 \times 10^{-3}$ |
| Linear + 4 (24) | $4.9000 \times 10^{-3}$ | $9.0000 \times 10^{-3}$ | $5.6000 \times 10^{-3}$ | $9.6000 \times 10^{-3}$ | - | $2.9100 \times 10^{-2}$ | $9.0270 \times 10^{-1}$ | $7.2800 \times 10^{-2}$ | $2.0000 \times 10^{-3}$ | $7.7040 \times 10^{-1}$ | $2.6600 \times 10^{-2}$ | $4.3000 \times 10^{-3}$ | $8.7000 \times 10^{-3}$ | $7.1000 \times 10^{-3}$ | $3.9000 \times 10^{-3}$ | $6.3000 \times 10^{-3}$ |
| Linear + 4 (36) | $5.4000 \times 10^{-3}$ | $9.2000 \times 10^{-3}$ | $5.4000 \times 10^{-3}$ | $7.8000 \times 10^{-3}$ | $2.9100 \times 10^{-2}$ | - | $3.7800 \times 10^{-2}$ | $1.6260 \times 10^{-1}$ | $2.6500 \times 10^{-2}$ | $5.3400 \times 10^{-2}$ | $7.0730 \times 10^{-1}$ | $1.0300 \times 10^{-2}$ | $8.5000 \times 10^{-3}$ | $7.5000 \times 10^{-3}$ | $4.4000 \times 10^{-3}$ | $5.2000 \times 10^{-3}$ |
| Linear + 3, 4 (12) | $2.7000 \times 10^{-3}$ | $7.9000 \times 10^{-3}$ | $3.7000 \times 10^{-3}$ | $4.6000 \times 10^{-3}$ | $9.0270 \times 10^{-1}$ | $3.7800 \times 10^{-2}$ | - | $5.5900 \times 10^{-2}$ | $5.2000 \times 10^{-3}$ | $7.0480 \times 10^{-1}$ | $1.0200 \times 10^{-2}$ | $7.2000 \times 10^{-3}$ | $4.2000 \times 10^{-3}$ | $4.1000 \times 10^{-3}$ | $1.5000 \times 10^{-3}$ | $2.1000 \times 10^{-3}$ |
| Linear + 3, 4 (24) | $9.0000 \times 10^{-4}$ | $3.6000 \times 10^{-3}$ | $1.7000 \times 10^{-3}$ | $1.9000 \times 10^{-3}$ | $7.2800 \times 10^{-2}$ | $1.6260 \times 10^{-1}$ | $5.5900 \times 10^{-2}$ | - | $7.0000 \times 10^{-3}$ | $1.5170 \times 10^{-1}$ | $1.3330 \times 10^{-1}$ | $8.5000 \times 10^{-3}$ | $3.5000 \times 10^{-3}$ | $2.0000 \times 10^{-3}$ | $5.0000 \times 10^{-4}$ | $1.5000 \times 10^{-3}$ |
| Linear + 3, 4 (36) | $1.7000 \times 10^{-3}$ | $5.3000 \times 10^{-3}$ | $2.4000 \times 10^{-3}$ | $3.7000 \times 10^{-3}$ | $2.0000 \times 10^{-3}$ | $2.6500 \times 10^{-2}$ | $5.2000 \times 10^{-3}$ | $7.0000 \times 10^{-3}$ | - | $6.2000 \times 10^{-3}$ | $5.4000 \times 10^{-3}$ | $5.4090 \times 10^{-1}$ | $6.2000 \times 10^{-3}$ | $3.7000 \times 10^{-3}$ | $1.1000 \times 10^{-3}$ | $1.6000 \times 10^{-3}$ |
| None (12) | $2.8000 \times 10^{-3}$ | $6.3000 \times 10^{-3}$ | $2.1000 \times 10^{-3}$ | $5.5000 \times 10^{-3}$ | $7.7040 \times 10^{-1}$ | $5.3400 \times 10^{-2}$ | $7.0480 \times 10^{-1}$ | $1.5170 \times 10^{-1}$ | $6.2000 \times 10^{-3}$ | - | $5.0600 \times 10^{-2}$ | $7.7000 \times 10^{-3}$ | $7.1000 \times 10^{-3}$ | $5.9000 \times 10^{-3}$ | $1.0000 \times 10^{-3}$ | $2.6000 \times 10^{-3}$ |
| None (24) | $9.0000 \times 10^{-4}$ | $4.9000 \times 10^{-3}$ | $1.7000 \times 10^{-3}$ | $3.6000 \times 10^{-3}$ | $2.6600 \times 10^{-2}$ | $7.0730 \times 10^{-1}$ | $1.0200 \times 10^{-2}$ | $1.3330 \times 10^{-1}$ | $5.4000 \times 10^{-3}$ | $5.0600 \times 10^{-2}$ | - | $8.3000 \times 10^{-3}$ | $2.9000 \times 10^{-3}$ | $2.8000 \times 10^{-3}$ | $8.0000 \times 10^{-4}$ | $1.1000 \times 10^{-3}$ |
| None (36) | $4.3000 \times 10^{-3}$ | $9.2000 \times 10^{-3}$ | $3.8000 \times 10^{-3}$ | $7.4000 \times 10^{-3}$ | $4.3000 \times 10^{-3}$ | $1.0300 \times 10^{-2}$ | $7.2000 \times 10^{-3}$ | $8.5000 \times 10^{-3}$ | $5.4090 \times 10^{-1}$ | $7.7000 \times 10^{-3}$ | $8.3000 \times 10^{-3}$ | - | $6.5000 \times 10^{-3}$ | $6.6000 \times 10^{-3}$ | $4.2000 \times 10^{-3}$ | $5.5000 \times 10^{-3}$ |
| Naive | $2.4000 \times 10^{-2}$ | $9.7000 \times 10^{-3}$ | $7.5000 \times 10^{-3}$ | $8.2000 \times 10^{-3}$ | $8.7000 \times 10^{-3}$ | $8.5000 \times 10^{-3}$ | $4.2000 \times 10^{-3}$ | $3.5000 \times 10^{-3}$ | $6.2000 \times 10^{-3}$ | $7.1000 \times 10^{-3}$ | $2.9000 \times 10^{-3}$ | $6.5000 \times 10^{-3}$ | - | $9.3300 \times 10^{-2}$ | $1.0730 \times 10^{-1}$ | $8.5800 \times 10^{-2}$ |
| Threshold (12) | $1.7660 \times 10^{-1}$ | $1.2000 \times 10^{-1}$ | $1.6300 \times 10^{-2}$ | $6.4000 \times 10^{-3}$ | $7.1000 \times 10^{-3}$ | $7.5000 \times 10^{-3}$ | $4.1000 \times 10^{-3}$ | $2.0000 \times 10^{-3}$ | $3.7000 \times 10^{-3}$ | $5.9000 \times 10^{-3}$ | $2.8000 \times 10^{-3}$ | $6.6000 \times 10^{-3}$ | $9.3300 \times 10^{-2}$ | - | $9.4570 \times 10^{-1}$ | $8.5990 \times 10^{-1}$ |
| Threshold (24) | $1.5690 \times 10^{-1}$ | $1.0610 \times 10^{-1}$ | $2.1500 \times 10^{-2}$ | $1.9000 \times 10^{-3}$ | $3.9000 \times 10^{-3}$ | $4.4000 \times 10^{-3}$ | $1.5000 \times 10^{-3}$ | $5.0000 \times 10^{-4}$ | $1.1000 \times 10^{-3}$ | $1.0000 \times 10^{-3}$ | $8.0000 \times 10^{-4}$ | $4.2000 \times 10^{-3}$ | $1.0730 \times 10^{-1}$ | $9.4570 \times 10^{-1}$ | - | $9.3590 \times 10^{-1}$ |
| Threshold (36) | $2.2510 \times 10^{-1}$ | $1.5400 \times 10^{-1}$ | $5.3300 \times 10^{-2}$ | $3.6000 \times 10^{-3}$ | $6.3000 \times 10^{-3}$ | $5.2000 \times 10^{-3}$ | $2.1000 \times 10^{-3}$ | $1.5000 \times 10^{-3}$ | $1.6000 \times 10^{-3}$ | $2.6000 \times 10^{-3}$ | $1.1000 \times 10^{-3}$ | $5.5000 \times 10^{-3}$ | $8.5800 \times 10^{-2}$ | $8.5990 \times 10^{-1}$ | $9.3590 \times 10^{-1}$ | - |

Supplementary Table 17: F1-score detection metric for the fine-tuned networks on the P. Vivax dataset. The F-statistic from all groups was bootstrapped resulting in a $p$-value of $<1.0000 \times 10^{-4}$. A post-hoc resampling statistical test was performed to compare the distributions of each groups in a one-to-one manner (see Methods). Color code : increase (cyan), decrease (red), and no significant changes (black) of each line compared to each column in the F1-score.

a

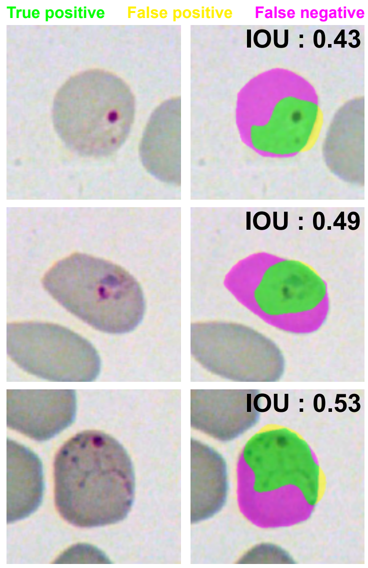


b

Segmentation (F1-score)

| | 12 images | 24 images | 36 images |
|---|---|---|---|
| Linear | 0.66 ± 0.06 | 0.67 ± 0.06 | 0.67 ± 0.06 |
| Linear + 4 | 0.61 ± 0.02 | 0.62 ± 0.02 | 0.626 ± 0.007 |
| Linear + 3, 4 | 0.59 ± 0.01 | 0.61 ± 0.02 | 0.61 ± 0.02 |
| All | 0.59 ± 0.02 | 0.60 ± 0.02 | 0.62 ± 0.02 |
| Naive | | 0.55 ± 0.07 | |

Segmentation (IOU)

| | 12 images | 24 images | 36 images |
|---|---|---|---|
| Linear | 0.52 ± 0.06 | 0.53 ± 0.06 | 0.53 ± 0.06 |
| Linear + 4 | 0.47 ± 0.02 | 0.48 ± 0.02 | 0.48 ± 0.007 |
| Linear + 3, 4 | 0.44 ± 0.02 | 0.46 ± 0.02 | 0.47 ± 0.02 |
| All | 0.44 ± 0.02 | 0.46 ± 0.02 | 0.48 ± 0.03 |
| Naive | | 0.43 ± 0.06 | |

Segmentation (SBD)

| | 12 images | 24 images | 36 images |
|---|---|---|---|
| Linear | 0.299 ± 0.006 | 0.301 ± 0.007 | 0.285 ± 0.002 |
| Linear + 4 | 0.285 ± 0.003 | 0.288 ± 0.003 | 0.287 ± 0.006 |
| Linear + 3, 4 | 0.278 ± 0.007 | 0.282 ± 0.008 | 0.283 ± 0.008 |
| All | 0.277 ± 0.009 | 0.281 ± 0.009 | 0.286 ± 0.01 |
| Naive | | 0.29 ± 0.01 | |

Supplementary Figure 26: Segmentation performance of MICRA-Net on the P. Vivax dataset. a) Example of segmentation with three different IOU. The IOU were chosen according to the average of IOU in b). Color code: True positive (green), false positive (yellow), and false negatives (magenta). b) Comparison of the 3 common performance metrics (F1-score, IOU, and SBD) between the fine-tuned and Naive models. The statistical analysis is presented in Supplementary Tab. 18. The presented scores are mean ± 95% confidence interval bootstrapped from the trained and fine-tuned models.

| **F1-score** | Naive | Linear (12) | Linear (24) | Linear (36) | Linear + 4 (12) | Linear + 4 (24) | Linear + 4 (36) | Linear + 3, 4 (12) | Linear + 3, 4 (24) | Linear + 3, 4 (36) | None (12) | None (24) | None (36) |
|---|---|---|---|---|---|---|---|---|---|---|---|---|---|
| Naive | - | $7.9000 \times 10^{-3}$ | $3.5000 \times 10^{-3}$ | $4.2000 \times 10^{-3}$ | $2.0600 \times 10^{-2}$ | $8.2000 \times 10^{-3}$ | $7.1000 \times 10^{-3}$ | $1.6810 \times 10^{-1}$ | $2.7400 \times 10^{-2}$ | $3.7400 \times 10^{-2}$ | $1.8570 \times 10^{-1}$ | $6.9700 \times 10^{-2}$ | $2.2000 \times 10^{-2}$ |
| Linear (12) | $7.9000 \times 10^{-3}$ | - | $7.3650 \times 10^{-1}$ | $7.3230 \times 10^{-1}$ | $8.9700 \times 10^{-2}$ | $1.6070 \times 10^{-1}$ | $1.4110 \times 10^{-1}$ | $9.0000 \times 10^{-3}$ | $5.7300 \times 10^{-2}$ | $5.2600 \times 10^{-2}$ | $1.6800 \times 10^{-2}$ | $3.0900 \times 10^{-2}$ | $1.1030 \times 10^{-1}$ |
| Linear (24) | $3.5000 \times 10^{-3}$ | $7.3650 \times 10^{-1}$ | - | $9.3230 \times 10^{-1}$ | $4.1200 \times 10^{-2}$ | $7.6800 \times 10^{-2}$ | $8.3200 \times 10^{-2}$ | $5.1000 \times 10^{-3}$ | $1.5200 \times 10^{-2}$ | $2.7900 \times 10^{-2}$ | $4.0000 \times 10^{-3}$ | $1.2900 \times 10^{-2}$ | $5.5000 \times 10^{-2}$ |
| Linear (36) | $4.2000 \times 10^{-3}$ | $7.3230 \times 10^{-1}$ | $9.3230 \times 10^{-1}$ | - | $4.8000 \times 10^{-2}$ | $9.7700 \times 10^{-2}$ | $9.7900 \times 10^{-2}$ | $5.9000 \times 10^{-3}$ | $3.3000 \times 10^{-2}$ | $3.7600 \times 10^{-2}$ | $1.2900 \times 10^{-2}$ | $2.1200 \times 10^{-2}$ | $7.0800 \times 10^{-2}$ |
| Linear + 4 (12) | $2.0600 \times 10^{-2}$ | $8.9700 \times 10^{-2}$ | $4.1200 \times 10^{-2}$ | $4.8000 \times 10^{-2}$ | - | $4.2040 \times 10^{-1}$ | $2.6080 \times 10^{-1}$ | $3.3500 \times 10^{-2}$ | $6.0020 \times 10^{-1}$ | $6.8650 \times 10^{-1}$ | $4.4400 \times 10^{-2}$ | $2.5840 \times 10^{-1}$ | $8.4530 \times 10^{-1}$ |
| Linear + 4 (24) | $8.2000 \times 10^{-3}$ | $1.6070 \times 10^{-1}$ | $7.6800 \times 10^{-2}$ | $9.7700 \times 10^{-2}$ | $4.2040 \times 10^{-1}$ | - | $8.9200 \times 10^{-1}$ | $6.9000 \times 10^{-3}$ | $1.7440 \times 10^{-1}$ | $2.4640 \times 10^{-1}$ | $2.2300 \times 10^{-2}$ | $8.6600 \times 10^{-2}$ | $5.5920 \times 10^{-1}$ |
| Linear + 4 (36) | $7.1000 \times 10^{-3}$ | $1.4110 \times 10^{-1}$ | $8.3200 \times 10^{-2}$ | $9.7900 \times 10^{-2}$ | $2.6080 \times 10^{-1}$ | $8.9200 \times 10^{-1}$ | - | $8.9000 \times 10^{-3}$ | $3.8000 \times 10^{-2}$ | $1.2990 \times 10^{-1}$ | $1.0100 \times 10^{-2}$ | $3.2500 \times 10^{-2}$ | $3.6820 \times 10^{-1}$ |
| Linear + 3, 4 (12) | $1.6810 \times 10^{-1}$ | $9.0000 \times 10^{-3}$ | $5.1000 \times 10^{-3}$ | $5.9000 \times 10^{-3}$ | $3.3500 \times 10^{-2}$ | $6.9000 \times 10^{-3}$ | $8.9000 \times 10^{-3}$ | - | $3.2400 \times 10^{-2}$ | $7.3000 \times 10^{-2}$ | $9.8500 \times 10^{-1}$ | $1.3920 \times 10^{-1}$ | $2.7700 \times 10^{-2}$ |
| Linear + 3, 4 (24) | $2.7400 \times 10^{-2}$ | $5.7300 \times 10^{-2}$ | $1.5200 \times 10^{-2}$ | $3.3000 \times 10^{-2}$ | $6.0020 \times 10^{-1}$ | $1.7440 \times 10^{-1}$ | $3.8000 \times 10^{-2}$ | $3.2400 \times 10^{-2}$ | - | $9.6690 \times 10^{-1}$ | $6.9200 \times 10^{-2}$ | $4.0800 \times 10^{-1}$ | $5.4980 \times 10^{-1}$ |
| Linear + 3, 4 (36) | $3.7400 \times 10^{-2}$ | $5.2600 \times 10^{-2}$ | $2.7900 \times 10^{-2}$ | $3.7600 \times 10^{-2}$ | $6.8650 \times 10^{-1}$ | $2.4640 \times 10^{-1}$ | $1.2990 \times 10^{-1}$ | $7.3000 \times 10^{-2}$ | $9.6690 \times 10^{-1}$ | - | $9.8400 \times 10^{-2}$ | $4.9120 \times 10^{-1}$ | $5.8520 \times 10^{-1}$ |
| None (12) | $1.8570 \times 10^{-1}$ | $1.6800 \times 10^{-2}$ | $4.0000 \times 10^{-3}$ | $1.2900 \times 10^{-2}$ | $4.4400 \times 10^{-2}$ | $2.2300 \times 10^{-2}$ | $1.0100 \times 10^{-2}$ | $9.8500 \times 10^{-1}$ | $6.9200 \times 10^{-2}$ | $9.8400 \times 10^{-2}$ | - | $2.0500 \times 10^{-1}$ | $5.1200 \times 10^{-2}$ |
| None (24) | $6.9700 \times 10^{-2}$ | $3.0900 \times 10^{-2}$ | $1.2900 \times 10^{-2}$ | $2.1200 \times 10^{-2}$ | $2.5840 \times 10^{-1}$ | $8.6600 \times 10^{-2}$ | $3.2500 \times 10^{-2}$ | $1.3920 \times 10^{-1}$ | $4.0800 \times 10^{-1}$ | $4.9120 \times 10^{-1}$ | $2.0500 \times 10^{-1}$ | - | $2.3590 \times 10^{-1}$ |
| None (36) | $2.2000 \times 10^{-2}$ | $1.1030 \times 10^{-1}$ | $5.5000 \times 10^{-2}$ | $7.0800 \times 10^{-2}$ | $8.4530 \times 10^{-1}$ | $5.5920 \times 10^{-1}$ | $3.6820 \times 10^{-1}$ | $2.7700 \times 10^{-2}$ | $5.4980 \times 10^{-1}$ | $5.8520 \times 10^{-1}$ | $5.1200 \times 10^{-2}$ | $2.3590 \times 10^{-1}$ | - |

| **IOU** | Naive | Linear (12) | Linear (24) | Linear (36) | Linear + 4 (12) | Linear + 4 (24) | Linear + 4 (36) | Linear + 3, 4 (12) | Linear + 3, 4 (24) | Linear + 3, 4 (36) | None (12) | None (24) | None (36) |
|---|---|---|---|---|---|---|---|---|---|---|---|---|---|
| Naive | - | $1.2800 \times 10^{-2}$ | $1.6500 \times 10^{-2}$ | $1.4500 \times 10^{-2}$ | $9.2400 \times 10^{-2}$ | $5.3700 \times 10^{-2}$ | $1.6200 \times 10^{-2}$ | $6.1920 \times 10^{-1}$ | $1.3960 \times 10^{-1}$ | $1.0960 \times 10^{-1}$ | $5.7050 \times 10^{-1}$ | $1.8490 \times 10^{-1}$ | $6.3900 \times 10^{-2}$ |
| Linear (12) | $1.2800 \times 10^{-2}$ | - | $7.1100 \times 10^{-1}$ | $7.2740 \times 10^{-1}$ | $6.2800 \times 10^{-2}$ | $1.1180 \times 10^{-1}$ | $1.2720 \times 10^{-1}$ | $1.2000 \times 10^{-3}$ | $3.4000 \times 10^{-2}$ | $4.3500 \times 10^{-2}$ | $9.4000 \times 10^{-3}$ | $3.1900 \times 10^{-2}$ | $1.2870 \times 10^{-1}$ |
| Linear (24) | $1.6500 \times 10^{-2}$ | $7.1100 \times 10^{-1}$ | - | $9.1980 \times 10^{-1}$ | $3.5300 \times 10^{-2}$ | $6.2800 \times 10^{-2}$ | $6.6500 \times 10^{-2}$ | $2.4000 \times 10^{-3}$ | $1.1800 \times 10^{-2}$ | $3.2400 \times 10^{-2}$ | $5.9000 \times 10^{-3}$ | $7.7000 \times 10^{-3}$ | $5.6400 \times 10^{-2}$ |
| Linear (36) | $1.4500 \times 10^{-2}$ | $7.2740 \times 10^{-1}$ | $9.1980 \times 10^{-1}$ | - | $4.0100 \times 10^{-2}$ | $5.5300 \times 10^{-2}$ | $6.3800 \times 10^{-2}$ | $1.9000 \times 10^{-3}$ | $2.0400 \times 10^{-2}$ | $3.2000 \times 10^{-2}$ | $1.0100 \times 10^{-2}$ | $1.6000 \times 10^{-2}$ | $6.1100 \times 10^{-2}$ |
| Linear + 4 (12) | $9.2400 \times 10^{-2}$ | $6.2800 \times 10^{-2}$ | $3.5300 \times 10^{-2}$ | $4.0100 \times 10^{-2}$ | - | $4.4250 \times 10^{-1}$ | $2.2220 \times 10^{-1}$ | $4.3400 \times 10^{-2}$ | $6.8790 \times 10^{-1}$ | $9.1590 \times 10^{-1}$ | $6.2900 \times 10^{-2}$ | $4.1720 \times 10^{-1}$ | $5.5000 \times 10^{-1}$ |
| Linear + 4 (24) | $5.3700 \times 10^{-2}$ | $1.1180 \times 10^{-1}$ | $6.2800 \times 10^{-2}$ | $5.5300 \times 10^{-2}$ | $4.4250 \times 10^{-1}$ | - | $7.1200 \times 10^{-1}$ | $1.1200 \times 10^{-2}$ | $2.1990 \times 10^{-1}$ | $3.6340 \times 10^{-1}$ | $3.4300 \times 10^{-2}$ | $1.4010 \times 10^{-1}$ | $9.2850 \times 10^{-1}$ |
| Linear + 4 (36) | $1.6200 \times 10^{-2}$ | $1.2720 \times 10^{-1}$ | $6.6500 \times 10^{-2}$ | $6.3800 \times 10^{-2}$ | $2.2220 \times 10^{-1}$ | $7.1200 \times 10^{-1}$ | - | $5.6000 \times 10^{-3}$ | $6.0000 \times 10^{-2}$ | $1.6150 \times 10^{-1}$ | $6.7000 \times 10^{-3}$ | $3.5200 \times 10^{-2}$ | $7.0420 \times 10^{-1}$ |
| Linear + 3, 4 (12) | $6.1920 \times 10^{-1}$ | $1.2000 \times 10^{-3}$ | $2.4000 \times 10^{-3}$ | $1.9000 \times 10^{-3}$ | $4.3400 \times 10^{-2}$ | $1.1200 \times 10^{-2}$ | $5.6000 \times 10^{-3}$ | - | $3.8500 \times 10^{-2}$ | $4.4600 \times 10^{-2}$ | $9.6250 \times 10^{-1}$ | $1.1480 \times 10^{-1}$ | $9.4000 \times 10^{-3}$ |
| Linear + 3, 4 (24) | $1.3960 \times 10^{-1}$ | $3.4000 \times 10^{-2}$ | $1.1800 \times 10^{-2}$ | $2.0400 \times 10^{-2}$ | $6.8790 \times 10^{-1}$ | $2.1990 \times 10^{-1}$ | $6.0000 \times 10^{-2}$ | $3.8500 \times 10^{-2}$ | - | $8.1610 \times 10^{-1}$ | $9.2800 \times 10^{-2}$ | $6.3950 \times 10^{-1}$ | $2.7390 \times 10^{-1}$ |
| Linear + 3, 4 (36) | $1.0960 \times 10^{-1}$ | $4.3500 \times 10^{-2}$ | $3.2400 \times 10^{-2}$ | $3.2000 \times 10^{-2}$ | $9.1590 \times 10^{-1}$ | $3.6340 \times 10^{-1}$ | $1.6150 \times 10^{-1}$ | $4.4600 \times 10^{-2}$ | $8.1610 \times 10^{-1}$ | - | $8.6900 \times 10^{-2}$ | $5.2110 \times 10^{-1}$ | $4.6470 \times 10^{-1}$ |
| None (12) | $5.7050 \times 10^{-1}$ | $9.4000 \times 10^{-3}$ | $5.9000 \times 10^{-3}$ | $1.0100 \times 10^{-2}$ | $6.2900 \times 10^{-2}$ | $3.4300 \times 10^{-2}$ | $6.7000 \times 10^{-3}$ | $9.6250 \times 10^{-1}$ | $9.2800 \times 10^{-2}$ | $8.6900 \times 10^{-2}$ | - | $2.0240 \times 10^{-1}$ | $5.1100 \times 10^{-2}$ |
| None (24) | $1.8490 \times 10^{-1}$ | $3.1900 \times 10^{-2}$ | $7.7000 \times 10^{-3}$ | $1.6000 \times 10^{-2}$ | $4.1720 \times 10^{-1}$ | $1.4010 \times 10^{-1}$ | $3.5200 \times 10^{-2}$ | $1.1480 \times 10^{-1}$ | $6.3950 \times 10^{-1}$ | $5.2110 \times 10^{-1}$ | $2.0240 \times 10^{-1}$ | - | $1.8450 \times 10^{-1}$ |
| None (36) | $6.3900 \times 10^{-2}$ | $1.2870 \times 10^{-1}$ | $5.6400 \times 10^{-2}$ | $6.1100 \times 10^{-2}$ | $5.5000 \times 10^{-1}$ | $9.2850 \times 10^{-1}$ | $7.0420 \times 10^{-1}$ | $9.4000 \times 10^{-3}$ | $2.7390 \times 10^{-1}$ | $4.6470 \times 10^{-1}$ | $5.1100 \times 10^{-2}$ | $1.8450 \times 10^{-1}$ | - |

| **SBD** | Naive | Linear (12) | Linear (24) | Linear (36) | Linear + 4 (12) | Linear + 4 (24) | Linear + 4 (36) | Linear + 3, 4 (12) | Linear + 3, 4 (24) | Linear + 3, 4 (36) | None (12) | None (24) | None (36) |
|---|---|---|---|---|---|---|---|---|---|---|---|---|---|
| Naive | - | $6.8600 \times 10^{-2}$ | $4.5000 \times 10^{-2}$ | $4.6200 \times 10^{-2}$ | $3.8720 \times 10^{-1}$ | $7.6360 \times 10^{-1}$ | $6.8260 \times 10^{-1}$ | $4.9900 \times 10^{-2}$ | $2.2260 \times 10^{-1}$ | $3.4230 \times 10^{-1}$ | $6.7600 \times 10^{-2}$ | $1.5540 \times 10^{-1}$ | $6.4290 \times 10^{-1}$ |
| Linear (12) | $6.8600 \times 10^{-2}$ | - | $5.3590 \times 10^{-1}$ | $5.6170 \times 10^{-1}$ | $7.5000 \times 10^{-3}$ | $8.2000 \times 10^{-3}$ | $1.6100 \times 10^{-2}$ | $9.3000 \times 10^{-3}$ | $8.8000 \times 10^{-3}$ | $1.4400 \times 10^{-2}$ | $8.2000 \times 10^{-3}$ | $7.7000 \times 10^{-3}$ | $4.2700 \times 10^{-2}$ |
| Linear (24) | $4.5000 \times 10^{-2}$ | $5.3590 \times 10^{-1}$ | - | $8.8700 \times 10^{-1}$ | $9.2000 \times 10^{-3}$ | $8.4000 \times 10^{-3}$ | $8.2000 \times 10^{-3}$ | $8.2000 \times 10^{-3}$ | $9.0000 \times 10^{-3}$ | $7.9000 \times 10^{-3}$ | $8.4000 \times 10^{-3}$ | $6.9000 \times 10^{-3}$ | $3.0100 \times 10^{-2}$ |
| Linear (36) | $4.6200 \times 10^{-2}$ | $5.6170 \times 10^{-1}$ | $8.8700 \times 10^{-1}$ | - | $7.4000 \times 10^{-3}$ | $7.8000 \times 10^{-3}$ | $8.5000 \times 10^{-3}$ | $7.2000 \times 10^{-3}$ | $6.5000 \times 10^{-3}$ | $9.4000 \times 10^{-3}$ | $7.0000 \times 10^{-3}$ | $7.4000 \times 10^{-3}$ | $3.0200 \times 10^{-2}$ |
| Linear + 4 (12) | $3.8720 \times 10^{-1}$ | $7.5000 \times 10^{-3}$ | $9.2000 \times 10^{-3}$ | $7.4000 \times 10^{-3}$ | - | $1.0200 \times 10^{-1}$ | $4.8380 \times 10^{-1}$ | $1.6400 \times 10^{-2}$ | $4.2740 \times 10^{-1}$ | $7.0730 \times 10^{-1}$ | $6.6900 \times 10^{-2}$ | $2.4470 \times 10^{-1}$ | $8.4030 \times 10^{-1}$ |
| Linear + 4 (24) | $7.6360 \times 10^{-1}$ | $8.2000 \times 10^{-3}$ | $8.4000 \times 10^{-3}$ | $7.8000 \times 10^{-3}$ | $1.0200 \times 10^{-1}$ | - | $7.5170 \times 10^{-1}$ | $6.5000 \times 10^{-3}$ | $4.5500 \times 10^{-2}$ | $3.0560 \times 10^{-1}$ | $2.1200 \times 10^{-2}$ | $1.1850 \times 10^{-1}$ | $7.5620 \times 10^{-1}$ |
| Linear + 4 (36) | $6.8260 \times 10^{-1}$ | $1.6100 \times 10^{-2}$ | $8.2000 \times 10^{-3}$ | $8.5000 \times 10^{-3}$ | $4.8380 \times 10^{-1}$ | $7.5170 \times 10^{-1}$ | - | $3.0300 \times 10^{-2}$ | $1.9390 \times 10^{-1}$ | $4.0800 \times 10^{-1}$ | $5.6700 \times 10^{-2}$ | $1.7200 \times 10^{-1}$ | $8.9550 \times 10^{-1}$ |
| Linear + 3, 4 (12) | $4.9900 \times 10^{-2}$ | $9.3000 \times 10^{-3}$ | $8.2000 \times 10^{-3}$ | $7.2000 \times 10^{-3}$ | $1.6400 \times 10^{-2}$ | $6.5000 \times 10^{-3}$ | $3.0300 \times 10^{-2}$ | - | $3.2900 \times 10^{-1}$ | $2.0230 \times 10^{-1}$ | $7.9220 \times 10^{-1}$ | $5.1770 \times 10^{-1}$ | $1.6090 \times 10^{-1}$ |
| Linear + 3, 4 (24) | $2.2260 \times 10^{-1}$ | $8.8000 \times 10^{-3}$ | $9.0000 \times 10^{-3}$ | $6.5000 \times 10^{-3}$ | $4.2740 \times 10^{-1}$ | $4.5500 \times 10^{-2}$ | $1.9390 \times 10^{-1}$ | $3.2900 \times 10^{-1}$ | - | $7.3130 \times 10^{-1}$ | $2.7170 \times 10^{-1}$ | $7.7540 \times 10^{-1}$ | $4.4860 \times 10^{-1}$ |
| Linear + 3, 4 (36) | $3.4230 \times 10^{-1}$ | $1.4400 \times 10^{-2}$ | $7.9000 \times 10^{-3}$ | $9.4000 \times 10^{-3}$ | $7.0730 \times 10^{-1}$ | $3.0560 \times 10^{-1}$ | $4.0800 \times 10^{-1}$ | $2.0230 \times 10^{-1}$ | $7.3130 \times 10^{-1}$ | - | $1.6080 \times 10^{-1}$ | $5.3890 \times 10^{-1}$ | $6.6080 \times 10^{-1}$ |
| None (12) | $6.7600 \times 10^{-2}$ | $8.2000 \times 10^{-3}$ | $8.4000 \times 10^{-3}$ | $7.0000 \times 10^{-3}$ | $6.6900 \times 10^{-2}$ | $2.1200 \times 10^{-2}$ | $5.6700 \times 10^{-2}$ | $7.9220 \times 10^{-1}$ | $2.7170 \times 10^{-1}$ | $1.6080 \times 10^{-1}$ | - | $4.3890 \times 10^{-1}$ | $1.4020 \times 10^{-1}$ |
| None (24) | $1.5540 \times 10^{-1}$ | $7.7000 \times 10^{-3}$ | $6.9000 \times 10^{-3}$ | $7.4000 \times 10^{-3}$ | $2.4470 \times 10^{-1}$ | $1.1850 \times 10^{-1}$ | $1.7200 \times 10^{-1}$ | $5.1770 \times 10^{-1}$ | $7.7540 \times 10^{-1}$ | $5.3890 \times 10^{-1}$ | $4.3890 \times 10^{-1}$ | - | $3.6500 \times 10^{-1}$ |
| None (36) | $6.4290 \times 10^{-1}$ | $4.2700 \times 10^{-2}$ | $3.0100 \times 10^{-2}$ | $3.0200 \times 10^{-2}$ | $8.4030 \times 10^{-1}$ | $7.5620 \times 10^{-1}$ | $8.9550 \times 10^{-1}$ | $1.6090 \times 10^{-1}$ | $4.4860 \times 10^{-1}$ | $6.6080 \times 10^{-1}$ | $1.4020 \times 10^{-1}$ | $3.6500 \times 10^{-1}$ | - |

Supplementary Table 18: Statistical analysis on the comparison of the F1-score, IOU, and SBD segmentation metric for the fine-tuned networks for the P. Vivax dataset. The main results are reported in Figure 26. The F-statistic from all groups was bootstrapped resulting in a $p$-value of $<1.0000 \times 10^{-4}$. A post-hoc resampling statistical test was performed to compare the distributions of each groups in a one-to-one manner (see Methods). Color code : increase (cyan), decrease (red), and no significant changes (black) of each line compared to each column of the metric scores.

| Models | Accuracy |
|---|---|
| 2 : 1 | $(84 \pm 1)$ % |
| 1 : 1 | $(86.7 \pm 0.8)$ % |
| 1 : 2 | $(88 \pm 1)$ % |
| 1 : 5 | $(88.4 \pm 0.9)$ % |
| 1 : 8 | $(89 \pm 1)$ % |
| 1 : 16 | $(89.3 \pm 0.6)$ % |

Supplementary Table 19: Classification accuracy of the trained MICRA-Net models using different positive-unlabeled ratio on the scanning electron microscopy dataset. The accuracy is reported as the mean ± standard deviation calculated from 5 network models.

| **F1-score** | 2 : 1 | 1 : 1 | 1 : 2 | 1 : 5 | 1 : 8 | 1 : 16 |
|---|---|---|---|---|---|---|
| 2 : 1 | - | $2.8900 \times 10^{-2}$ | $5.0000 \times 10^{-3}$ | $4.3000 \times 10^{-3}$ | $5.5000 \times 10^{-3}$ | $7.4000 \times 10^{-3}$ |
| 1 : 1 | $2.8900 \times 10^{-2}$ | - | $1.3520 \times 10^{-1}$ | $2.3100 \times 10^{-2}$ | $1.3140 \times 10^{-1}$ | $4.7400 \times 10^{-2}$ |
| 1 : 2 | $5.0000 \times 10^{-3}$ | $1.3520 \times 10^{-1}$ | - | $2.7600 \times 10^{-1}$ | $6.8550 \times 10^{-1}$ | $4.5460 \times 10^{-1}$ |
| 1 : 5 | $4.3000 \times 10^{-3}$ | $2.3100 \times 10^{-2}$ | $2.7600 \times 10^{-1}$ | - | $3.9850 \times 10^{-1}$ | $7.1410 \times 10^{-1}$ |
| 1 : 8 | $5.5000 \times 10^{-3}$ | $1.3140 \times 10^{-1}$ | $6.8550 \times 10^{-1}$ | $3.9850 \times 10^{-1}$ | - | $6.6630 \times 10^{-1}$ |
| 1 : 16 | $7.4000 \times 10^{-3}$ | $4.7400 \times 10^{-2}$ | $4.5460 \times 10^{-1}$ | $7.1410 \times 10^{-1}$ | $6.6630 \times 10^{-1}$ | - |

Supplementary Table 20: Statistical analysis on the comparison of the F1-score detection metric of MICRA-Net for different positive-unlabeled ratios on the scanning electron microscopy dataset (Figure 6). The F-statistic from all groups was bootstrapped resulting in a $p$-value of 0. A post-hoc resampling statistical test was performed to compare the distributions of each groups in a one-to-one manner (see Methods). Color code : increase (cyan), decrease (red), and no significant changes (black) of each line compared to each column in the F1-score.

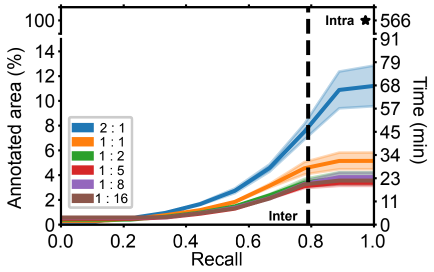


Supplementary Figure 27: Validation of MICRA-Net suggestions accuracy. Two Experts with variable levels of experience (high: [A], and intermediate: [B]) annotated all positive detections from MICRA-Net with a detection threshold set to have a recall of 1, *i.e.* all the original identifications (from Expert [A], generated *without* MICRA-Net assistance) are detected by MICRA-Net. An intra-expert (Expert [A]) recall of 0.973 was calculated for the visualisation of 100% of the field of view (without MICRA-Net assistance, star marker). Comparison of the decisions of [B] with [A] resulted in a inter-expert recall of 0.791 (dashed line). Using MICRA-Net as expert annotation assistance reduces the detection time by 25 folds (right vertical axis) while maintaining intra- and inter-experts recall levels. For constant recall, positive-unlabeled ratios of 1:2 and above allow a reduction of the annotation time compared to the 2:1 and 1:1 ratios, showing the importance of negative instances in the training dataset. The solid lines and associated pale regions are the bootstrapped mean and 95% confidence interval of 5 networks instantiated randomly for each condition.

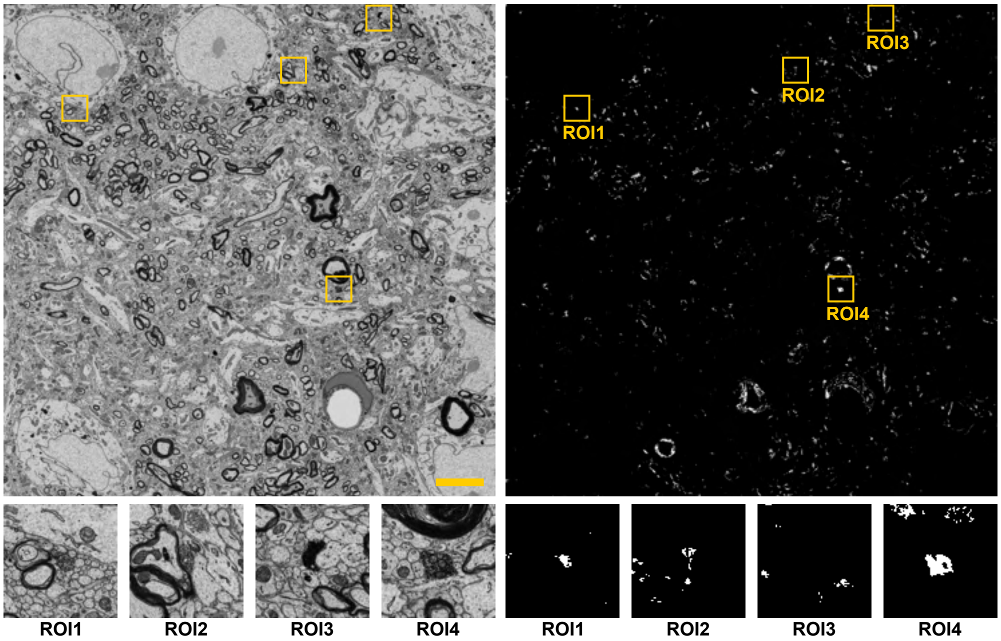


Supplementary Figure 28: Representative example of Ilastik trained on the scanning electron microscopy dataset in a weakly-supervised fashion. Point annotations with a constant radius of 10 pixels were used to detect Axon-DAB markers. This ensured that no positive pixels were outside of the Axon-DAB markers. Following training, Ilastik infers the image (*left)*) to output a binary map on the presence of an Axon-DAB marker (*right*, white). The positive regions (labeled as positive by an Expert) are extracted and shown in the bottom row. Scale bar: 5 µm. Crops: 2.56 µm × 2.56 µm

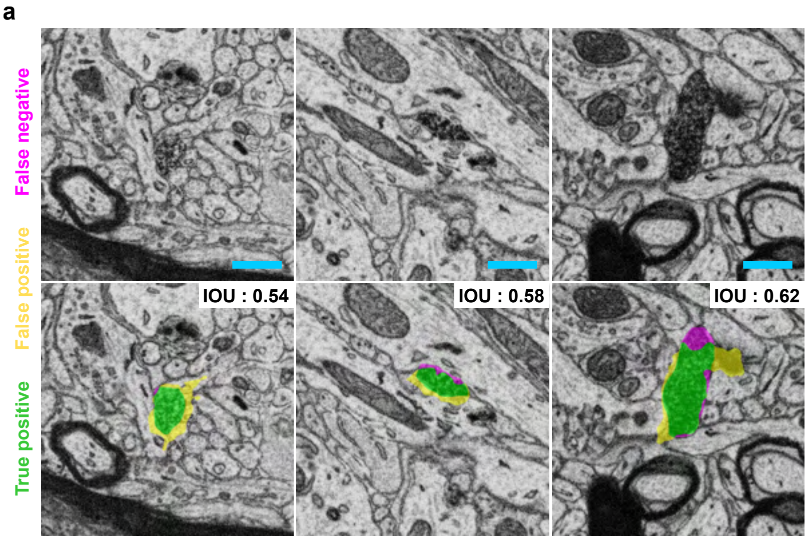


**b**

| | PU ratio | Score |
|---|---|---|
| F1-score | 2 : 1 | 0.71 ± 0.04 |
| | 1 : 1 | 0.76 ± 0.03 |
| | 1 : 2 | 0.75 ± 0.06 |
| | 1 : 5 | 0.76 ± 0.03 |
| | 1 : 8 | 0.73 ± 0.02 |
| | 1 : 16 | 0.72 ± 0.07 |
| IOU | 2 : 1 | 0.55 ± 0.04 |
| | 1 : 1 | 0.61 ± 0.03 |
| | 1 : 2 | 0.59 ± 0.07 |
| | 1 : 5 | 0.62 ± 0.03 |
| | 1 : 8 | 0.58 ± 0.02 |
| | 1 : 16 | 0.56 ± 0.08 |
| SBD | 2 : 1 | 0.298 ± 0.009 |
| | 1 : 1 | 0.31 ± 0.01 |
| | 1 : 2 | 0.32 ± 0.01 |
| | 1 : 5 | 0.318 ± 0.005 |
| | 1 : 8 | 0.31 ± 0.01 |
| | 1 : 16 | 0.30 ± 0.01 |

Supplementary Figure 29: Segmentation performance of MICRA-Net on the scanning electron microscopy dataset. a) Original images (top) with their corresponding segmentation (bottom). Scale bars are 1 µm. Color code : true positives (green), false positives (yellow), and false negatives (magenta). b) F1-score, IOU, and SBD evaluated on the precisely testing set. Reported is the mean ± 95% confidence interval from 5 network instantiations. A statistical difference was measured with resampling for SBD only ($p_{\text{SBD}}$ = 0.0059, $p_{\text{F1-score}=0.1831}$, and $p_{\text{IOU}=0.1845}$). The post-hoc resampling statistical test for SBD is reported in Supplementary Tab. 21.

| SBD | 2 : 1 | 1 : 1 | 1 : 2 | 1 : 5 | 1 : 8 | 1 : 16 |
|---|---|---|---|---|---|---|
| 2 : 1 | - | $4.7000 \times 10^{-2}$ | $2.1000 \times 10^{-2}$ | $4.5000 \times 10^{-3}$ | $3.5800 \times 10^{-2}$ | $6.6000 \times 10^{-1}$ |
| 1 : 1 | $4.7000 \times 10^{-2}$ | - | $2.0040 \times 10^{-1}$ | $5.4100 \times 10^{-2}$ | $7.4360 \times 10^{-1}$ | $2.5440 \times 10^{-1}$ |
| 1 : 2 | $2.1000 \times 10^{-2}$ | $2.0040 \times 10^{-1}$ | - | $7.2870 \times 10^{-1}$ | $5.3210 \times 10^{-1}$ | $5.6200 \times 10^{-2}$ |
| 1 : 5 | $4.5000 \times 10^{-3}$ | $5.4100 \times 10^{-2}$ | $7.2870 \times 10^{-1}$ | - | $3.3020 \times 10^{-1}$ | $2.0300 \times 10^{-2}$ |
| 1 : 8 | $3.5800 \times 10^{-2}$ | $7.4360 \times 10^{-1}$ | $5.3210 \times 10^{-1}$ | $3.3020 \times 10^{-1}$ | - | $1.7660 \times 10^{-1}$ |
| 1 : 16 | $6.6000 \times 10^{-1}$ | $2.5440 \times 10^{-1}$ | $5.6200 \times 10^{-2}$ | $2.0300 \times 10^{-2}$ | $1.7660 \times 10^{-1}$ | - |

Supplementary Table 21: Statistical analysis on the comparison of the SBD segmentation metric of MICRA-Net for different positive-unlabeled ratio on the scanning electron microscopy dataset. The main results are reported in Figure 29. The F-statistic from all groups was bootstrapped (10 000 repetitions) resulting in a $p$-value of 0.0059. A post-hoc resampling statistical test (10 000 repetitions) was performed to compare the distributions of each groups in a one-to-one manner (see Methods). Color code: increase (cyan), decrease (red), and no significant changes (black) in the F1-score.

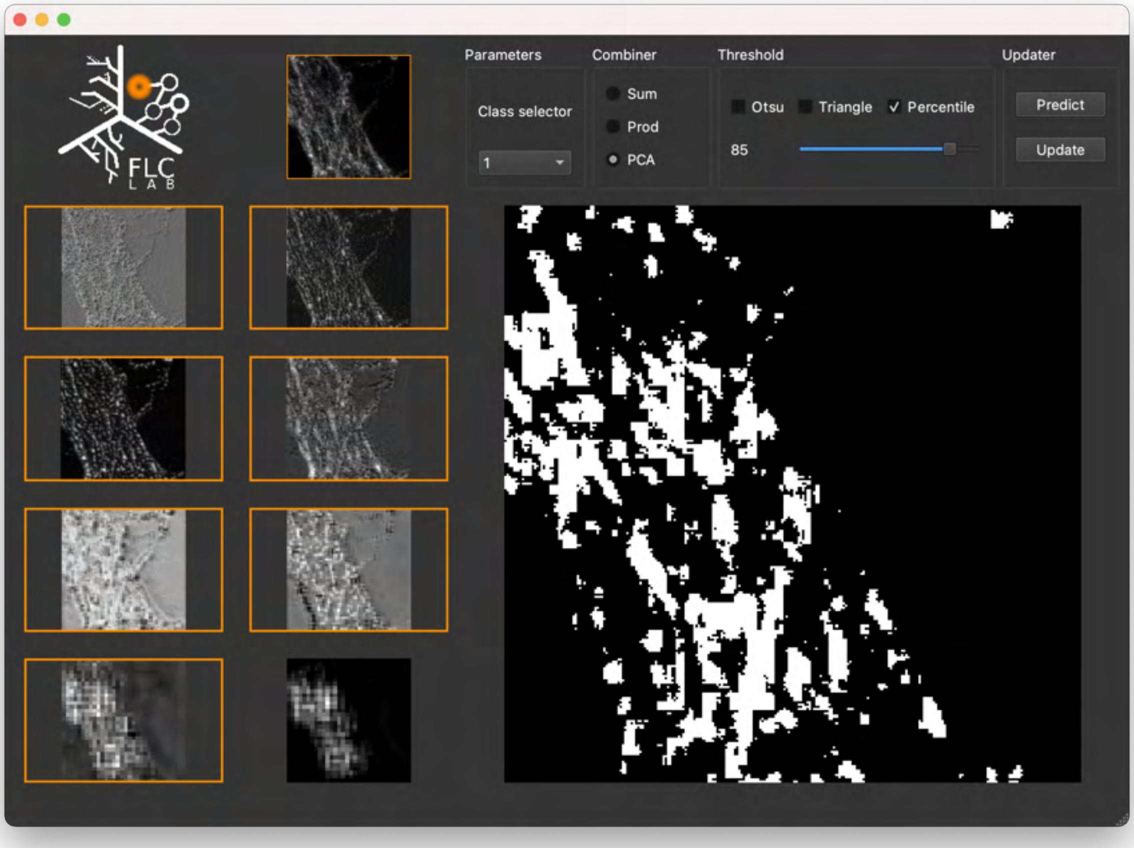


Supplementary Figure 30: Graphical user interface (GUI) developed to facilitate the visualisation of the extracted local maps from MICRA-Net. The detailed steps to use the application can be found on the GitHub repository (`https://github.com/FLClab/MICRA-Net`). Briefly, one can load a trained MICRA-Net model and an image to predict the presence of a specific structure. The GUI shows the extracted local maps $L^{1-8}$ to the user for each activated class of the selected image. The user can select the desired local maps which are combined into a detailed feature map that can be thresholded to generate a final segmentation mask.

### Supplementary Note 1: Modified MNIST Dataset

| Parameters | Values |
|---|---|
| Epochs | 150 |
| Batch size | 64 |
| Objective function | Binary cross entropy with logits |
| Learning rate | 0.01 |
| Learning rate scheduler | N/A |
| Minimal learning rate | N/A |
| Data augmentation | N/A |

Supplementary Table 22: Training parameters of MICRA-Net for the modified MNIST dataset.

#### 1.1 MICRA-Net segmentation and evaluation

The procedure detailed in section 4.1.1 was followed to obtain the coarse segmentation. For each detected digits, a minimal certainty of 30% (qualitatively chosen from the validation set) was imposed to lower the rate of false-positive detection during segmentation and the `argmax` projection was used over the 10 classes to avoid overlap between detections. The feature maps was extracted from the network using the procedure explained in section 4.1.1 (Figure 2a-c). Precise binary segmentation maps were generated by applying a local Otsu threshold [4] for each $8 \times 8$ patch from the extracted coarse segmented region, which resulted in more accurate segmentation maps (Figure 2d,e & Supplementary Fig. 5& 6). A bounding box of 28 pixels centered on each digit was used to compute the metrics, allowing a more balanced ratio between foreground and background pixels. The evaluated metrics are shown in Figure 2e & Supplementary Fig.5& 6.

#### 1.2 Baseline architecture

For each network (fully-supervised and $\{5, 10, 25\}$ dilation dataset), the U-Net architecture was trained from scratch, keeping all hyper-parameters constant in all instances. U-Net architecture: Each step in the contracting path consists of two sets of $3 \times 3$ convolutional layers, followed by a batch normalization, and a $2 \times 2$ max-pooling. The number of filters in each layers was doubled after each contraction and are of size $\{16, 32, 64, 128\}$. The expanding path was symmetrical to the contracting path, but a $2 \times 2$ transposed convolution (stride of 2) was used to increase the layer size. Skip links are used to propagate information from higher layers. A final $1 \times 1$ convolutional layer was used to output the 11 classes segmentation map. The maximal argument along the class axis of the output was used as the semantic segmentation of the input image. ReLU activation was used throughout the network.

The U-Net was trained from scratch using the ADAM optimizer with default values and with a learning rate 0.001 for 150 epochs. Binary cross entropy with logits was used as the loss to minimize. The model generalizing the most on the validation dataset was kept for testing. The U-Net was trained to output segmentation of 11 classes (all digits and background) resulting in a semantic segmentation of the images in the modified MNIST dataset. Since this architecture requires ground truth segmentation, the digits were used as binary masks to train U-Net in a fully-supervised manner (Figure 2f & Supplementary Fig. 7). This fully-supervised training consists in a high-standard baseline since U-Net has access to information that is not available in a weakly-supervised setting. For a more realistic baseline, the U-Net was also trained with weak annotations. To this end, the binary ground truth contours from the training dataset were dilated with an increasing size of the square structuring element ($\{5, 10, 25\}$ pixel), simulating an annotator contouring the images with coarse annotations, while still evaluating against undisturbed images (Figure 2f & Supplementary Fig. 7).

| Parameters | Values |
|---|---|
| Epochs | 250 |
| Batch size | 32 |
| Objective function | Binary cross entropy with logits with weights |
| Learning rate | 0.001 |
| Learning rate scheduler | - Reduce factor : 10 |
| | - Validation reduction : <0.01 |
| | - Patience : 10 epochs |
| Minimal learning rate | $1 \times 10^{-5}$ |
| Data augmentation | - Horizontal flip |
| | - Vertical flip |
| | - Intensity scale |
| | - Gamma adaptation |

Supplementary Table 23: Training parameters of MICRA-Net for the F-Actin dataset.

### Supplementary Note 2: F-actin Dataset

#### 2.1 MICRA-Net training procedure

The dataset is highly imbalanced towards negative crops. To account for data imbalance, a weighted binary cross entropy with logits was used:

$$l(\hat{y}, y) = L = \{l_1, ..., l_N\}^\top, l_n = p_n y_n \cdot \log \sigma(\hat{y}_n) + (1 - y_n) \cdot \log \sigma(1 - \hat{y}_n), \quad (1)$$

where $p_n$ is the positive weight associated with class $n$. The positive weights during training were set to 3.3 and 1.6 for the periodical lattice and fibers respectively.

#### 2.2 MICRA-Net segmentation and evaluation

The extracted feature map was globally thresholded to the $80^{\text{th}}$ percentile of the intensity of the image following a Gaussian blur with $\sigma = 1$. These parameters were selected based on a qualitative evaluation on the validation dataset. Data imbalance (between the number of foreground and background pixels) was accounted for by calculating the metrics inside the dendritic mask therefore reducing the number of negative pixels

#### 2.3 Baseline architectures

Three different baselines were trained to predict the F-actin nanostructures using weak supervision : U-Net, Mask R-CNN and Ilastik. The following sections describe the specific implementation and training details of each models.

##### 2.3.1 U-Net

U-Net was trained using polygonal bounding boxes as annotations on the same images provided to MICRA-Net for training. The architecture is the same as in Lavoie-Cardinal et al. [5]. It is trained to output two independent segmentation maps, *i.e.* periodical lattice and fibers.

The baseline was trained as in Lavoie-Cardinal et al. [5]. The proper thresholds to generate binary segmentation maps for both structures were extracted from a receiver operating characteristic (ROC) curve on the validation dataset. No post-processing steps were applied on the generated masks.

##### 2.3.2 Mask R-CNN

Mask R-CNN was trained using weak supervision from polygonal bounding boxes similarly to the U-Net implementation. The model from the PyTorch library (not pre-trained) with slight modifications was used [6,

7]. The RGB input layer was replaced by a single channel layer. The number of classes to predict was set at 3 (periodical lattice, fibers and background). The normalization values were calculated from the normalized training images (`image-mean` : 0.0110, `image-std` : 0.0212).

The Adam learning optimizer was used with an initial learning rate of $1 \times 10^{-4}$ and default parameters. The model with the best generalization properties on the validation set was kept for testing (calculated from the objective loss function). Only the crops containing a positive detections were kept for training. The network was trained for 700 epochs with a batch size of 32. The size of the input crops were set at $256 \times 256$ pixel. The same data augmentation technique were used for Mask R-CNN as in MICRA-Net training.

At inference, a segmentation threshold of 0.5 was used on the predicted masks and a non maximum suppression threshold of 0.7 between predicted bounding boxes was used as in the seminal implementation [7]. No post-processing steps were applied on the generated masks.

###### 2.3.3 Ilastik

Ilastik was trained with weak scribbles annotations simulated from the polygonal bounding boxes. The positive scribbles were generated by skeletonizing the polygonal bounding boxes of both structures (rings and fibers). Negative scribbles were simulated from the skeleton of the dendritic mask with the polygonal bounding boxes of both structures removed. A number of background pixels (outside the dendritic mask) corresponding to the sum of both positive class were sampled. The default parameters of Ilastik were used for training.

At inference, Ilastik was used in `headless` mode to predict all pixels from the testing images. No post-processing steps were applied on the generated masks.

### Supplementary Note 3: Cell Tracking Challenge Dataset

| Parameters | Values |
|---|---|
| Epochs | 700 |
| Batch size | 48 |
| Objective function | Binary cross entropy with logits |
| Learning rate | $1 \times 10^{-4}$ |
| Learning rate scheduler | - Reduce factor : 2 |
| | - Validation reduction : <0.01 |
| | - Patience : 100 epochs |
| Minimal learning rate | $1 \times 10^{-5}$ |
| Data augmentation | - Horizontal flip |
| | - Vertical flip |
| | - Intensity scale |
| | - Gamma adaptation |

Supplementary Table 24: Training parameters of MICRA-Net on the Cell Tracking Challenge dataset.

#### 3.1 MICRA-Net training procedure

The number of generated crops from each cell line can greatly vary which could introduce imbalance bias between cell lines. On the other hand, the number of annotated images is more balanced than the number of generated crops from each cell lines. Hence, at the beginning of each epoch a single crop was randomly sampled from each annotated image in the dataset in order to create a subset of crops that is balanced in terms of cell lines.

#### 3.2 MICRA-Net segmentation and instance segmentation

MICRA-Net was trained on two different version of the dataset: i) 256 × 256 pixel, and ii) 128 × 128 pixel. A dense prediction was computed on the testing images using crops with a 50% overlap. To avoid undesired edge effects, each testing images was padded using a symmetric padding and a 10 pixel border was removed from the predicted crops. A Gaussian blur was applied on the extracted feature map and a Otsu threshold [4] generated the segmentation mask. For the instance segmentation task, the contact between cell was subtracted from the detailed cell feature map. The segmentation was obtained by a Otsu threshold of the resultant feature map.

#### 3.3 Baseline architectures

Three different baseline architectures were trained on the Cell Tracking Challenge: pre-trained U-Net, Mask R-CNN and Ilastik. All baseline architectures were trained in both fully- and weakly-supervised manner. For both level of supervision, the baseline model and training procedure are the same.

##### 3.3.1 Pre-trained U-Net

The pre-trained fully-supervised U-Net provided by Falk et al. [3] was used. This implementation of U-Net was pre-trained on the Cell Tracking Challenge dataset and other in-house datasets in an instance segmentation task (differentiate between the cells and the background). The Caffe model was first converted in PyTorch. The original architecture [3] was used with small modifications. A 0-padding of the convolutional steps is used in order to output a segmentation map that has the same shape as the input image. To maintain similar classification task between MICRA-Net and U-Net, 6 output channels are required (5 cell lines and background).

For training, the Adam learning optimizer with a learning rate of $1 \times 10^{-4}$ and default parameters is used. The model with the best generalization properties on the validation dataset was kept for testing. A weighted cross-entropy loss was used as in Falk et al. [3] to help with the separation of cells. Both the weights between adjacent cells and the weights on cells compared to background were increased. The network was trained for 700 epochs with a batch size of 16. The size of the input crops were set at 256 × 256 pixel and the pixel step was set at 192 pixels. Similarly to MICRA-Net, a single crop per image was sampled from the annotated images. The same data augmentation procedure as MICRA-Net was used.

At inference 256 × 256 pixel crops were extracted with a 50% overlap. Overlapping crops were averaged and the `argmax` function generated the semantic segmentation.

##### 3.3.2 Mask R-CNN

The implemented model from the PyTorch library (not pre-trained) with slight modifications [6, 7] was used. The input layer was modified to take a single channel image as input instead of the common RGB image. The number of classes to predict was set at 6 (5 cell lines and background). The normalization values were calculated from the normalized training images (`image-mean` : 0.2695, `image-std` : 0.0664).

Adam learning optimizer was used with a learning rate of $1 \times 10^{-4}$ and default parameters. The model with the best generalization properties on the validation dataset was kept for testing. Only the crops which contained cells were kept for training. The network was trained for 700 epochs with a batch size of 16. The size of the input crops were set at 256 × 256 pixel and the pixel step was set at 192 pixels. The same data augmentation technique as in MICRA-Net were used for Mask R-CNN. A single crop per annotated image was sampled at each epoch of training.

Similarly to pre-trained U-Net, 256 × 256 pixel crops were extracted with a 50% overlap. The objects detected in all overlapping crops were kept. As in the seminal implementation, a segmentation threshold of 0.5 and a non maximum suppression threshold of 0.7 between predicted bounding boxes [7] were used.

##### 3.3.3 Ilastik

The Graphical User Interface (GUI) with the default parameters provided by the Ilastik software [8] were used. At training, the user is required to annotate images at the pixel level. Ilastik requires to load the complete training dataset into the memory of the computer. Unfortunately, the complete Cell Tracking Challenge dataset could not be loaded all at once on the computer. Hence, a subset of positive and negative pixels were sampled to generate a dataset which could fit into memory (same number of positive and negative pixels).

At inference, Ilastik was used in `headless` mode to predict all pixels from the testing images.

### Supplementary Note 4: P. Vivax Dataset

| Parameters | Values |
|---|---|
| Epochs | 1200 |
| Batch size | 64 |
| Objective function | Binary cross entropy with logits |
| Learning rate | $1 \times 10^{-4}$ |
| Learning rate scheduler | - Reduce factor : 2 |
| | - Validation reduction : <0.005 |
| | - Patience : 50 epochs |
| Minimal learning rate | $1 \times 10^{-5}$ |
| Data augmentation | - Horizontal flip |
| | - Vertical flip |
| | - Random rotations (0-360°) |
| | - Intensity scale |
| | - Shearing |

Supplementary Table 25: Training parameters of MICRA-Net on the P. Vivax dataset.

#### 4.1 MICRA-Net training procedure

The dataset was highly unbalanced towards uninfected red blood cells. Hence, the training procedure was adapted to balance the number of positive (infected red blood cells) and negative (background or uninfected cells) crops. Instead of training with all crops from an image at each epoch, 5 to 8 crops were randomly sampled per images. The sampled crops were then randomly assigned to a mini-batch. The number of positive and negative crops was balanced by sampling an uninfected cell (or background) with a probability of 10%.

#### 4.2 MICRA-Net fine-tuning procedure

To fine-tune MICRA-Net, $\{12, 24, 36\}$ images were randomly selected from the testing dataset. It is important to note that the 12 images were a subset of the 24 images, and the 24 images a subset of the 36 images (see Supplementary Fig 25). Several approaches were tested to fine-tune MICRA-Net and their results were compared on the resultant testing set (84 images). The approaches included i) adjusting the detection threshold [*Threshold*], fine-tuning the ii) linear layer [*Linear*] iii) linear layer and depth 4 [*Linear + 4*] iv) linear layer and depths 3 and 4 [*Linear + 3, 4*], and v) all [*All*] layers. To adjust the detection threshold, the 5 trained models were used and the detection threshold were adjusted on the $\{12, 24, 36\}$ sampled images. For fine-tuning, a 3-fold training procedure was used to generate a small validation set for early stopping and to reduce the probability of over-fitting of the model (see Supplementary Fig. 25). Adam was used as the optimizer with a learning rate of $1 \times 10^{-6}$. The models were trained for 100 epochs. The state of the model generalizing the most on the validation set for each fold were kept for testing. The objective function used was binary cross entropy with logits. The learning rate of the models were not reduced in contrast to the original training phase. At inference, an ensemble model of the 3 trained models from each fold was used and their predictions were averaged. The 3-fold training was repeated 5 times from each of the 5 naive models as base model, generating a total of 25 ensemble models (see Supplementary Fig. 25). The same procedure for the detection and segmentation task as described above were applied.

##### 4.3 MICRA-Net detection

At inference, all $256 \times 256$ pixel crops with a pixel step of 32 were extracted from the testing image and the probability of presence of an infected red blood cell was predicted. The prediction of overlapping crops were averaged and all maxima using the `peak_local_max` function from the Scikit-Image Python library [9] were located. From these predicted positions and their associated probability, a precision-recall curve was generated to optimize the detection level of a given model on the validation set. A maximal distance of 128 pixels was used to associate detected and ground truth objects.

##### 4.4 MICRA-Net segmentation

A Gaussian blur with a sigma parameter of 3 was applied and the resultant feature map was thresholded at the $80^{\text{th}}$ percentile. Small post-processing operations on the segmentation maps were used. Small holes ($< 50 \times 50$ pixel) were removed from the generated binary masks and the most prominent object was kept in cases where multiple infected cells were present in the extracted crops. This was necessary to evaluate the performance only on the subset of cells that were precisely annotated to generate the testing set. To do so, the object which contained the maximal intensity from the extracted feature map of MICRA-Net was selected. This procedure was qualitatively evaluated on the validation set.

#### Supplementary Note 5: Scanning Electron Microscopy Dataset

| Parameters | Values |
|---|---|
| Epochs | 600 |
| Batch size | 16 |
| Learning rate | $1 \times 10^{-4}$ |
| Objective function | Binary cross entropy with logits |
| Learning rate scheduler | - Reduce factor : 2<br>- Epochs : 50, 200, 300 |
| Minimal learning rate | $1 \times 10^{-5}$ |
| Data augmentation | - Horizontal flip<br>- Vertical flip<br>- Random rotations (0-360°)<br>- Intensity scale<br>- Gamma adaptation<br>- Shearing<br>- Elastic transform<br>- Random position |

Supplementary Table 26: Training parameters of MICRA-Net on the Scanning Electron Microscopy dataset.

##### 5.1 MICRA-Net training procedure for SEM segmentation

Different Positive-Unlabeled (PU) ratios [10] were compared for training $\{2:1, 1:1, 1:2, 1:5, 1:8, 1:16\}$. Negative crops were randomly assigned according to the PU ratio. The negative crops of a specified PU ratio are a subset of the next higher PU ratio. Considering that the extracted crops are larger in size ($1024 \times 1024$ pixel) than the training size ($512 \times 512$ pixel), random position sampling could be used in training. This method increased the effective number of crops and served as another data augmentation technique.

##### 5.2 MICRA-Net Detection

All crops from a testing image were extracted using a sliding window of size $512 \times 512$ pixel with a 128 pixels step. The probability of presence of a Axon DAB marker was predicted using MICRA-Net. The overlapping crops were averaged and all maxima were located using the `peak_local_max` function from the Scikit-Image Python library [9]. Using these predicted positions and their associated probability, a precision-recall curve was generated to optimize the detection level of a given model on the validation set. A maximal distance of 512 pixels was used to associate detected and ground truth objects.

##### 5.3 MICRA-Net Segmentation

A Gaussian blur with a sigma parameter of 5 was applied and the resultant feature map was thresholded at the $90^{\text{th}}$ percentile. Small holes ($< 45 \times 45$ pixel) were removed from the generated binary masks and only the most prominent object was kept for the same reason explained for the P. Vivax dataset. The object which contained the maximal intensity from the generated feature map of MICRA-Net was selected. This procedure was qualitatively evaluated on the validation set.

##### 5.4 Baseline architectures

The performance of MICRA-Net at the detection of Axon DAB was compared with Ilastik using a weakly-supervised training procedure.

###### 5.4.1 Ilastik

The Graphical User Interface (GUI) of Ilastik with the default parameters provided by the Ilastik software [8] was used. Simulated weak positive annotations from an Expert were used by creating a circle of fixed radius (15 pixels) which was approximately the size of an Axon DAB marker. Ten times more negative pixels were sampled than positive pixels from the input images to account for the high diversity of structure within the electron microscopy images. This allowed a good compromise between diversity of pixels and class imbalance.

At inference, Ilastik was used in `headless` mode to predict all pixels within the scanning electron microscopy testing images.

#### References


[1] Vladimír Ulman, Martin Maška, Klas EG Magnusson, Olaf Ronneberger, Carsten Haubold, Nathalie Harder, Pavel Matula, Petr Matula, David Svoboda, Miroslav Radojevic, et al. An objective comparison of cell-tracking algorithms. *Nature methods*, 14(12):1141–1152, 2017.

[2] Juan C. Caicedo, Jonathan Roth, Allen Goodman, Tim Becker, Kyle W. Karhohs, Matthieu Broisin, Csaba Molnar, Claire McQuin, Shantanu Singh, Fabian J. Theis, and Anne E. Carpenter. Evaluation of Deep Learning Strategies for Nucleus Segmentation in Fluorescence Images. *Cytometry Part A*, 95 (9):952–965, 2019. ISSN 1552-4930. doi: 10.1002/cyto.a.23863.

[3] Thorsten Falk, Dominic Mai, Robert Bensch, Özgün Çiçek, Ahmed Abdulkadir, Yassine Marrakchi, Anton Böhm, Jan Deubner, Zoe Jäckel, Katharina Seiwald, et al. U-net: deep learning for cell counting, detection, and morphometry. *Nature Methods*, 16(1):67, 2019.

[4] Nobuyuki Otsu. A threshold selection method from gray-level histograms. *IEEE Transactions on Systems, Man, and Cybernetics*, 9(1):62–66, 1979.

[5] Flavie Lavoie-Cardinal, Anthony Bilodeau, Mado Lemieux, Marc-André Gardner, Theresa Wiesner, Gabrielle Laramée, Christian Gagné, and Paul De Koninck. Neuronal activity remodels the f-actin based submembrane lattice in dendrites but not axons of hippocampal neurons. *Scientific Reports (Nature Publisher Group)*, 10(1), 2020.

[6] Adam Paszke, Sam Gross, Soumith Chintala, Gregory Chanan, Edward Yang, Zachary DeVito, Zeming Lin, Alban Desmaison, Luca Antiga, and Adam Lerer. Automatic differentiation in pytorch. In *31st Conference on Neural Information Processing Systems*, 2017.

[7] Kaiming He, Georgia Gkioxari, Piotr Dollár, and Ross Girshick. Mask R-CNN. *arXiv:1703.06870 [cs]*, January 2018.

[8] Stuart Berg, Dominik Kutra, Thorben Kroeger, Christoph N. Straehle, Bernhard X. Kausler, Carsten Haubold, Martin Schiegg, Janez Ales, Thorsten Beier, Markus Rudy, Kemal Eren, Jaime I. Cervantes, Buote Xu, Fynn Beuttenmueller, Adrian Wolny, Chong Zhang, Ullrich Koethe, Fred A. Hamprecht, and Anna Kreshuk. ilastik: interactive machine learning for (bio)image analysis. *Nature Methods*, September 2019. ISSN 1548-7105. doi: 10.1038/s41592-019-0582-9. URL `https://doi.org/10.1038/s41592-019-0582-9`.

[9] Stefan Van der Walt, Johannes L Schönberger, Juan Nunez-Iglesias, François Boulogne, Joshua D Warner, Neil Yager, Emmanuelle Gouillart, and Tony Yu. scikit-image: image processing in python. *PeerJ*, 2:e453, 2014.

[10] Jessa Bekker and Jesse Davis. Learning from positive and unlabeled data: a survey. *Mach. Learn.*, 109 (4):719–760, 2020.
